# Gastruloid patterning reflects division of labor among biased stem cell clones

**DOI:** 10.1101/2025.07.12.664536

**Authors:** Vinay Ayyappan, Catherine G Triandafillou, Kavitha Sarma, Arjun Raj

**Affiliations:** Department of Bioengineering, School of Engineering and Applied Sciences, University of Pennsylvania, Philadelphia, PA, USA; The Wistar Institute, Gene Expression and Regulation Program, Philadelphia, PA, USA; Department of Cell and Developmental Biology, Perelman School of Medicine, University of Pennsylvania, Philadelphia, PA, USA; Department of Genetics, Perelman School of Medicine, University of Pennsylvania, Philadelphia, PA, USA

## Abstract

Division of labor, whereby individuals specialize in complementary roles to collectively achieve beneficial outcomes, is a recurring phenomenon in economics, ecology, and microbiology. In development, individual cells specialize, but this specialization is thought to arise from instruction by external signaling cues upon otherwise interchangeable progenitors. It is unclear if progenitors exhibit some degree of specialization, and if so, whether that specialization is optimized for developmental outcomes. Using fluorescence-based lineage tracing in combination with spatial transcriptomics, we show that, in the gastruloid model of early development, individual stem cell clones harbored reproducible spatial propensities for anterior or posterior fates that result in a spontaneous division of labor during axial organization. Gastruloids derived from pure clones generated elongated structures less frequently than a polyclonal population, but mixing clones allowed clones to follow their inherent propensity, restoring proper axial elongation. Spatial transcriptomics reveals that pure clones show disrupted gene expression with inappropriate coexpression of anterior and posterior markers, while clone combinations restore proper spatial organization. We further show that propensities are globally utilized: a clone with a particular propensity can adopt different fates depending on what is more optimal for development as a whole. Perturbations to key developmental signaling pathways disrupted this sorting and profiling revealed molecular characteristics of propensity. Proper developmental outcomes may thus emerge from the coordinated action of intrinsically biased clonal populations.

## Introduction

Division of labor, in which distinct cells cooperate by specializing in complementary roles, is an organizing principle that recurs across biological systems at every scale. Social insects coordinate through specialized castes^1^, microbial communities distribute metabolic functions among subpopulations^2^, and even cancer cell populations exhibit cooperative specialization to proliferate and invade^2,3^. Division of labor is made possible because of differences in the participating cells, which occupy roles for which they are best suited. Can such a principle extend to cells of the early embryo? It is well established that heterogeneity emerges even at the earliest stages of embryonic development^4^. While differences among embryonic stem cells can influence their differentiation outcomes^4–6^, it remains unknown whether these differences are deleterious or beneficial to overall development.

What if heterogeneity seeds a division of labor among stem cells? Pluripotent stem cells are capable of forming all tissues, but individual stem cells might still have a propensity (preference) to form a particular tissue. If such biases existed, it is possible that developing systems could preferentially assign cells to roles that they are best suited to fill, even if all cells were competent to adopt any role. This behavior is analogous to the notion of “comparative advantage” in economics, whereby individuals can specialize according to their relative strengths. Cells with stronger biases for particular fates could have a comparative advantage in forming the corresponding tissues, and therefore could specialize in forming those tissues. A prediction of this model is that a developing system that channels cells toward their preferred fates could produce more consistent developmental outcomes than one that assigns fates indiscriminately. Testing whether concepts like division of labor and comparative advantage apply to developing systems in which genetically identical cells differ only in their epigenetic states remains untested, in part because it requires a system where the clonal composition of a population of cells can be precisely controlled while also allowing individual lineages to be traced through development.

A precedent for division of labor in development comes from blastocyst complementation experiments. In these experiments, a blastocyst is made of cells with a mutation that prevents them from differentiating into a particular tissue. Exogenous pluripotent stem cells that are competent for forming all tissues are injected into this blastocyst^7^. These added cells rescue development of the missing tissue. Remarkably, they contribute almost exclusively to the tissue the host cannot make, rather than contributing proportionally across all lineages, as though they recognize which developmental niche is vacant and fill precisely that role. This division of labor has been demonstrated in eye^8,9^, pancreas^8^, and lung development^10^. These examples demonstrate that developmental systems can exploit differences among cells to allocate complementary roles. However, in blastocyst complementation, the division of labor arises between genetically distinct populations. Whether such division of labor emerges among genetically identical cells has not been tested.

The gastruloid is a model of development that is well-suited for testing whether cells undergoing early morphogenetic changes display a division of labor^11,12^. Gastruloids begin as 3D aggregates of embryonic stem cells (ES cells) that break symmetry in response to a uniformly administered Wnt agonist and elongate. Even in the absence of maternal cues or predefined patterning centers, gastruloids self-organize to spontaneously produce cell types derived from all three germ layers, organized along the anterior-posterior (A-P) body axis. Unlike the embryo, gastruloids develop from a starting population of defined size and composition that can be systematically manipulated, making it possible to directly test whether pre-existing differences in the propensities of individual cells for filling specific developmental roles gives rise to a division of labor that shapes both the fates those cells adopt and overall gastruloid morphogenesis.

Standard gastruloid protocols treat ES cells as functionally equivalent, with morphogenetic outcomes determined by aggregate cell number and signaling conditions rather than the specific composition of the starting population^13,14^. Here, cells are thought to acquire distinct identities through morphogen gradients and signaling centers, with the pre-existing differences among naive stem cells less relevant to the final outcome^15–20^. However, multiple studies^21–23^ have revealed that ES cells frequently fluctuate in state even under carefully controlled culture conditions, with expression of pluripotency genes and lineage markers changing over time^24,25^. Recent work has begun to characterize the developmental consequences of this variation: monoclonal gastruloids exhibit heritable lineage biases^5^, differential Wnt signaling activity shapes individual cell contributions to axial patterning^5,26^, and wild-type cells can rescue defects in chimeric aggregates with mutant cells^27^. But cells differing in their outcomes does not tell us whether this variation matters for the system as a whole. Pre-existing heterogeneity could be noise that gastruloid development tolerates or averages over, or it could be that gastruloid self-organization actively exploits cell-cell variability, with cells dividing labor as each cell takes on specific developmental roles. Distinguishing these possibilities requires testing whether development proceeds differently when that variation is absent.

Here, we show that clonal diversity among mouse ES (mES) cells facilitates consistent gastruloid morphogenesis. Using a multicolor fluorescence-based lineage-tracing system, spatial transcriptomics, and chromatin accessibility profiling, we show that individual clones consistently favor specific positional fates along the A-P axis. When combined, biased clones complement each other’s limitations, allowing gastruloids to robustly polarize and elongate. These findings reveal that clonal diversity within a stem cell population is not merely tolerated during gastruloid morphogenesis but is organized into a division of labor, where biased clones cooperate by specializing in complementary developmental roles. Clones’ biases, which emerge from subtle differences in developmental signaling pathways, enable them to partition the task of gastruloid development and achieve proper spatial patterning in gastruloids. The principle that morphogenetic precision emerges from the coordinated contributions of intrinsically different cells may represent a general feature of self-organizing developmental systems.

## Results

### Clonal diversity promotes proper gastruloid organization

Gastruloids are typically generated from polyclonal mouse embryonic stem cell (mES cell) aggregates. These cells, while often assumed to be homogeneous, can have highly variable states, even under carefully controlled culture conditions^5,28–32^. Gastruloids, therefore, provide a rich system for mapping the potentially variable characteristics of individual mES cells to their fate outcomes in a self-organizing cellular community. As per the standard mouse gastruloid protocol, we seeded ∼300 mES cells in non-adherent conditions (t=0) and applied a timed pulse of Wnt agonist (Chiron, CHIR99021) at 48 hours post-aggregation (t=48). Chiron was removed after 24 hours (t=72), and gastruloids were grown to 120 hours, at which time they should have elongated along a single anterior-posterior (A-P) axis **(Figure 1A)**^26,33–36^.

**Figure 1.**
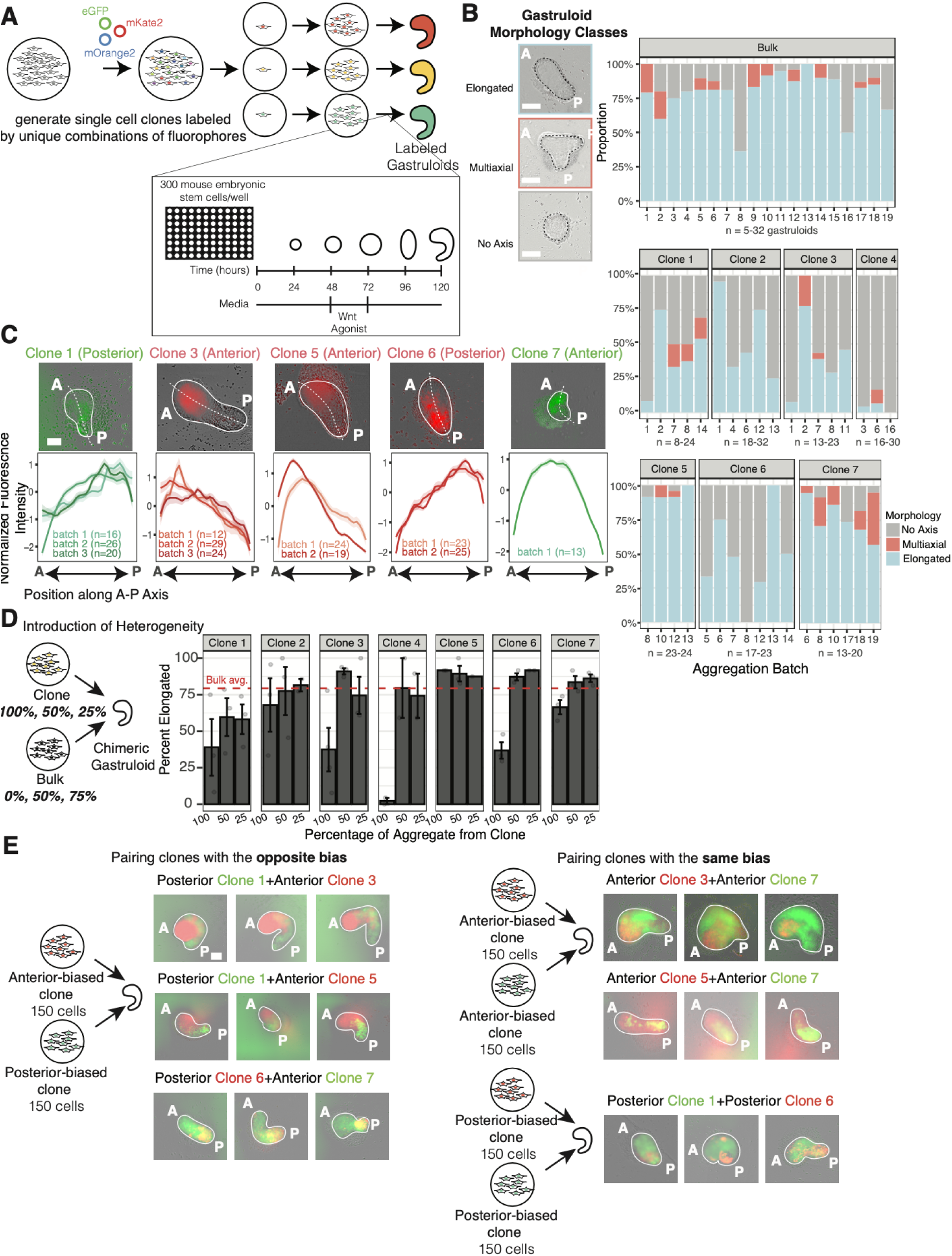
mES cell clone diversity promotes gastruloid self-organization. **(A)**. Schematic describing gastruloid formation and our fluorescent lineage tracing scheme. mES cells expressing unique combinations of the fluorophores eGFP, mKate2, and mOrange2 are dilution cloned and expanded into clonal mES cell lines, which are then used to generate gastruloids. **(B)** Gastruloid morphologies are classified as “elongated,” “multiaxial,” or “no axis”. Bargraphs represent the proportion of gastruloids within each batch of aggregation classified as belonging to a given class of morphologies (n=8-33 gastruloids per batch). The scale bar represents 100px. Annotations (A) and (P) label the gastruloid anterior and posterior, respectively. Each experiment was assigned an “aggregation batch number.” Experiments presented in this panel included pure clone gastruloids and bulk gastruloids; gastruloids aggregated on the same day share a batch number to enable comparison of pure clone morphologies to bulk gastruloids from the same day/plate. **(C)** Representative Incucyte images of clone-bulk combinations are shown, alongside line graphs showing the normalized (as described in **Methods**) fluorescence intensity along the gastruloid A-P axis. Bold lines show the average intensity profile for gastruloids within a given batch, and error ribbons represent one standard deviation. The scale bar represents 100px. **(D)** 300-cell chimeric aggregates where 100%, 50%, or 25% of the cells belong to a given clone and the remainder of the cells belong to the bulk were generated. Bargraphs showing the average fraction of gastruloids that are “elongated” for each chimeric combination are shown. A dotted red line depicts the average fraction of gastruloids that are “elongated” across aggregation batches. **(E)** Three representative images for each combination of clones. Shown are pairings of anterior-biased clones with posterior-biased clones, anterior-biased clones with anterior-biased clones, and posterior-biased clones with posterior-biased clones. The scale bar represents 100px.

Do clonal mES cell populations form gastruloids more or less efficiently than a polyclonal bulk population? We generated single-cell-derived clones of mES cells, leading to a population that is more homogeneous than the original bulk population. After isolating individual mESCs by dilution cloning (**Figure 1A**), we compared gastruloids generated from seven individual clonal lines to those from bulk populations, classifying 120-hour gastruloid morphologies into three categories: "elongated" (single A-P axis), "multiaxial" (multiple axes or projections), and "no axis." Each clone could form gastruloids, indicating that they could differentiate into anterior and posterior cell types. Gastruloids generated from bulk mES cells formed elongated gastruloids approximately 75% of the time, in keeping with literature^35^. Gastruloids generated from pure mES cell clones, however, formed elongated structures less frequently, rather forming multiaxial gastruloids or failing to develop a clear axis (3-17 batches, n=8-32 gastruloids per batch; **Figure 1B**). Most clones were thus less efficient than the heterogeneous bulk population at producing properly organized structures, suggesting these cell populations were more sensitive to initial conditions and thus less robust at gastruloid formation. Successful morphogenesis may therefore depend not on the properties of any individual clone, but on the collective contributions of a diverse starting population.

### Individual mES cell clones have reproducible spatial biases despite retaining broad differentiation potential

We observed that heterogeneous bulk populations of cells form gastruloids more efficiently than pure clones. One oft-cited possibility is that of lineage priming, in which some cells may, at a particular point in time, be predisposed or “primed” for a particular fate and thus enable symmetry-breaking. An alternative possibility is that individual cells have propensities that propagate over multiple cell divisions—i.e., they have a persistent memory of propensity^37,38^. In the latter scenario, individual clones may play distinct, complementary roles during morphogenesis, a division of labor in which each clone preferentially contributes to fates it is biased toward. To distinguish between these possibilities, we needed to track the location and fate of individual clones. We implemented a fluorescence-based lineage-tracing strategy whereby mES cells express varying combinations of three fluorophores (mKate2, mOrange2, eGFP) that localize exclusively to the nucleus (**Figure 1A**). We could thereby visually observe how they organize within gastruloids as they self-organize.

In generating gastruloids from a polyclonal, multicolor population, we observed that cells from the same clone tended to localize to similar regions in the gastruloid (**Supplemental Figure 1**, **Supplemental Movie 1**). Perhaps each clone is biased, such that its cells’ fates are skewed to a particular outcome. We generated chimeric aggregates, where 50% (∼150 cells) of the starting aggregate was obtained from an unlabeled, bulk mES cell population and the remaining 50% of cells belonged to a single labeled clone. If the clones were unbiased, we might expect labeled cells to be either scattered throughout the gastruloid or to appear in clumps as labeled cells proliferate during gastruloid development. Instead, we noticed that labeled cells in chimeric gastruloids separated from unlabeled bulk ES cells (**Supplemental Movie 2**); that is, the clone seemed to have different sorting properties from the remaining population.

Was spatial separation of labeled clones from the unlabeled bulk mES cells the result of differential adhesion between clone cells and bulk cells or an inherent tendency to form specific tissues in the gastruloid? If differential adhesion were the explanation, we might expect that, across aggregates, fluorescently labeled clones would segregate from their unlabeled counterparts, but for any given clone, labeled cells in some gastruloids would appear in the gastruloid posterior and those in other gastruloids might occupy the anterior. However, we noticed that in the resulting chimeric gastruloids, cells from the labeled clone consistently tended to appear toward either the gastruloid anterior (3 clones) or posterior (2 clones). Each clone had a specific propensity for anterior or posterior domains (Clones 2 and 4 were excluded for technical reasons, see **Methods**), with nearly every gastruloid across several batches of aggregation showing the same pattern for a given labeled clone when combined with unlabeled bulk mES cells (**Figure 1C**, **Supplemental Figure 2A-C**).

We also wanted to confirm that each clone could still form tissues derived from all three germ layers, and that propensities did not necessarily reflect a general loss of differentiation potential. We subjected each clone to differentiation protocols intended to produce endoderm, mesoderm, or neural cell fates (**Supplemental Figure 3**) and immunostaining for germ layer markers T (mesoderm), Sox17 (endoderm), and Pax6 (neural/ectoderm). Under standard pluripotency conditions, neither clones nor bulk mES cells expressed markers of the germ layers. However, upon directed differentiation, all clones successfully expressed appropriate markers: Sox17 (endoderm), T (mesoderm), and Pax6 (neural/ectoderm). Therefore, clones seem to retain broad differentiation potential.

While we were able to demonstrate that several individual clones have spatial propensities, we wondered whether most or all individual mES cell clones had this same property. To do so, we made chimeric gastruloids comprising unlabeled cells from a bulk mES cell population and a small number of cells from a polyclonal GFP-labeled mES cell population, corresponding to 1-3 clones (**Supplemental Figure 4A-C**). Looking across a large number of gastruloids, we could determine whether the GFP-labeled clones were located on either end of gastruloids, indicating a propensity; or if they were evenly distributed throughout. Of the 129 elongated gastruloids, 59 included GFP-labeled cells in the gastruloid anterior, and 43 included GFP-labeled cells in the posterior, with the remaining 27 gastruloids including GFP+ cells throughout the anterior-posterior axis (**Supplemental Figure 4D**). The fact that most gastruloids showed GFP-labeled clones at either end of the A-P axis suggests that many clones do harbor a spatial propensity for occupying the gastruloid anterior or posterior. Taken together, these results suggest that spatial propensities are widespread among mES cell clones rather than a property of rare outliers, and argue against the possibility that propensities are somehow introduced through the process of dilution cloning and single cell isolation.

### Division of labor promotes proper gastruloid organization

In economics, division of labor maximizes overall productivity because each individual performs the task it is best suited for. We wondered whether an analogous principle applies in gastruloids: when a clone is forced to perform all developmental roles on its own, it may do poorly at those roles for which it has no propensity, but in a mixed population, each clone could preferentially contribute to the fates it is biased toward, potentially benefiting the gastruloid as a whole. To test this idea, we mixed cells from individual clones with bulk cells and assessed the frequency of proper elongation. When we mixed cells from individual clones with bulk cells, the resulting gastruloids had elongated morphologies much more frequently than gastruloids formed from the pure clone alone. We found that as the percentage of bulk cells in the starting aggregate increased, the proportion of elongated gastruloids increased as well (**Figure 1D**). Even in pure clones (e.g., clone 4) that, alone, rarely formed elongated gastruloids, 50%/50% clone-bulk aggregates formed elongated gastruloids as frequently as bulk mES cell aggregates. To check whether clones that less efficiently formed elongated gastruloids were not being eliminated (e.g., due to competition) from chimeric aggregates, we confirmed that both clones were present in clone-clone and clone-bulk combinations by detecting fluorescence corresponding to cells from either clone (**Supplemental Figure 2F**). Even in clone 4, which rarely formed elongated gastruloids on its own, we saw fluorescent cells from clone 4 in gastruloids where clone 4 comprised 25% of the cells in the initial aggregate, indicating that cells from clone 4 were not completely eliminated during gastruloid elongation. Therefore, introducing heterogeneity to aggregates by combining clones with bulk mES cells improved gastruloids’ ability to form elongated structures.

Gastruloids generated from combinations of clones also tended to more consistently form properly elongated structures than pure clones (statistically-significantly so, **Supplemental Figure 2E, Supplemental Figure 5**). Most clone combinations resulted in elongated gastruloids, regardless of whether clones in the chimera were both anterior, both posterior, or if one clone was anterior and the other was posterior, and in all cases one clone predominantly formed the gastruloid anterior and the other formed the gastruloid posterior. One exception was the combination of clone 4 (likely posterior, see **Methods, Supplemental Figure 2F**) and clone 6 (posterior), which only infrequently (∼20%) resulted in elongated gastruloids. On its own, clone 4 formed elongated gastruloids <10% of the time, while clone 6 formed gastruloids that elongated ∼50% of the time (**Supplemental Figure 4**). A more heterogeneous mES cell pool in chimeric aggregates thus generally compensates for pure clones’ inefficiencies in producing elongated gastruloids, though there are still some clone combinations that do not improve gastruloid morphology over their respective pure clones.

Taken together, these results suggest that although individual mES cell clones might harbor specific spatial propensities and constraints in their ability to generate gastruloids, heterogeneous aggregates can be more capable of properly forming an elongated gastruloid.

### Propensities reflect comparative advantage among clones

There are two explanations for the observation that individual clones had a strong bias towards occupying a particular niche in the gastruloid. One is that those cells are always directed to a particular region based on their propensity. However, it is also possible that cells are directed to a particular region based on their *comparative* propensity; that is, it is the relative propensity that dictates which clone’s cells go where, with the latter requiring cells to have considerably more flexibility in what fates they adopt, as is the case in the blastocyst complementation experiments alluded to earlier. Such a comparative mechanism is in many ways analogous to the concept of comparative advantage in economics, in which two parties may both have an advantage for producing, say, hats instead of shoes, but the optimal division of labor is one in which the one with the greater advantage for producing hats produces hats and the other produces shoes^39^. Our system allowed us to ask whether individual clones within a gastruloid might exhibit a similar division of labor based on comparative advantage by asking what happens when we mix clones with seemingly the same propensity relative to bulk: if we mixed two clones with an anterior propensity together and formed a gastruloid, would the clones intermix, or would one adopt an anterior fate and the other a posterior fate based on their comparative advantage? The latter scenario would indicate that cells’ propensities enable them to collectively coordinate their fates to optimally pattern gastruloids.

Indeed, when clones with opposite propensities were aggregated together, the anterior clone occupied the gastruloid anterior and the posterior clone occupied the posterior, consistent with each clone specializing in the role for which it is best suited (**Figure 1E**, **Supplemental Figure 2D**). Importantly, however, pairing two anterior clones still resulted in relative sorting between them, though they appeared more mixed than pairings of clones with opposite propensity. Propensity therefore varies continuously among clones, such that even clones with the same spatial bias can exhibit relative differences among them.

To more directly test whether clones are capable of forming anterior and posterior tissues in gastruloids, we performed immunofluorescence staining for the posterior marker T and manually annotated gastruloid outlines and T-expressing regions. Gastruloids generated from pure clones 1 and 6, which each have posterior propensities, had a larger T-positive region on average (∼40% of the gastruloid area) than bulk gastruloids (∼20% of gastruloids’ area) and those made from clones 3, 5, and 7, which have anterior propensity. However, each clone could generate gastruloids containing cells expressing T as well as cells that do not express T (**Supplemental Figure 6A**, **B**). Therefore, each clone, despite exhibiting spatial specialization, is broadly capable of establishing both anterior and posterior identities. This distinction between what a clone preferentially does and what it can do further parallels the concept of comparative advantage, as clones specialize not because they necessarily lack the capacity for other fates, but because they are relatively better suited to particular ones.

Patterning in gastruloids made from bulk mES cell populations has been shown in the literature to be consistent^33,40^. If clones divide labor according to their comparative advantages, how does the system respond when one clone is over- or underrepresented? We generated aggregates comprising clone 1 (posterior) and clone 3 (anterior) in ratios ranging from 1:4 to 4:1 and measured each clone’s spatial distribution along the A-P axis. Regardless of starting composition, the boundary between clone territories fell at a consistent position (∼35% of the distance) along the A-P axis (**Supplemental Figure 7**, **8A**). Clones that started at high abundance grew proportionally less; underrepresented clones compensated by growing more. There was batch-to-batch variation in where the clone boundary lay; however, the boundary was not sensitive to the ratio of clone 1 to clone 3 in the initial aggregate. Even co-culturing clones before aggregation did not alter this outcome (**Supplemental Figure 8B**, **C**). This constant final proportionality is inconsistent with passive sorting and instead demonstrates active regulation of clones’ proliferation. The division of labor among clones thus does not require precise starting ratios. Rather, the gastruloid compensates for an imbalance in the starting composition by preferentially expanding whichever clone is underrepresented relative to the role it needs to fill. Thus gastruloids converge on a consistent spatial organization even when the balance of comparative advantages in the starting population is skewed.

### Removing clonal diversity disrupts spatial patterning of marker gene expression

What happens when cells are forced to act against their natural propensity? In a pure-clone gastruloid, every developmental role must be filled by the same clone, including roles for which it has no inherent propensity. Since gastruloids formed from pure clones so often fail to elongate or properly polarize, one possibility is that spatial propensities of an individual clone disrupt the establishment of positional identities, especially in regions where cells are forced to act against their propensity. Such disruption may manifest as inappropriate expression or coexpression of anterior and posterior markers in the posterior or anterior region, respectively. We used spatial transcriptomics to directly examine spatial gene expression patterns in pure-clone versus bulk gastruloids. We used a panel of 21 genes, comprising marker genes for cell types expected to be in the gastruloid at 120 hours, to perform sequential single-molecule RNA FISH (seqFISH)^41^ (**Figure 2A**). We tested gastruloids from a bulk mES cell population (n=2), posterior clone 1 (n=5), anterior clone 3 (n=5), and posterior clone 6 (n=2).

**Figure 2.**
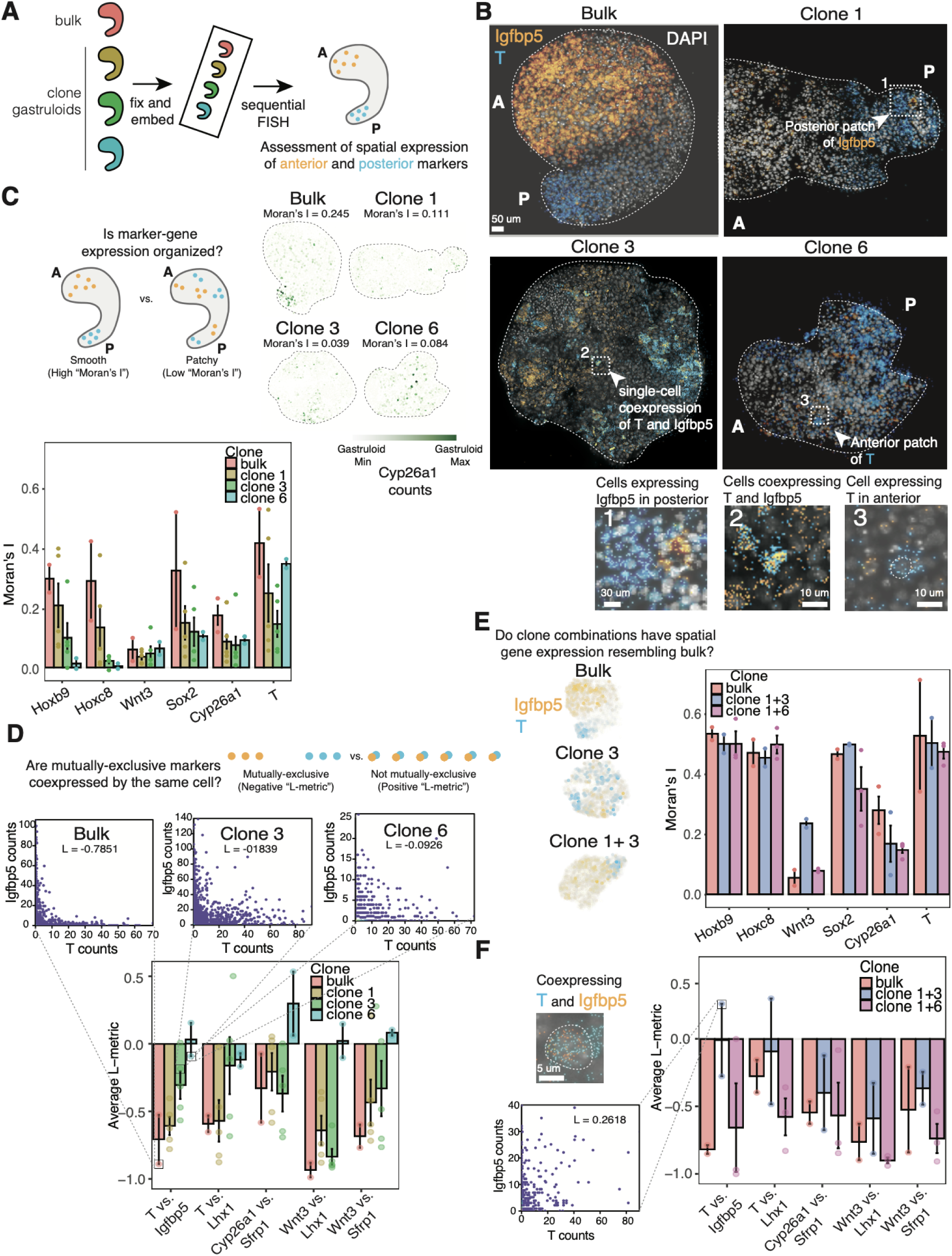
Spatial transcriptomics reveals patterns of gene expression underlying morphologies of pure-clone gastruloids. **(A)** Schematic describing the experimental workflow. seqFISH was performed on gastruloids made from bulk mES cells and from labeled mES cell clones. **(B)** Representative images showing the expression of *T* (blue) and *Igfbp5* (gold) in gastruloids made from bulk mES cells or mES cell clones 1, 3, and 6. Insets highlight examples where cells are coexpressing *T* and *Igfbp5*, or highlight posterior *Igfbp5* expression or anterior *T* expression. **(C)** Quantification of Moran’s I across bulk and pure-clone gastruloids (bulk: n=2, posterior clone 1: n=5, anterior clone 3: n=5, posterior clone 6: n=2) within a batch. Representative gastruloids, where cells are colored on the basis of their *Cyp26a1*, are shown above. **(D)** Quantification of the L-metric across bulk and pure-clone gastruloids within the same batch. Representative hexbin plots are shown, comparing the expression of *T* and *Igfbp5* in selected gastruloids. **(E)** Quantification of Moran’s I across bulk and chimeric gastruloids from a second seqFISH batch (bulk: n = 2, clone 1+3: n = 2, clone 1+6: n = 3). **(F)** Quantification of L-metric across bulk and chimeric gastruloids from this second seqFISH batch. A representative hexbin plot of *T* and *Igfbp5* expression from a selected clone 1+3 gastruloid is shown, alongside an image of a cell from this gastruloid coexpressing *T* and *Igfbp5*.

To test whether pure-clone gastruloids fail to properly form and organize cells into anterior and posterior identities, we first examined the spatial relationship between T, a marker of the gastruloid posterior, and Igfbp5, an anterior marker. In well-organized gastruloids, we expect these markers to be spatially segregated and mutually exclusive, reflecting proper patterning along the anterior-posterior axis. Gastruloids generated from a bulk mES cell population had polarized expression of *T*, which was mutually exclusive with expression of *Igfbp5*. However, gastruloids generated from a pure clone, which were less capable of forming elongated structures, had visible patches of *T* and *Igfbp5* that were often intermixed, with some cells coexpressing the two genes (**Figure 2B**, **Supplemental Figure 9A**). Of note, bulk gastruloids had strong expression of *Igfbp5* and *T* in the right locations. Clone 3 gastruloids (anterior propensity) had expression of *Igfbp5* and *T* similar to bulk (**Supplemental Figure 9D**). In contrast, clone 6 gastruloids (posterior propensity) had low overall expression of *Igfbp5* but still had strong expression of posterior markers like *T* and *Cyp26a1* (another posterior marker), consistent with its propensity.

To quantify differences in the spatial organization of marker gene expression, we first used the Moran’s I statistic to determine whether gene expression patterns are clustered (cells with similar gene expression are near one another, corresponding to a positive Moran’s I), spatially dispersed (i.e., cells with high gene expression must be surrounded by cells with low gene expression and vice versa, corresponding to a negative Moran’s I), or random (zero). In general, Moran’s I values for genes in our panel were higher for the bulk gastruloids than for gastruloids generated from a pure clone, indicating more spatially clustered gene expression in bulk gastruloids (**Figure 2C**, **Supplemental Figure 9B**). Even in an example of a bulk gastruloid that was not properly elongated, markers like *Igfbp5*, *T*, *Hoxb9*, and *Sox2* were appropriately confined, with relatively high Moran’s I values (**Supplemental Figure 10**). Our results were consistent with pure-clone gastruloids doing a poorer job of polarizing marker gene expression.

Next, we wanted to determine whether pure-clone gastruloids coexpressed anterior and posterior marker genes. In general, we expected the expression of these marker genes to be mutually exclusive. A cell with high expression of anterior markers would likely have low expression of posterior markers, and a cell with high expression of posterior markers would have low expression of anterior markers, resulting in an “L-shaped” scatter between the two genes. We devised a metric, which we term the “L-metric,” to quantify the degree to which gene expression pairs are mutually exclusive (**Methods**). Negative values indicate mutual exclusivity, positive values indicate coexpression. For example, we expect *T* and *Igfbp5* to have a negative L-metric value, as might *Wnt3* with the Wnt inhibitor *Sfrp1*. Indeed, for these gene pairs that we expect to be mutually exclusive, bulk gastruloids consistently had negative L-metric values (**Figure 2D**, **Supplemental Figure 9C**). Pure-clone gastruloids, on the other hand, had less strongly negative L-metric values. Of the clones tested, clone 1 showed L-metric values most similar to bulk, while clone 6 (posterior propensity) showed positive L-metric values, on average, for *T* and *Igfbp5*, *Cyp26a1,* and *Sfrp1*, and *Wnt3* and *Sfrp1* (**Supplemental Figure 9**, see **Methods** for statistical comparison of gene mutual exclusivity). Thus, cells from clones with anterior propensity showed more "mixed" expression patterns when forming cells with posterior identities, and cells from clones with posterior propensity showed mixed expression patterns when forming anterior tissues.

### Restoring clonal diversity improves spatial gene expression patterning

Since clone combinations more consistently produced elongated gastruloids, we tested whether chimeric gastruloids showed the coherent spatial gene expression seen in bulk gastruloids. We performed seqFISH on gastruloids generated from a combination of one anterior clone (clone 3) and one posterior clone (clone 1) (n=2) and from a combination of two posterior clones (clone 1 and clone 6) (n=3). Chimeric gastruloids formed from clone pairs now showed more similar Moran’s I values to bulk (n=2) for most genes in the panel (**Figure 2E**, **Supplemental Figure 9B**). There were some exceptions; for example, the Moran’s I value for the anterior marker *Igfbp5* was much lower than bulk in gastruloids generated from the posterior-posterior combination of clone 1 and clone 6. These gastruloids had robustly polarized *T* expression, but only sparse expression of *Igfbp5*. Thus, combining distinct mES cell clones at least partially restores the spatial gene expression coherence we observed in bulk gastruloids, in contrast to the disrupted patterns seen above in pure-clone gastruloids.

Next, we wondered if these chimeric gastruloids now had less coexpression of marker genes that were normally mutually exclusive. We again quantified these differences using the L-metric as a measure of mutual exclusivity. For marker gene pairs that showed significant coexpression in pure clones (high L-metric; e.g., *Cyp26a1* vs. *Sfrp1*), we found that the L-metric values in the chimeric gastruloids were far lower, indeed, showing no significant differences from bulk (**Figure 2E**). On average across chimeric gastruloids, we observed negative L-metric values for marker gene pairs like *T* and *Igfbp5*, *Cyp26a1* and *Sfrp1*, and *Wnt3* and *Sfrp1.* However, there was substantial variability among gastruloids, with one gastruloid generated from clone 1 and clone 3 showing many cells that coexpressed *T* and *Igfbp5* (**Supplemental Figure 9C**), and the coexpressing cells tended to belong to clone 3 (**Supplemental Figure 11**). Our analysis of marker gene coexpression thus confirmed our above Moran’s I findings through a different measure of spatial gene expression coherence: combining clones at least partially restores more organized gene expression patterns compared to pure-clone gastruloids, consistent with their improved morphogenetic outcomes.

### Clones partition cell type production and retain signatures of their original bias

A key prediction of a model in which division of labor is organized by comparative advantage is that when a cell fills a role outside its specialty, it should still retain signatures of its original bias. Gastruloids contain multiple cell types in complex patterns that extend beyond simple anterior-posterior organization, and the division of labor among clones might reflect biases for particular developmental lineages beyond a purely spatial preference for forming the gastruloid’s anterior or posterior. We therefore distinguished between two related but distinct concepts: "fate bias" refers to the intrinsic cellular tendency toward specific lineage outcomes, while "spatial propensity" describes the resulting positional preferences that clones exhibit during gastruloid self-organization. If division of labor operates at the level of lineage biases rather than spatial domains, then clones might adopt fates outside their usual territory when their partners cannot fully supply those cell types. We asked whether clones showed this kind of flexibility, and if so, whether they retained transcriptional evidence of their original bias.

We looked at localization in chimeric gastruloids generated from clone 1 (posterior propensity) and clone 3 (overall anterior propensity). While cells from clone 3 were typically anteriorly localized and cells from clone 1 were posteriorly localized, some clone 3 cells could be found in the posterior tip of the gastruloid (**Supplemental Figure 12A**, **C**). When we immunostained gastruloids from different clone combinations (**Supplemental Figure 12C-E**, **Supplemental Figure 13**), we found that in many clone pairings, like clone 1+3 and clone 6+7, the anterior clone occupied the anterior region of the gastruloid, and also had cells residing in the *T*-positive posterior tip. Division of labor may thus extend beyond anterior-posterior segregation, with clones’ biases potentially extending to cell types in addition to broad spatial territories.

We used spatial transcriptomics to compare cells from posterior clone 1 and from anterior clone 3 within the posterior of chimeric gastruloids formed from these clones. Cells from clone 3 coexpressed anterior marker *Igfbp5* with posterior marker *T*, retaining a transcriptional signature of their anterior origin even while occupying a posterior role. Conversely, clone 1 cells, despite their posterior spatial propensity, actually expressed relatively low levels of *T* compared to clone 3 cells within the gastruloid posterior, where *T* should be expressed most strongly (**Supplemental Figure 12A**, **B**). However, clone 1 still had comparatively higher expression of *Thy1*, a marker for posterior neuroectoderm as well as some mesenchymal cell types^42–44^, confirming its general bias for the posterior. In this particular pairing, clone 3 had a stronger bias for *T*-expressing cell types than clone 1, even though clone 3 typically occupies anterior regions. It is possible that it is thus able to “fill in” for the missing *T*-positive cells based on its propensity. These results are consistent with a division of labor organized by comparative advantage. Clones can contribute outside their usual domain when needed, but retain some transcriptional signatures of their original propensities when doing so.

### Propensity erodes through rounds of passaging

If propensity is a stable, heritable property of the clone rather than a transient fluctuation, it should persist through moderate expansion but might eventually erode as cells divide and their molecular states shift over time. We wondered whether clones’ propensities were stable over further rounds of cell division beyond the expansion following isolation (**Supplemental Figure 14**). We generated chimeric gastruloids from pairs of mES cell clones and from pairings of clones with bulk mES cells at multiple passages and found that, though clones had clear propensities at passage 1 post-thaw, aggregates generated at passage 3 were well-mixed. Beyond passage 3, chimeric aggregates, rather than having well-separated clones, frequently developed into gastruloids that featured a mix of labeled clones throughout (**Figure 3A**, **Supplemental Figure 15A-D**). Thus, while clonal propensity was robust to large-scale expansion, it was not preserved through serial passaging after thawing.

**Figure 3.**
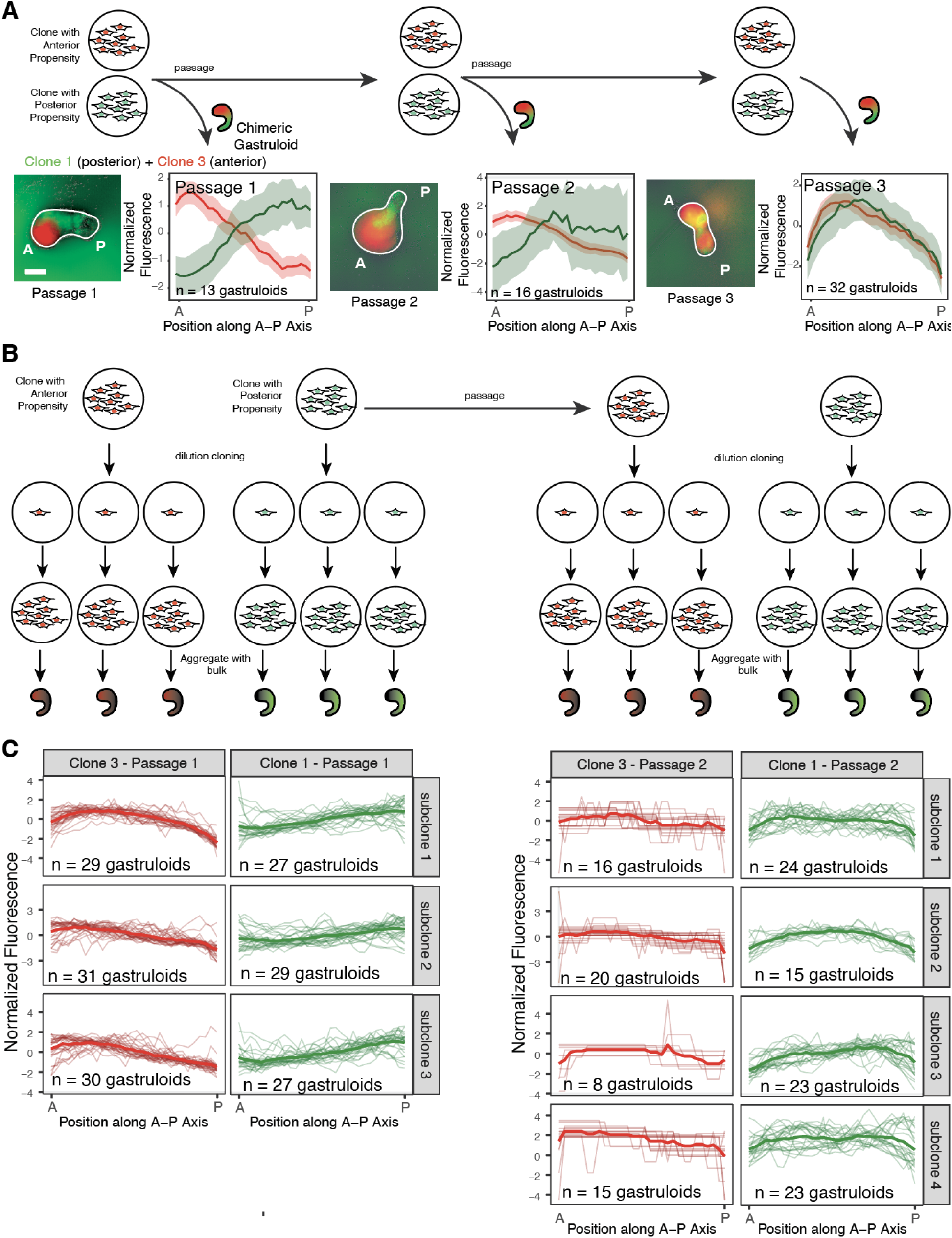
Propensity decreases across passages. **(A)** Representative images and quantification of changes in clones’ propensity across three rounds of passaging. Gastruloids from the combination of clone 1 and clone 3 are shown. Scale bars represent 100px. **(B)** Schematic describing isolation of subclones from mES cell clones. **(C)** Individual fluorescence intensity curves for every gastruloid generated from each subclone isolated from clone 1 and clone 3 at passage 1 and at passage 2. Normalization is described in **Methods**.

We performed replicate experiments with several parental clones across multiple passages. In most cases, clones lost their characteristic propensity by passage 4. However, in some replicate experiments, we observed variability; for example, clone 3 maintained its propensity at passage 4 in a third replicate (**Supplemental Figure 15C**-**D**). These results suggested that propensity is not stably maintained through extended passaging and that loss of propensity may be partially influenced by stochastic or context-dependent factors.

We considered two possibilities for how clones lose spatial propensity during passaging. One possibility is that a small subset of unbiased cells lacking any propensity pre-exist in the population and selectively expand during passaging, overtaking the culture. A second possibility is that individual cells change their propensity during passaging. To distinguish between these scenarios, we performed an additional round of bottlenecking to derive subclones from clonal lines **(Figure 3B)**. We then generated chimeric gastruloids by aggregating each subclone with bulk mES cells. If truly unbiased cells existed within the population, then those unbiased subclones isolated from the parental clone population would mix evenly with bulk mES cells in chimeric gastruloids. Subclones isolated from passage 1 cells of anterior clone 3 and posterior clone 1 retained the spatial propensity of their parent line **(Figure 3C),** and none of the subclones lacked propensity. These results are consistent with the loss of propensity not being due to expansion of an unbiased subpopulation that was already present, although we cannot rule out the possibility that some rare unbiased clones exist in the population. We next passaged the parental clones once and then isolated subclones. Differently then passage 1, subclones isolated from the passage 2 parental clones lacked clear spatial propensities **(Figure 3C)**. This result shows that individual cells change their propensity, reverting to a state with seemingly less propensity. Together, these results suggest that passaging leads to a loss or “forgetting” of the original propensity across the population.Moreover, these results suggest against the possibility that the process of bottlenecking the bulk mES cell population itself somehow prompts the establishment of propensities.

Some post-passage-2 subclones generated gastruloids mostly with a uniform distribution of labeled cells along the A-P axis. Since these subclones tended to uniformly arrange themselves along the gastruloid A-P axis, they likely “lost” their propensity during passaging. Other subclones seemed to feature a mix of propensities, generating some gastruloids with labeled cells predominantly in the anterior and other gastruloids with labeled cells predominantly in the posterior (**Supplemental Figure 16**); some cells within these subclones therefore had anterior propensity and others had posterior propensity. These results suggest that after passaging and over the course of subsequent rounds of cell division, cells within each subclone can acquire anterior or posterior propensities, which may resemble the emergence of diverse propensities within a bulk population.

### Transcriptional differences among mES cell clones do not reflect differences in pluripotency or anterior-posterior patterning programs

We wondered what molecular differences, if any, underlie clonal propensities. We have shown that clones exhibit reproducible spatial biases while retaining broad developmental competence. To ask whether these biases correspond to transcriptional differences between clones, we performed bulk RNA-seq on our clonal mES cell lines (3 anterior clones, 2 posterior clones).

We identified 230 differentially expressed genes between anterior and posterior clones out of 14,883 total genes detected (**Figure 4B**). The genes that were differentially expressed showed diverse functions, including cell adhesion (*St8sia6, Nphs1*), chemokine signaling (*Cxcl14*), and growth factor regulation (*Igfbp7*), but did not correspond to coherent developmental pathways or well-known anterior-posterior patterning programs (**Supplemental Figure 17**). There were no significant differences in the expression of pluripotency markers (e.g., *Oct4, Sox2, Klf4, Nanog*) (**Figure 4A-C**), nor were there significant differences in expression of markers for cell types present in mature gastruloids (**Figure 4D**), nor in the expression of cadherins (**Supplemental Figure 17A**). The subtle transcriptional differences identified likely reflect heterogeneity in cellular states rather than commitment to specific lineages.

**Figure 4.**
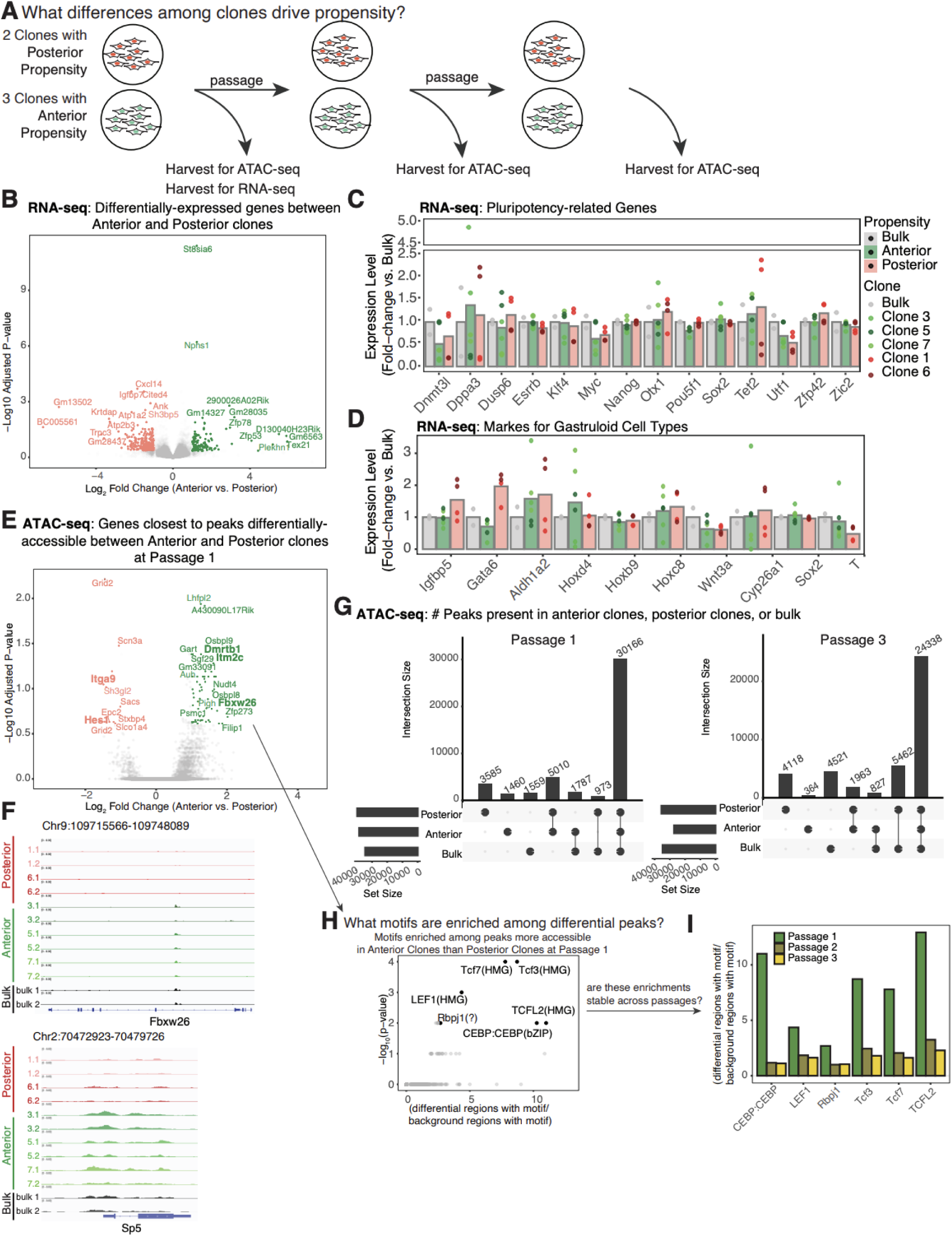
ATAC-seq and RNA-seq of mES cell clones show differences in chromatin accessibility associated with propensity. **(A)** Schematic describing the workflow for harvesting samples for RNA-seq and ATAC-seq. **(B)** Volcano plot showing differentially-expressed genes between anterior clones and posterior clones. **(C)** Bargraph comparing the expression of pluripotency markers among bulk mES cells, anterior clones, and posterior clones. **(D)** Bargraph comparing the expression of marker genes for cell types expected to be present in gastruloids. **(E)** Volcano plot showing genes near differentially-accessible genomic regions between anterior clones and posterior clones. **(F)** Representative ATAC-seq tracks for Fbxw26 and Sp5, genes with genomic regions more accessible in anterior clones. **(G)** Upset plots showing the number of peaks unique to posterior clones, anterior clones, or bulk mESCs or peaks shared among these samples. Plots are shown for samples harvested at passage 1 and at passage 3. **(H)** Scatterplots showing motifs identified by HOMER as being more enriched among peaks more accessible in anterior clones than posterior clones at passage 1. The x-axis shows enrichment scores: the fold-change between the proportion of sequences containing each motif among differentially accessible peaks versus the proportion of sequences containing each motif among peaks randomly selected through the mouse genome. The y-axis shows adjusted p-values. Labeled peaks have enrichment scores > 2.5 and adjusted p-values < 0.05. **(I)** Barplots comparing motif enrichment scores across passages, for each motif annotated as significant in **(H)**.

### Propensities are consistent with changes in chromatin accessibility at developmentally relevant motifs

Since propensity did not seem to be associated with coherent expression differences between clones, we wondered if differences in chromatin accessibility were more strongly associated with propensity. Our results showing that propensity diminished over successive passages indicated a form of epigenetic memory that, while not permanent, is stable over many rounds of cell division. We hypothesized that clones with propensities for anterior or posterior fates had epigenetic differences that make drivers of anterior or posterior fates more likely to be expressed. We therefore sought to determine what chromatin accessibility differences exist between mES cell clones differing in propensity. ATAC (assay for transposase-accessible chromatin) sequencing is a way to measure differences in chromatin accessibility. We harvested chromatin from bulk mES cells and mES cells from each clonal line across three rounds of passaging to perform ATAC-seq (**Figure 4A**).

We first focused on differences among clones at passage 1. Were there differences in chromatin accessibility that might indicate differences in priming for anterior or posterior fates among clones (**Figure 4E-G)**? Across all samples, we identified 44,540 peaks. The majority of peaks (67.7%) were shared among all clones and bulk mES cells, likely reflecting their common pluripotent identity. However, the remaining peaks we observed were specific to anterior and posterior clones: 3.22% of peaks were unique to anterior clones, while 8.05% were unique to posterior clones. In addition, 3.50% of peaks were unique to bulk, 2.18% were shared between only bulk and posterior clones, 4.01% were shared only between bulk and anterior clones, and 11.24% were shared only between anterior and posterior clones (**Figure 4G**).

In comparing differentially-accessible peaks between clones with anterior and posterior propensities, we observed significant differences in some genomic regions near genes with known functions in development (**Figure 4E-G**). For example, posterior clones showed increased accessibility in peaks near *Hes1*, which has a known role in coordinating the timing of differentiation among mES cells in a population^45,46^ and in skewing differentiation toward mesodermal trajectories^46^. We did not find accessibility differences in genomic regions associated with cadherin genes (**Supplemental Figure 17B**).

Next, we also considered later passages, by which time clones had generally lost their visible propensities. We looked at the number of peaks present in anterior clones, posterior clones, or bulk samples. Across all samples, we identified 41,593 peaks, of which 58.5% of peaks were shared among anterior clones, posterior clones, and bulk. 9% of peaks were unique to posterior clones, while <1% of peaks were unique to anterior clones (**Figure 4G**, **Supplemental Figure 19A**, **B**).

What transcription factors might be related to accessibility differences that we observe? At each passage, we used HOMER to identify transcription factor DNA binding motifs enriched among regions differentially-accessible between anterior clones and posterior clones (**Figure 4H**, **Supplemental Figure 19E-G**). At passage 1, we found six motifs significantly enriched in anterior clones compared to posterior clones, including CEBP (a retinoic acid signaling target^47^), Rbpj (a Nodal signaling regulator and Notch target involved in ectoderm-derived tissue differentiation^48,49^), Tcf3 (which inhibits Wnt signaling to promote anterior identity^50,51^), and three Wnt pathway components (TCFL2, Tcf7, LEF1) known to regulate anterior-posterior axis specification^26,52^.

The enrichment of CEBP, Rbpj, and Tcf3 in anterior clones is consistent with increased accessibility at regulatory sites for pathways that promote anterior identity–CEBP for retinoic acid-mediated anteriorization, Rbpj for Notch-driven ectoderm specification, and Tcf3 for Wnt pathway repression. The enrichment of Wnt pathway transcription factors in sites of increased chromatin accessibility in anterior clones was initially surprising, since these factors typically promote posterior fates. However, we reasoned that since mES cells were cultured in the presence of Chiron (a Wnt agonist that maintains active Wnt signaling in all cells), anterior clones might have enhanced accessibility at binding sites for Wnt targets involved in negative feedback regulation rather than pathway activation. Consistent with this hypothesis, we observed increased accessibility near Wnt inhibitory genes like *Sp5* and *Fbxw26* in anterior clones^53^(**Figure 4E**, **F, Supplemental Figure 19C-D**). RNA-seq analysis revealed 1.57-fold higher expression of the Wnt inhibitor *Axin2* in anterior versus posterior clones (**Supplemental Figure 18C**). These results suggest that anterior clones are primed to more readily activate Wnt pathway inhibitors even under uniform Wnt stimulation, which could explain their propensity for anterior regions during gastruloid development.

To further validate these findings, we performed transcription factor footprinting analysis using TOBIAS, which detects evidence of transcription factor occupancy from ATAC-seq data rather than just the presence of motif sequences. Footprinting indicated differential binding at Wnt pathway components (Tcf7, Tcf7l1, Lef1) and additionally revealed differential binding at the Nodal target Mixl1, Notch targets Hes1 and Hey2, Fox family members, and Hox and Pax family members (**Supplemental Figure 20**).

While anterior clones showed a reduction in unique peaks, the overall patterns remained similar between passages, suggesting that gross chromatin accessibility differences are maintained even as propensities are lost. The preservation of broad accessibility patterns raises the possibility that the regulatory changes that drive the loss of propensity are specific to particular regulatory pathways. To test this possibility, we checked whether differences in motif enrichment between anterior and posterior clones persisted across passages. We looked at differentially accessible peaks (|log2FC| > 1) between anterior and posterior clones at passage 2 and passage 3. For motifs enriched in anterior clones at passage 1, we asked what percentage of regions more accessible in anterior clones contained that motif and computed an enrichment score by comparing this percentage to the percentage of randomly selected sequences in the mouse genome that contain that motif using HOMER. Enrichment for all motifs that were enriched in anterior clones at passage 1 decreased in passages 2 and 3 (**Figure 4I)**. The loss of motif enrichment over passages suggests that accessibility differences at developmentally relevant transcription factor binding sites, like Wnt, Nodal, and retinoic acid, are associated with clonal propensities.

Transcription factor footprinting analysis across passages supported this conclusion. Comparing footprints at passage 1 versus passage 3, we found that the majority of inferred differentially-bound motifs were passage-specific rather than shared (**Supplemental Figure 20B**). For posterior clones, only 10 of 113 differentially-bound motifs were shared between P1 and P3; for anterior clones, 61 of 144 were shared. Critically, the motifs lost by P3 included the developmentally relevant factors identified at passage 1. *Tcf7*, *Lef1*, and multiple Fox family members were differentially bound at passage 1 but not passage 3. Clones seem to remain epigenetically distinct at later passages. Even though, if anything, the total number of differentially accessible peaks increases, propensity is lost nonetheless. It is thus possible that the nature of the epigenetic differences changes over 3rounds of passaging. At passage 1, clones differ at regulatory sites for the pathways that pattern the anterior-posterior axis. However, by passage 3, clones still differ, but at loci that no longer predict spatial behavior.

### Perturbing developmental signaling disrupts propensity

Morphogens, like Wnt, retinoic acid (RA), and Nodal, are thought to shape gastruloid self-organization by forming gradients along which anteroposterior identity is established^54–56^. A cell’s position within a gradient thus influences its own fate choice and its ability to modulate the signaling microenvironment for itself and its neighbors. If clones’ propensity is shaped by differential responses to morphogen gradients, then perturbing morphogen pathways might distort their arrangement within the gastruloid.

Our ATAC-seq experiments revealed RA and Nodal signaling as plausible pathways associated with propensity. Both pathways are thought to affect the organization of cells in the embryo, in part by modulating their response to Wnt signaling^55–57^. To test whether chromatin-accessibility differences between anterior and posterior clones result in transcriptional differences when clones are challenged to differentiate during gastruloid development, we performed bulk RNA-seq on gastruloids collected 48 hours after aggregation. We compared gastruloids derived from clonal mES cells with posterior propensity, and two clonal lines with anterior propensity (one strongly anterior, and one with a weaker anterior propensity, **Supplementary Figure 21A**), as well as the polyclonal bulk population. Among differentially expressed genes, we observed clone-specific variation in Nodal and RA pathway components (**Supplementary Figure 21**), indicating that differences in Nodal and RA signaling inferred from chromatin accessibility in mES cells manifest transcriptionally during gastruloid morphogenesis.

We hypothesized that perturbing RA or Nodal signaling should disrupt the characteristic spatial arrangement of clones within gastruloids, either by affecting their ability to segregate from one another or their ability to consistently occupy the gastruloid’s anterior or posterior. As above, we defined “propensity” as the tendency of a clone to occupy a specific axial domain, and “sorting” as the spatial segregation of different clones within the gastruloid (**Figure 5A**).

**Figure 5.**
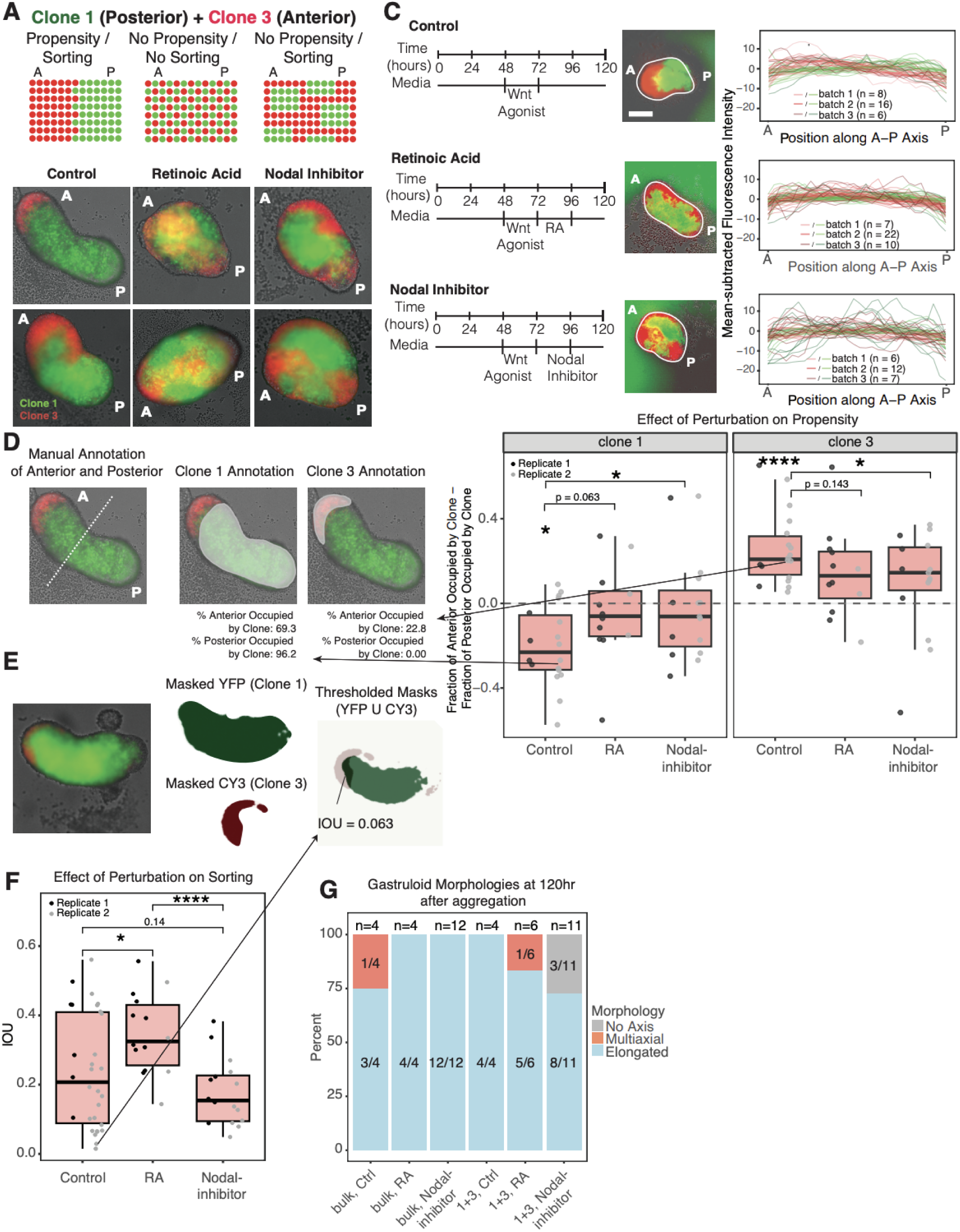
Perturbing Nodal signaling disrupts clones’ propensity while maintaining their sorting. **(A)** Schematic describing how cells might sort if they have propensity, how cells might lack propensity and fail to sort, or might sort without having propensity. **(B)** Representative images of gastruloids comprising clone 1 and clone 3 under control conditions, or when treated with retinoic acid or SB431542 (Nodal inhibitor) from 96-120 hours after aggregation. **(C)** Line scans describing the fluorescence intensities in red and green channels along the A-P axis for gastruloids under each condition. Normalization is described in **Methods**. Here, the mean of each line scan was subtracted to enable direct comparison of variation within each condition. Fluorescence intensities in the red channel were scaled to match the range of green fluorescence intensities. Scale bars for corresponding images correspond to 50px. **(D)** Quantification of the effect of chemical perturbation on gastruloid sorting. The fraction of the gastruloid anterior occupied by a given clone, minus the fraction of the gastruloid posterior occupied by that clone, is plotted for clone 1 and clone 3. Asterisks denote statistical significance comparing differences to zero,and statistical comparisons of chemical treatment against control are also shown (* p<0.05, ****p<0.0005). **(E)** Workflow for computing IOU. Masks for each channel are shown, along with composite images displaying masks for both channels (YFP colored in green, Cy3 colored in red). The region of overlap between masks is also shown, along with the IOU for each representative example. **(F)** Boxplots displaying IOU values for gastruloids under each condition. Asterisks denote statistically significant differences between conditions (* p<0.05, ****p<0.0005). **(G)** Stacked barplots showing the percentage of gastruloids that are elongated, multiaxial, or that have no axis. The plot shows bulk mES cells under control conditions, treated with RA, or treated with Nodal inhibitor and clone 1+3 gastruloids under control conditions, treated with RA, or treated with Nodal inhibitor.

To perturb RA and Nodal signaling, we administered, respectively, 0.15nM RA or 5µM SB-431542 (a disruptor of Activin/Nodal/TGF signaling) after aggregating gastruloids comprising anterior clone 3 and posterior clone 1. We applied perturbations from 96-120 hours after aggregation to minimize effects on initial gastruloid axis establishment and elongation. By 96 hours, *Wnt* signaling has been established and genes like *T* are expressed and polarized, so disrupting developmental signaling pathways should, in principle, have less of an effect on overall gastruloid development than if applied earlier^58,59^.

We first wanted to assess the effect of RA treatment or inhibition on propensity. Visually, RA treatment produced a "blended" appearance where clone boundaries became indistinct and cells from both clones were intermixed throughout the gastruloid **(Figure 5B)**. Tracing the fluorescence intensities for red and green channels along the A-P axis of control and treated gastruloids, RA treatment caused average red (clone 3) and green (clone 1) fluorescence intensity profiles to become more similar to one another, suggesting a loss of a difference in propensity between the two clones (**Supplemental Figure 22**). Though gastruloids comprising paired posterior clones 1 and 6 had some degree of mixing at baseline, RA treatment again collapsed the average fluorescence intensity curves for this pairing of clones (**Supplemental Figure 23**). We also tested 100nM AGN-193109, an RA inhibitor, which also resulted in a loss of differential propensity between clone 1 and clone 3 and clone 1 and clone 6 (**Supplemental Figure 24**). Activating or disrupting RA signaling may thus “wash out” differences between clones by either forcing them to a more anterior or posterior fate, respectively, resulting in a loss of differences in propensity between clones in a pair.

Unlike RA treatment, which blended clones throughout the gastruloid, Nodal inhibition preserved clear boundaries between clones. Visually, Nodal inhibition resulted in large, discrete patches where each region was dominated by a single clone, but the patches appeared throughout the A-P axis rather than consistently in anterior or posterior regions (**Figure 5B**). Fluorescence intensity profiles in Nodal-inhibited gastruloids were highly variable (**Figure 5C**), with large peaks in intensity corresponding to each clone’s main territories appearing at unpredictable positions along the A-P axis. To quantify the loss of propensity, we measured what fraction of each gastruloid’s anterior and posterior regions was occupied by each clone (described in **Methods**). In control gastruloids, clone 3 (anterior propensity) occupied more of the anterior region while clone 1 (posterior propensity) occupied more of the posterior region. However, like RA treatment, Nodal inhibition eliminated preferential localization of either clone to anterior or posterior regions (**Figure 5D**).

We next compared how RA treatment and Nodal inhibition affect clone sorting. We measured the degree of spatial overlap between clones. We generated binary masks from each fluorescent channel, YFP for clone 1 and Cy3 for clone 3, to identify the regions each clone occupied. We then calculated the intersection over union (IOU) between the two masks: the area of overlap divided by the total area covered by either mask. Higher IOU values indicate more overlap between clones, and therefore greater mixing (**Figure 5E-F**). RA-treated gastruloids showed significantly higher IOU values than both control and Nodal-inhibited gastruloids, consistent with a higher degree of mixing. In contrast, Nodal inhibition had little effect on IOU, indicating that clones remained spatially separated despite a loss of propensity. These results suggest that RA disrupted both propensity and sorting. However, Nodal inhibition primarily disrupted propensity but preserved some degree of sorting, demonstrating that the two processes can be uncoupled.

To determine whether the loss of propensity and sorting we observed was due to specific signaling disruption or a general failure of gastruloids to develop, we examined whether other aspects of gastruloid organization remained intact. Gastruloids treated with RA or Nodal inhibition could still elongate (**Figure 5G, Supplemental Figure 25A**) We also were able to confirm that RA treatment and Nodal inhibition could preserve the expression and polarization of *T,* indicating that basic morphogenetic processes can proceed even when propensities were lost (**Supplemental Figure 25B**). seqFISH analysis in SB-431542-treated bulk gastruloids (n = 2) revealed that *T* expression remained polarized, although we observed some coexpression in those *T*-positive cells with the anterior marker *Igfbp5* (**Supplemental Figure 25C**). Similarly, RA-treated chimeric gastruloids composed of clone 1 and clone 3 (n = 3) and RA-treated bulk-gastruloids (n=2) could polarize *T* expression but showed coexpression of *T* and *Igfbp5* (**Supplemental Figure 25D, E**). We also observed widespread expression of *Sox2* in RA-treated chimeric gastruloids composed of clone 1 and clone 3, a gene typically marking posterior identity in our mES cell line^60^, suggesting further differences between the RA-treated chimeric gastruloids and untreated gastruloids (**Supplemental Figure 25F**). Unlike the gene expression differences observed between clones in control chimeric gastruloids, both RA and Nodal inhibitor treatment reduced gene expression differences between clones (**Supplemental Figure 25H**). Therefore, even in gastruloids that could polarize T and elongate, propensities were disrupted under treatment. Overall, RA and Nodal signaling perturbations disrupted clonal organization while leaving some aspects of axial patterning intact, consistent with a role for morphogen signaling in regulating the spatial behavior of individual clones during gastruloid development.

### Markers of clonal propensities *in vivo*

A key feature of the division of labor we observed in gastruloids is that clones contributing to roles outside their propensity retain transcriptional signatures of their original bias, for example the "confused" expression patterns visible in pure-clone gastruloids and in anterior-biased clones contributing to the posterior of chimeric gastruloids. We wondered whether analogous signatures might be detectable *in vivo*^21,61^.^21,61^, We analyzed published single-cell RNA-seq data from E12.5 embryos in which epiblast cells had been barcoded (E5.5) and clonally traced through development (**Figure 6A-B**). In this dataset, cells were annotated by tissue type, and we further classified these tissues according to their germ layer of origin. Some barcodes were predominantly found in ectoderm-derived tissues, suggesting that these clones were biased toward ectoderm differentiation (ectoderm propensity). Others showed bias toward mesoderm or endoderm lineages, with correspondingly skewed tissue distributions (ME propensity) (**Supplemental Figure 26A-C**). We asked whether clones contributing to tissues outside their primary bias retained expression signatures of their preferred lineage, analogous to the mixed marker expression we observed in gastruloids.

**Figure 6.**
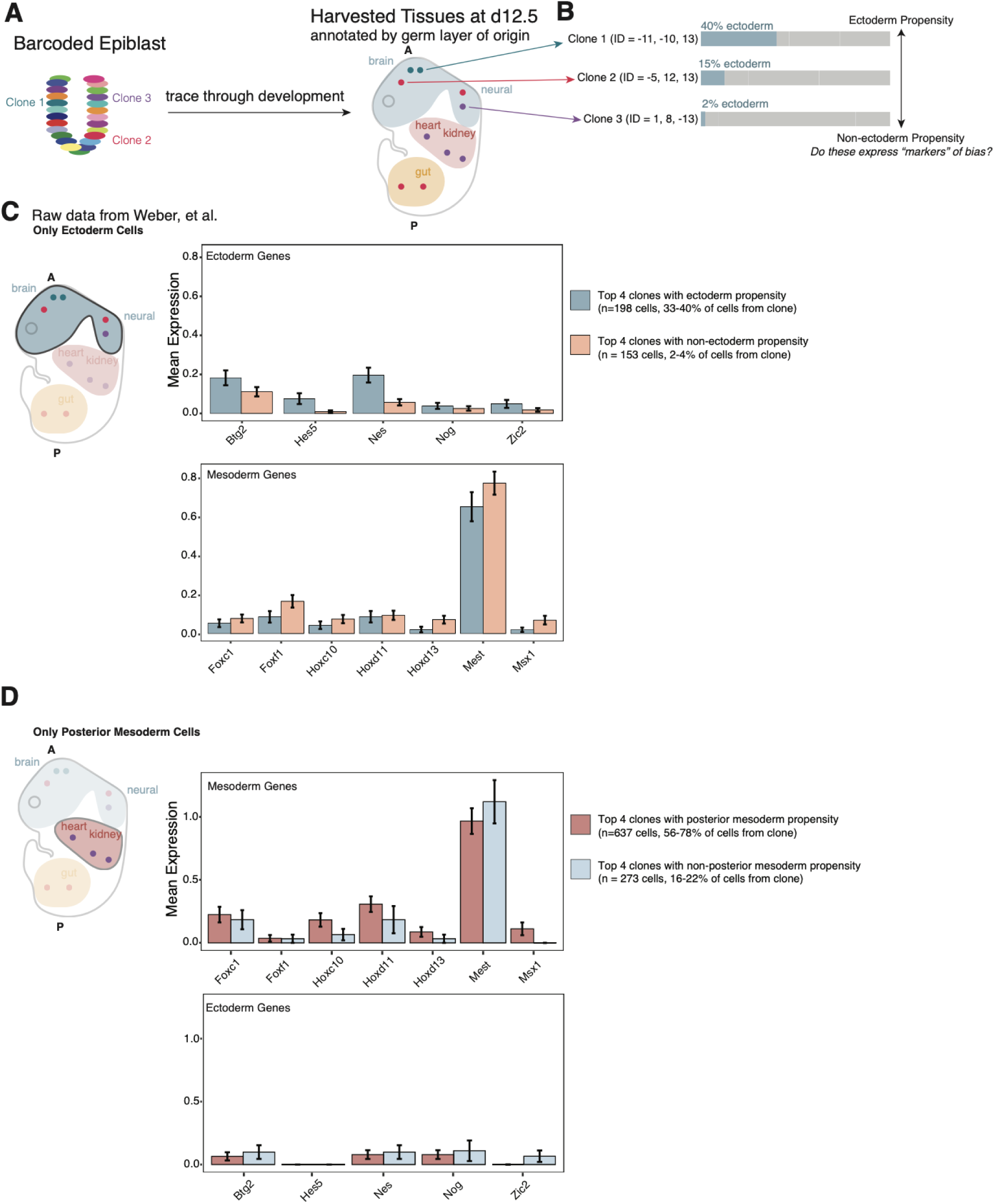
Analysis of an *in vivo* lineage tracing dataset reveals persistent bias through development. **(A)** Schematic depicting the conceptual workflow. **(B)** The fraction of cells belonging to ectoderm-derived tissues for three clones in the dataset. **(C)** For only cells annotated as belonging to ectoderm, derived tissues, barplots are shown comparing gene expression values for ectoderm markers (*top*) and mesoderm markers *(bottom*), comparing the top 4 clones with the highest proportion of cells in ectoderm tissues and the 4 clones with the lowest proportion of cells in ectoderm tissues. **(D)** For only cells annotated as belonging to posterior mesoderm, derived tissues, barplots are shown comparing gene expression values for mesoderm markers (*top*) and ectoderm markers *(bottom*), comparing the top 4 clones with the highest proportion of cells in posterior mesoderm tissues and the 4 clones with the lowest proportion of cells in posterior mesoderm tissues.

We first subsetted the dataset to examine gene expression exclusively within barcoded cells belonging to tissues derived from ectoderm (**Figure 6C**). In these ectoderm cells, we asked, do ectoderm cells from clones that produce fewer ectoderm cells (i.e., non-ectoderm propensity) still express markers of other lineages, and do clones biased toward ectoderm (ectoderm propensity) show stronger ectoderm marker expression? We compared the top four clones with the lowest contribution to ectoderm and the top four clones with the highest ectoderm contribution (**Supplemental Figure 26D**). We looked at marker genes selected based on established associations with ectodermal or mesodermal identity in the developing mouse embryo. Within ectodermal cells, clones with non-ectoderm propensity tended to show elevated expression of mesoderm-associated genes, while clones with ectoderm propensity tended to more strongly express ectoderm markers such as *Btg2*, *Hes5*, *Nes*, and *Zic2*. However, many differences were modest, as some genes showed comparable expression across groups (e.g., *Nog*, *Hoxd11*). Even though slight, the transcriptional differences we noted existed despite all analyzed cells being annotated as ectoderm, suggesting that early clonal identity leaves lasting transcriptional signatures that were not fully erased upon lineage commitment, much like the expression patterns we observed in our gastruloid experiments, where clones performing roles misaligned with their propensities showed mixed marker expression.

We next analyzed cells from tissues derived specifically from posterior mesoderm. First, we asked whether clones with propensities to form mesoderm-derived tissues inappropriately express ectodermal genes within posterior mesoderm cells. These effects were subtle because ectoderm-associated genes were expressed at low levels in posterior mesoderm for all the clones (**Figure 6D**). Next, we compared mesoderm marker gene expression between clones with a propensity toward posterior mesoderm (posterior mesoderm propensity) and those with propensities for other tissues. Six out of seven mesoderm markers were more highly expressed in clones with posterior mesoderm propensity, indicating that propensity is reflected in transcriptional signatures even within the same germ layer fate. While this analysis is descriptive, we present it as suggestive evidence that the division of labor framework may extend beyond gastruloids.

## Discussion

Here, we show that gastruloid morphogenesis operates through a division of labor among stem cell clones. Even though individual clones retain broad differentiation potential, when a clone is forced to fill every role in gastruloid development, even those for which it seemingly has an antogonistic propensity, gastruloids often fail to elongate properly. In mixed aggregates composed of bulk populations or defined pairs of clones, clones instead specialize in complementary fates aligned with their intrinsic biases, collectively producing organized structures.

There is an extensive literature around the concept of lineage priming, in which cells show some propensity towards a particular cell fate. Our results show that lineage priming is not, in and of itself, sufficient to drive cells towards a particular fate. Take, for instance, our results on comparative advantage. We show that when mixing clones that both have, say, a posterior propensity, one clone will reproducibly adopt the posterior propensity while the other will adopt anterior fates. Thus, the fate of a clone depends not entirely on their intrinsic bias, but rather their bias and the global context of the entire gastruloid. The same clone can adopt a different fate depending on what is collectively more optimal. These results suggest an emergent cooperative behavior between individual cell clones within the developing gastruloid.

Our findings notably point to cooperation among clones as a driving force for morphogenesis. Recent work^62,63^ has pointed to bottlenecks in polyclonal aggregates that reduce clonal complexity. Moreover, *in vivo*, a wave of apoptosis in the epiblast precedes gastrulation^64^. It is not our view that these processes of cooperation and competition are mutually exclusive^62^. In our study, chimeric aggregates did not show obvious evidence of clone takeover; perhaps, maximizing “collaborative” interactions among non-competitive clone combinations functions to eliminate potentially lethal lineage-specific defects, while allowing for proper morphogenesis via division of labor. Our findings build upon work by Braccioli et al.^27^ who demonstrated that communication between tissues is essential for proper gastruloid development, showing that wild-type cells can rescue the developmental defects of transcription factor knockout cells through chimeric interactions. Our study shows that even among genetically identical stem cells, clonal diversity creates specialized subpopulations that spontaneously divide labor during gastruloid development. Cooperation among intrinsically biased clones may be a fundamental organizing principle that underlies communication among tissues.

The division of labor that we observed in gastruloids appears to operate on similar principles to the ability of pluripotent stem cells to rescue lethal organ deficiencies via blastocyst complementation. A number of emerging studies have explored the use of stem cells to promote the development of lungs^10^, heart^65^, liver^66^, pancreas^8^, and eyes^9^, in embryos that were engineered to otherwise lack these organs. Exogenous stem cells could rescue organogenesis, preferentially forming cell types belonging to the organs that the embryos were unable to make themselves. Just like our observations in gastruloids, organogenesis *relied* on division of labor among a heterogeneous cell population in the context of rescuing organ development. Current regenerative medicine approaches often assume that homogeneous cell populations are optimal for therapeutic applications^67–69^. However, if natural development relies on coordinated heterogeneity, then therapeutic strategies might benefit from harnessing rather than eliminating cellular diversity. Our work establishing a framework for describing beneficial clonal interactions in development could inform the design of more effective stem cell therapies and tissue engineering approaches that use the ability of cells to cooperate.

Differences in propensities between individual clones must arise from molecular differences between the clone-originating ES cells, and these differences must further be stably maintained for several divisions. Much recent work has identified molecular differences among ES cells that correlate with differences in their fate outcomes^4,5,70^, although it has remained challenging to definitively prove that these differences represent fluctuations instead of partially differentiated cells^31^. Regardless, while it has been theorized that development may benefit from the interactions of cells with different propensities and states, these theories have largely remained untested. We demonstrate that heterogeneity can positively affect developmental outcomes, not simply because a diverse population is more likely to contain cells capable of filling each role, but rather because differently biased cells actively specialize in complementary roles. This principle may extend beyond gastruloids to other self-organizing systems. The combination of intrinsic cellular diversity with intercellular communication creates opportunities for emergent properties that exceed the capabilities of individual components. Rather than requiring perfect uniformity, developmental precision may actually depend on the orchestrated behavior of diverse cellular players, each contributing their specialized capabilities to the collective task of morphogenesis.

Our spatial transcriptomics experiments revealed that pure-clone gastruloids showed disrupted spatial organization of marker genes, with lower Moran’s I values indicating poor spatial clustering, and positive L-metric values revealing coexpression of normally mutually exclusive anterior and posterior markers like *T* and *Igfbp5*. Such a "confused" gene expression pattern suggests that when clones are forced to generate all cell types rather than specializing in their preferred fates, they produce cells with mixed identity signatures.

Importantly, combining clones largely rescued defects in spatial gene expression. Chimeric gastruloids showed Moran’s I values similar to bulk controls and restored mutual exclusivity between anterior and posterior markers. However, some residual signatures of propensity persisted even in successful chimeric gastruloids, with clone identity still detectable through subtle differences in marker gene expression levels. For instance, chimeric gastruloids composed of posterior clone 1 and posterior clone 6 exhibited relatively sparse expression of the anterior marker *Igfbp5*, consistent with both clones’ posterior propensity. In gastruloids combining clone 1 with anterior clone 3, cells from clone 3 occasionally coexpressed *Igfbp5* and *T*. Thus, clones preferentially contribute to lineages aligned with their propensity, but retain molecular signatures of their intrinsic propensity even when filling misaligned roles.

Signatures of propensity become most apparent when clones must deviate from their preferred roles. Just as clone 3 cells revealed their anterior identity by maintaining *Igfbp5* expression in *T*-expressing populations, clones with a propensity for ectoderm formation *in vivo* showed stronger expression of ectodermal markers within ectoderm-derived tissues, whereas clones less likely to form ectoderm expressed mesoderm markers in these tissues. We therefore propose a model whereby clones’ propensities guide their roles in development and form a basis for how they organize their respective fates.

Our findings align with recent work by Regalado et al.^5^, which highlights how individual mES cells are remarkably heterogeneous, forming monoclonal gastruloids that can be biased toward mesodermal, neural, or other lineages. Relatedly, work from McNamara et al.^26^ showed that differential Wnt signaling activity can shape individual cell contributions to gastruloids. These studies and others^42,60^ point to the existence of heterogeneity in differentiation decisions among ES cells, but leave open the question of how heterogeneity is integrated across the population to produce consistent, reproducible morphogenetic outcomes. We connect lineage information with spatial gene expression, explicitly showing how lineage-dependent fate biases affect the spatial organization of tissues within the gastruloid.

What are the molecular differences among clones that underlie their propensities? We found that clonal propensities were accompanied by subtle but reproducible differences in chromatin accessibility. In line with recent work^26,56,71^, differentially-accessible loci included those involved in Wnt and RA signaling. Interestingly, we observed that propensity degraded with passaging, suggesting that though persistent, it was not permanently encoded. In contrast, we observed that overall chromatin accessibility differences between clones with anterior propensity and those with posterior propensity persisted across passaging. It is possible that the majority (or even all) of these differences do not have functional consequences. Instead, it may be that only the specific epigenetic differences at developmentally relevant motifs drive propensity. We saw that chromatin accessibility differences at motifs related to Wnt, RA, and Nodal signaling were present even though clones retained similar expression of pluripotency-related genes, suggesting that the potential for propensity is present in chromatin landscapes even before overt lineage divergence.

Our ATAC-seq results indicate that clones’ spatial propensities may reflect altered responsiveness to developmental cues like Wnt, RA, and Nodal signaling. Indeed, clones’ propensities could be disrupted by RA and Nodal. RA treatment produced a "blended" phenotype where clone boundaries became indistinct and cells intermixed throughout the gastruloid, while Nodal inhibition maintained clear clonal boundaries but randomized their spatial arrangement along the A-P axis. These differential effects demonstrate that spatial propensity and sorting (segregation of clones from one another) can be uncoupled, and suggest that different signaling pathways regulate distinct aspects of clonal organization. Clones differ in how they interpret and respond to instructive signals, with consequences for tissue-level patterning under varying morphogen contexts.

We have demonstrated that economic principles like division of labor and comparative advantage can operate in development, and these concepts may govern how a populations of individually biased cells collectively produce consistent, organized structures^20^. Future work may reveal the intercellular communication mechanisms that mediate these global organizational strategies.

## Data and code availability

All raw and processed data, as well as code for analyses performed in this manuscript, can be found using this link: https://www.dropbox.com/scl/fo/yklfs2p4s9dg8uyb6w8nt/ADXsVD7ABiklQnTyCSmhU5Q?rlkey=gl8drm3l79j9k7oi1wuro4s6w&st=87jmyzpt&dl=0

## Acknowledgements

We thank all members of the Raj lab for scientific discussion and comments on the manuscript and data visualization. We thank Aoife O’ Farrell and Miles Arnett for their discussion and assistance with the ATAC library preparation workflow, Yael Heyman and Grant Kinsler for their discussion and assistance with the seqFISH workflow, the William Greenleaf lab for sharing expertise in ATAC sequencing and providing us with custom primers used in our ATAC-seq experiments, and the Sydney Shaffer lab, particularly Sydney Shaffer, Christopher Coté, and Connor Hennessey, for assistance and equipment with sequencing and seqFISH. We thank the Jared Toettcher lab for their discussion and expertise with the gastruloid workflow, as well as for their gift of the E14-tg2a cell line. We thank Harry McNamara and Sedona Murphy for their helpful discussions and insights. Custom probes and kits for seqFISH experiments were designed by Spatial Genomics.

A.R. acknowledges support from a center grant from the Mark Foundation for Cancer Research, NIH Director’s Transformative Research Award R01 GM137425, NIH R01 CA238237, NIH R01 CA232256, and NIH 4DN U01 DK127405. V.A. acknowledges support from NIH T32 148377. C.G.T. acknowledges support from the Damon Runyon Cancer Research Foundation, DRG-2465-22.

## Author contributions

V.A. and A.R. conceived and designed the project under the supervision of A.R. V.A. designed and performed experiments with help from C.G.T. C.G.T designed and optimized lineage labels. K.S. generated Tn5 enzyme for ATAC-seq experiments. V.A. and A.R. wrote the manuscript with input from all authors. All authors read and approved the final manuscript.

## Declaration of generative AI and AI-assisted technologies

During the preparation of this work, the authors used ChatGPT and Claude to generate and improve code for analyses and to suggest improvements to the clarity and flow of the text. The authors reviewed and edited the content and take full responsibility for the content of the manuscript.

## Materials and Methods

### Cell Culture

Experiments were performed using cell lines generated from E14-tg2a mouse embryonic stem cells (ATCC CRL-1821). mES cells were thawed and plated on 25cm^2^ cell culture flasks coated with 0.1% gelatin. Cells were grown in 2i + LIF media comprising GMEM (Millipore Sigma, G6148) supplemented with 10% ESC qualified fetal bovine serum (R&D Systems, S10250), 1x GlutaMAX (Gibco, 35050–061), 1x MEM non-essential amino acids (Gibco, 11140– 470 050), 1 mM sodium pyruvate (Gibco, 11360–070), 100 μM 2-mercaptoethanol (Gibco, 471 21985–023), and 100 units/mL penicillin/streptomycin (Gibco, 15140–122), 1000 units/mL LIF (Millipore 473 Sigma, ESG1107), 2 μM PD0325901 (Tocris, 4192), and 3 μM CHIR99021 (Tocris, 4423). Cells were maintained between 20% and 80% confluency and were passaged by aspirating growth media, washing with phosphate buffered saline (PBS, Gibco, 14190144), and trypsinizing for 5 minutes at 37C (TrypLE Express, Gibco, 12605028). Trypsin was quenched with 2i+LIF media and cells were subsequently centrifuged at 500rpm for 5 minutes and pelleted. Supernatant was aspirated and cells were resuspended in fresh media prior to replating. Passage ratios varied from 1:5 to 1:10.

For experiments involving bottlenecked mES cell clones, cells from a polyclonal population were passaged into 96-well plates at a density of 1 cell per well. These were grown for ∼2 weeks and colonies were imaged using an Incucyte S3 Live Cell Imaging Analysis System (Sartorius) with a 4× objective. Viable colonies were serially expanded into 24-well, 6-well plates and then 25cm^2^ cell culture flasks. These were allowed to grow before being used for experiments or frozen down for subsequent analysis. With the exception of experiments testing late-passage clones, experiments were performed on clones passaged 1-2 times after freezing, and vials were frozen at each round of passaging. Cell cultures were mycoplasma negative.

### Gastruloid Protocol

Gastruloids were grown in N2B27 media (1:1 mixture of DMEM/F-12 (Gibco, 11320033) and neurobasal medium (Gibco, 21103049), supplemented with 100 μM 2-mercaptoethanol, 1:100 N-2 (Gibco, 17502048), 1:50 B-27 (Gibco, 17504044), and 100 units/mL penicillin/streptomycin (Gibco, 15140–122)). E14-tg2a cells were trypsinized and pelleted, then washed twice with PBS with each wash followed by a 5-minute centrifugation at 500rpm. Cells were resuspended in N2B27 media and counted. To form gastruloids, 300 single cells in 40µl N2B27 media were pipetted into each well of an ultra-low attachment 96-well round-bottom microplate (Corning, 7007). These plates were then transferred to a cell culture incubator for 48h, after which each aggregate was administered 140µL of N2B27 media supplemented with 3µM Chiron. Media was replaced with 140µl fresh N2B27 without Chiron 24 hours afterward (at 72 hours after aggregation), and subsequently every 24 hours for the remaining 48 hours of the gastruloid protocol. Gastruloids were assayed either by imaging using an Incucyte, or by immunofluorescence, in which case samples were fixed for 2 hours in 4% PFA at 4 °C before being washed twice with PBS, or by seqFISH in which samples were fixed in 4% PFA before being permeabilized in 70% ethanol before embedding.

### Lineage Labeling

To generate the nuclear-localized fluorescent reporter constructs, eGFP, mOrange2, and mKate2 fluorescent proteins were cloned into a lentiviral backbone under the control of the EF1α promoter, as has been described^72^. Sequences encoding fluorophores were flanked by the SV40 nuclear localization signal. The constructs included the woodchuck hepatitis virus posttranscriptional regulatory element (WPRE). A β-lactamase (AmpR) resistance gene was used for bacterial selection.

Plasmids were amplified using plasmid electroporation into Endura electrocompetent Escherichia coli cells (Lucigen) using a Gene Pulser Xcell (Bio-Rad). An AmpR resistant clone was isolated and grown at 32 °C for 12-14 hours with agitation. Plasmids were isolated from the pelleted cells using the EndoFree Plasmid Maxi Kit (Qiagen, 12362) per the manufacturer’s protocol.

Lentivirus was produced by transfecting Lenti-X 293T cells with helper plasmids VSVG and psPAX2 using polyethylenimine (Polysciences 23966) in Opti-MEM (Thermo Fisher, #31985062). A mass ratio of 4:2:3 for plasmid DNA:VSVG:psPAX2 was used. The transfection mix was mixed and incubated for 15 minutes at room temperature before being added dropwise to the cell monolayer. Lentivirus was collected 24, 48, and 72 hours after transfection, pooled, filtered, and concentrated using ultracentrifugation. Lentiviral stocks were stored at −80 °C.

### Morphology Analysis

Experiments were conducted on an Incucyte S3 Live Cell Imaging Analysis System (Sartorius) with a 4× objective, which captures a single focal plane per well without user selection of imaging plane or orientation. Because gastruloids settle to the bottom of round-bottom wells in a consistent orientation, the imaging plane is determined by well geometry rather than user choice. Phase and fluorescence images were acquired, with 300ms exposure for GFP and 400ms exposure for mCherry. Images were exported as 8-bit TIFs for phase images and 16-bit TIFs for fluorescent images. A custom script (https://github.com/arjunrajlaboratory/process_incucyte_tiff_data) was used to upload and analyze data with NimbusImage (https://www.nimbusimage.com/).

Gastruloid segmentation was performed manually, and gastruloids were annotated as “normal” if they were elongated along a single axis, “multiaxial” if multiple axes of elongation or projections were present, and “no symmetry breaking” if no obvious axis had formed or if gastruloids assumed irregular shapes.

For experiments involving chimeric gastruloids, a line was drawn along the gastruloid’s long axis, and fluorescent intensities were measured along the length of each line. NimbusImage annotations were exported in JSON format, and fluorescence line-scans were exported as CSVs for subsequent analysis. The posterior region of the A-P axis was defined either by expression of T (eg, in immunofluorescence experiments, below) or manually by gastruloid morphology. Samples that were not amenable to this line-scan analysis or A/P annotation due to a failure to elongate or due to a lack of any clear anteroposterior axis were excluded from analysis. For plots including individual line scans and multiple channels (eg, red and green), the green channel was scaled down since intensity measurements in this channel were consistently higher. This scaling factor was 10 by default, and 20 for batches 1 and 3 in line scans showing the effect of RA treatment and Nodal inhibition.

Clones 2 and 4 were excluded from analyses of propensity due to their dim fluorescence. Gastruloids generated from clone 2 (red) occasionally lost fluorescence as it elongated, making assessment of propensity unreliable. While clone 4 (red) did often appear in the posterior of chimeric gastruloids generated from bulk and clone 4 (**Supplementary Figure 2F**), it was also very dim, and hence hard to tell apart from background in many/most images of chimeric gastruloids.

### Gastruloid Embedding

Gastruloids were fixed as described above in 4% PFA (2 hours for experiments involving pure clone gastruloids, as these were done alongside immunofluorescence, and 15 minutes for experiments involving combined-clone gastruloids) before being transferred to 70% ethanol. Gastruloids were first pooled in a low-bind tube and then washed with buffer containing 10% formamide, 2x SSC, and 0.2% tween-20, and then with PBS with 0.2% tween-20 10mL of a gel solution including 750µl of 5M NaCl, 625µl of 1M Tris-HCL, 1250µl of 40% bis-acrylamide and 9.9ml water was made. Gastruloids were resuspended in 30µl of gel, activated using 37.5 µl of 10% ammonium persulfate and 18.75µl tetramethylethylenediamine (TEMED), and spotted onto proprietary functionalized coverslips made by SpatialGenomics, anchored by a coverglass coated with GelSlick (Lonza, #50640). Embedded gastruloids were incubated at 37 °C for 2 hours. Samples were washed with PBS, covered with 8% SDS and allowed to incubate for 1 minute before being washed again with PBS and incubated for 1 hour at 37 °C with Proteinase K (Thermo Fisher, #EO0491, 3µl diluted into 3mL of PBS with 2% Tween-20, diluted 100-fold further into 8mL PBS with Tween-20). Samples were washed with PBS twice, and primary probes were added to each sample, which was incubated at 37 °C for 24 hours. Slides were next used to assemble a flow cell (components provided by SpatialGenomics), rinsed with a wash buffer, incubated with DAPI for 1 hour at 37 °C, and washed with a kit-provided rinse buffer before imaging.

### Sequential FISH

Spatial transcriptomics experiments were performed using sequential FISH using technology from SpatialGenomics. We used a small custom panel of marker genes guided by recent single-cell RNA-seq studies^42^: *T*, *Hoxc8*, *Hoxb9*, *Sox2*, *Cyp26a1*, *Igfbp5*, *Wnt3*, *Sfrp1*, *Ncam1*, *Sox9*, *Lhx1*, *Nrp2*, *Thy1*, *Gbx2*, *Pim2*, *Utf1*, *Zfp42*, *Mesp1*, *Epha5*, *Hoxaas3*. The panel was designed in collaboration with SpatialGenomics and synthesized by them. All genes were identified sequentially via single-molecule FISH. seqFish was performed using SpatialGenomics’ GenePS instrument (software v1.6.0), and images were acquired at a z-offset of 1-2µm to ensure adequately-focused nuclei and spot detection; all gastruloids within a given experiment were processed and imaged identically.

### Sequential FISH analysis

All image processing was done either on the GenePS machine or with SpatialGenomics’ proprietary software (software v0.9.0). Raw images were aligned across multiple rounds of hybridization to form composite images, and each channel was manually thresholded before spot-counting. In each experiment, genes that were either poorly detected or had high background were removed from the analysis. Transcript identities were decoded, and nuclei were segmented using DAPI images (or GFP images, for experiments involving segmenting only clone 1) via CellPose^73,74^. A mask was formed from segmented nuclei that included the nucleus plus a dilation around the nucleus, allowing individual transcripts to be assigned to individual cells, yielding a cell-by-gene count matrix along with nuclear xy-locations. For chimeric gastruloids where clone 1 was segmented, each cell was segmented using DAPI and GFP, as above, so for each GFP-annotated cell, the closest DAPI-annotated cell was annotated as belonging to clone 1. Subsequent analysis included the custom SGAnalysis package (https://github.com/arjunrajlaboratory/SGanalysis). Moran’s I was calculated using the *lctools* package in R.

L-metric scores were computed using the custom l-metric package (https://github.com/arjunrajlaboratory/l-metric). For each cell, counts from each pair of genes are read in, yielding a list of gene expression values for both genes across all cells in the dataset. These were sorted on the basis of the first gene in the pair. Three vectors were generated for gene 2: one exclusively containing the mean expression of gene 2, another sorted from lowest to highest, and a third sorted from highest to lowest. Cumulative sums for gene 1 and gene 2 (sorted on the expression of gene 1) are computed to compare how gene 2 accumulates relative to itself, a flat line, and the low-to-high and high-to-low-ranked vectors. The area between the cumulative sum for gene 2 and the flat line is computed and normalized by the low-to-high and high-to-low-ranked vectors. Negative values correspond to mutually exclusive gene expression, resulting in an L-shaped scatter of gene 1 vs. gene 2, and positive values correspond to coexpression of gene 1 and gene 2.

To further compare coexpression patterns between experimental groups, we used permutation testing based on pairwise Jensen-Shannon divergence (JSD). For each pair of genes, we computed a two-dimensional histogram over a fixed 100-by-100 bin grid spanning the central 98% of the pooled expression range among all samples. Histogram counts were normalized to sum to 1 to produce a discrete probability distribution over the joint gene expression space for each gastruloid. We then computed the Jensen-Shannon divergence between each pair of histograms, labeling these as within-group (e.g., bulk sample vs. bulk sample) or between-group (e.g., bulk vs. clone 3). We used the difference between the mean within-group and between-group JSDs as a test statistic. We then performed a nonparametric permutation test whereby group labels were shuffled 5000 times, computing the test statistic at each permutation to generate a null distribution. The p-value was computed as the proportion of permuted test statistics greater than or equal to the observed test statistic.

### Immunofluorescence of mES Cell Monolayers

Cells, plated on 24-well imaging plates (P24-0-N), were fixed in 70% ethanol and stored at 4C. For immunofluorescence, ethanol was aspirated and samples were washed with PBS and incubated for 5 minutes at room temperature. Samples were then washed with PBS with 0.1% Triton-X reagent. Samples were then incubated for 1 hour with primary antibody (anti-beta catenin: mouse anti-beta catenin antibody, Invitrogen #13-8400, anti-Sox17: goat anti-Sox17 antibody, R&D #AF1924; anti-T/Brachyury: Human/Mouse Goat anti-Brachyury, R&D #AF2085; anti-Pax6: rabbit anti-Pax6 antibody, Abcam ab195045) for 1 hour at a 1:200 dilution in PBS. Samples were then washed 3x with PBS with 0.1% Triton-X reagents, with 5 minutes in between washes. We then incubated for another hour with secondary antibody (A488 anti-mouse, Cell Signaling Technologies #4408S; A594 anti-mouse, Cell Signaling Technologies #8890; A488 donkey anti-sheep, Life Technologies #A11015; A647 goat anti-rabbit, Invitrogen #A21244) at a 1:400 dilution in PBS. Samples were then washed 3x with PBS, with 5 minutes in between washes. For the second PBS wash, DAPI was added. PBS was replaced with 2xSSC, and samples were imaged at 20X on a Nikon Ti-E with a 10X Plan-Apo objective with filter sets for Cy3, Atto647N, Alexa594, and Alexa488. Images were acquired with a 0.45 numerical aperture.

### Gastruloid Immunofluorescence

Gastruloids were fixed in 4% PFA for 2 hours at 4C as described above and transferred to PBS with 0.2% Triton-X (PBST) for overnight incubation at 4C. We next added primary antibody (Human/Mouse Goat anti-Brachyury, R&D #AF2085) at a 1:400 dilution and incubated overnight at 4C. We next washed out the primary antibody, first with three back-to-back washes with PBSFT, followed by three washes of PBSFT spaced 1 hour apart, to allow the primary antibody to diffuse out of the gastruloid. We next administered secondary antibody (A647 donkey anti-sheep, Invitrogen #A21448) at a 1:400 dilution and incubated overnight at 4C. Next, we again washed out the secondary antibody with three back-to-back PBSFT washes, followed by three washes of PBSFT spaced 1 hour apart before imaging. Samples were then transferred to a glass-bottom 8-chambered imaging slide for imaging (Cellvis C8-1-N). Samples were imaged on a Nikon Ti-E with a 10X Plan-Apo objective with filter sets for Cy3, Atto647N, Alexa594, and Alexa488. Images were acquired with a 0.45 numerical aperture objective. Seven z-slices were obtained per image, and images were maximum-intensity projected to generate images for visualization.

### Gastruloid Live Imaging

Gastruloids were immobilized in Matrigel for live imaging. Matrigel matrix (Corning #356231) was thawed overnight at 4 °C. Glass bottom 8-chambered imaging slides were chilled to 4C, and each chamber was coated with Matrigel reagent and then transferred to ice. Gastruloids were collected using 200µl pipette tips with the ends cut off and dispersed into the cooled Matrigel within each chamber. Slides were then warmed at 37 °C for 10 minutes to solidify the Matrigel, and 100µl N2B27 medium was added to each chamber. Samples were imaged using a Nikon Ti-E with a 4X or 10X Plan-Apo objective and kept in a microscope-mounted environmental chamber with controlled temperature, CO_2_, and humidity. Images were acquired every 60-90 minutes over 24-48 hours. Images were maximum-intensity projected to generate final movies.

### Differentiation Assay

For endoderm induction^75^, DMEM (Thermo Fisher, 11995065) was supplemented with 0.5% ESC-qualified fetal bovine serum, 100ng/mL activin A (Fisher Scientific 338AC010), and GlutaMAX. For mesoderm induction, DMEM was supplemented with 20ng/mL BMP4 (Fisher Scientific 315-27), 0.5ng/mL activin A, and 2.5ng/mL bFGF (Fisher Scientific PHG0367). For neural induction, we cultured cells in Neurobasal medium with Neural Induction Supplement (Gibco A1547701). Cells were trypsinized and plated at a density of 50,000 cells/well in 24-well glass-bottom plates (CellVis P24-0-N) containing the respective induction media. Cells were imaged every 24 hours for 72 hours following plating, and the media was refreshed every 24 hours.

### RNA sequencing and analysis

We sequenced mRNA from clonal E14-tg2a mES cell lines. As clones were expanded, cells were collected at a density of 75,000 cells per sample and stored in QIAzol. Samples from a bulk population were also harvested as a control. RNA was extracted (Direct-zol RNA-mini kit, Zymo Research, #R2072) We used the Watchmaker mRNA library prep kit, indexing samples using xGen Stubby Adapter-UDI primers for Element (IDT, #10017037) and sequenced each sample at a depth of ∼20 million reads on an Element AVITI sequencing instrument (75+75bp, high-output Cloudbreak Freestyle). We pseudoaligned reads to mm10 using *Kallisto* and imported the output into R via *tximport*. We used the Relative Log Expression (RLE) and calculated counts per million (CPM) per gene. To compare the expression of pluripotency markers between clonal mES cells and bulk, the genes *Dnmt3l*, *Dppa3*, *Dusp6*, *Esrrb*, *Klf4*, *Myc*, *Nanog*, *Otx1*, *Pou5f1*, *Sox2*, *Tet2*, *Utf1*, *Zfp42*, and *Zic2* were used. To compare expression of gastruloid cell type markers, we used the genes *Igfbp5*, *Gata6*, *Aldh1a2*, *Hoxd4*, *Hoxb9*, *Hoxc8*, *Wnt3a*, *Cyp26a1*, *Sox2*, and *T*.

For bulk RNA-sequencing of 48h gastruloids, gastruloids were grown in 96-well plates, with 288 gastruloids each per clone. Gastruloids for each clone were pooled and centrifuged for 5min at 300xg, resuspended in PBS. After cell counts were taken, 100,000 cells per sample were centrifuged again at 300xg for 5min and resuspended in Zymo DNA/RNA shield (R1100-50). Samples were subsequently sent to Plasmidsaurus for 3’-end RNA sequencing and processed using their pipeline. Briefly, FASTQ files were assessed for quality using FastQC v0.1.12.1. Quality filtering was performed using fastp v0.24.0 with poly-X tail trimming, 3’ quality-based tail trimming, a minimum Phred quality score of 15, and a minimum length requirement of 50 bp. Quality-filtered reads were aligned to the mm10 mouse genome using STAR aligner v2.7.11 with non-canonical splice junction removal and output of unmapped reads, followed by coordinate sorting using samtools v1.22.1. tail trimming, a minimum Phred quality score of 15, and a minimum length requirement of 50 bp. UMIcollapsev1.1.0 was used to remove PCR and optical duplicates, and RSeQC v5.0.4 and Qualimap v2.3 were used to assess alignment quality metrics, strand specificity, and read distribution across genomic features. Gene-level expression quantification was performed using featureCounts (subread package v2.1.1), with strand-specific counting, multi-mapping read fractional assignment, exons and three prime UTR as feature identifiers. Sample-sample correlations for PCA were calculated on normalized counts (TMM, trimmed mean of M-values) using Pearson correlation. Differential expression was performed using edgeR v4.0.16 using filtering for low-expressed genes with edgeR::filterByExprwith default values.

### ATAC sequencing

We performed ATAC sequencing based on a modified protocol based on the Omni-ATAC-seq method^76^. At each round of passaging that we tested, we collected 50,000 cells per technical replicate and lysed these in a buffer containing 10% NP-40, 10% Tween-20, and 1% Digitonin for 3 minutes at 4C. We transposed lysed cells with Tn5 (a gift from Dr. Kavitha Sarma, Wistar Institute) for 30 minutes at 37 °C. Transposed DNA was isolated using a Zymo DNA Clean and Concentrator kit, and then amplified using custom primers (**Supplemental Table 1**, originally designed by Greenleaf Lab, Stanford University) for 13 PCR cycles. Libraries were purified using double-sided selection with AMPure XP magnetic beads, and quantified using BioAnalyzer High Sensitivity DNA chips. We sequenced libraries at a depth of ∼50 million paired-end reads per sample on a NextSeq2000 with a P4 XLEAP 100-cycle kit.

### ATAC sequencing analysis

After aligning reads to mm10 using bowtie2. We called peaks using MACS3 (version 3.0.3), keeping an FDR cutoff of 0.001 and filtering to a fragment size less than 150bp. We generated one nonredundant peak list filtered to those found in at least two samples and that were not listed in the ENCODE blacklist. We then used GenomicAlignment’s *summarizeOverlaps* function to generate a counts matrix that included all samples. Peaks were annotated using *annotatePeak* from the ChIPseeker package^77^ and were mapped to their closest gene transcription start site. Upset plots were generated after making peak lists where, for each condition, at least one technical replicate had that peak for all clones within that condition. We performed differential peak analysis using DEseq2. For all comparisons tested, we extracted a peak list of differential peaks (|log2FC| > 1) and used this peak list as input to HOMER^78^, using default parameters for de novo motif discovery.

Transcription factor footprinting was performed using TOBIAS. BAM files from replicate samples were merged by condition: passage 1 anterior clones, passage 1 posterior clones, passage 3 anterior clones, passage 3 posterior clones, and bulk. Merged BAM files were corrected for Tn5 insertion bias using TOBIAS ATACorrect with the mm10 reference genome. Differential TF binding between anterior and posterior clones at each passage was assessed using TOBIAS BINDetect with position weight matrices from the JASPAR 2024 vertebrate database, with default parameters used for the analysis.

### Channel IOU Analysis

To test whether different chemical perturbations affected patterning of clones beyond their arrangement along the A-P axis, we segmented gastruloids and generated masks within each channel using Otsu thresholding, via the scikit-image Python package^79^. The number of pixels shared within fluorescent channel masks was computed as the intersection between those masks, and the number of pixels included within the composite of the masks defined the union of those masks. The quotient between intersection and union was compared across experimental conditions.

### Re-analysis of Weber et al

We analyzed a publicly available single-cell RNA-seq dataset generated by Weber et al^61^. The dataset came annotated, such that each cell had been assigned its clone barcode and a tissue type. We assigned each tissue type to its germ layer of origin. Ectoderm was assigned to migrating neural crest, enteric neurons, differentiating neurons, dorsal root ganglia, neuronal/glial progenitors, GABAergic neurons, neural crest, and neuroblasts. Endoderm was assigned to epithelial cells. Anterior mesoderm was assigned to pulmonary/gastrointestinal mesenchyme, ovarian mesenchyme, smooth muscle, valve/endocardial cushion, perivascular cells, myocardium, mesentery/epicardium, fibroblast pole, and erythrocytes - definitive. Posterior mesoderm was assigned to renal mesenchyme, nephron progenitor, musculoskeletal mesenchyme, skeletal muscle, erythrocytes - primitive, tendon/ligament pole, chondrocyte pole, pericytes, meningeal fibroblasts, and skeletal mesenchyme. Mesoderm, mixed, was assigned undifferentiated/proliferative mesenchyme, megakaryocytes/hematopoietic progenitors, and macrophages. We filtered clones on the basis of a dataset-provided exclusion list, and to remove barcodes with fewer than 5 cells assigned. The proportion of cells within each germ layer was calculated for each clone. Clones were then ranked on the basis of the fraction of their cells that belong to ectoderm-derived tissues.

We next subset our cell-by-gene matrix to only include cells from, respectively, ectoderm, or in another analysis, posterior mesoderm. Expression of ectoderm/anterior genes *Hes5, Btg2, Cenpf, Chrd, Nog, Nes, Zic2, Hoxa1, Pax6, Sfrp1, Dkk1, Tbx1*; and posterior mesoderm genes *Foxf1, Foxc1, Foxd1, Pbx1, Meis2, Mest, Cdh2, Hoxc10, Hoxd11*, *Eomes* was compared across clones.

## References

1. Hölldobler, B. & Wilson, E. O. The Ants. (Harvard University Press, 1990).

2. West, S. A. & Cooper, G. A. Division of labour in microorganisms: an evolutionary perspective. Nat Rev Microbiol 14, 716–723 (2016).

3. Janiszewska, M. et al. Subclonal cooperation drives metastasis by modulating local and systemic immune microenvironments. Nat Cell Biol 21, 879–888 (2019).

4. Junyent, S. et al. The first two blastomeres contribute unequally to the human embryo. Cell 187, 2838–2854.e17 (2024).

5. Regalado, S. G., et al. Lineage recording in monoclonal gastruloids reveals heritable modes of early development. bioRxiv (2025) doi:10.1101/2025.05.23.655664.

6. Lamba, A. & Zernicka-Goetz, M. The role of polarization and early heterogeneities in the mammalian first cell fate decision. Curr Top Dev Biol 154, 169–196 (2023).

7. Founta, K.-M. & Papanayotou, C. In Vivo Generation of Organs by Blastocyst Complementation: Advances and Challenges. International Journal of Stem Cells 15, 113 (2021).

8. Matsunari, H. et al. Blastocyst complementation generates exogenic pancreas in vivo in apancreatic cloned pigs. Proceedings of the National Academy of Sciences 110, 4557–4562 (2013).

9. Zhang, H. et al. Rescuing ocular development in an anophthalmic pig by blastocyst complementation. EMBO Molecular Medicine (2018) doi:10.15252/emmm.201808861.

10. Mori, M. et al. Generation of functional lungs via conditional blastocyst complementation using pluripotent stem cells. Nature Medicine 25, 1691–1698 (2019).

11. Arias, A. M., Marikawa, Y. & Moris, N. Gastruloids: Pluripotent stem cell models of mammalian gastrulation and embryo engineering. Dev. Biol. 488, 35–46 (2022).

12. Anlaş, K. et al. Early autonomous patterning of the anteroposterior axis in gastruloids. Development 151, (2024).

13. Bennabi, I. et al. Size-dependent temporal decoupling of morphogenesis and transcriptional programs in pseudoembryos. Science Advances (2025) doi:10.1126/sciadv.adv7790.

14. Fiuza, U. M. et al. Morphogenetic constraints in the development of gastruloids: Implications for mouse gastrulation. Cells & development 183, (2025).

15. Raj, A. & van Oudenaarden, A. Nature, nurture, or chance: stochastic gene expression and its consequences. Cell 135, 216–226 (2008).

16. Zernicka-Goetz, M., Morris, S. A. & Bruce, A. W. Making a firm decision: multifaceted regulation of cell fate in the early mouse embryo. Nat Rev Genet 10, 467–477 (2009).

17. Waddington, C. H. CANALIZATION OF DEVELOPMENT AND THE INHERITANCE OF ACQUIRED CHARACTERS. Nature 150, 563–565 (1942).

18. Rutherford, S. L. & Lindquist, S. Hsp90 as a capacitor for morphological evolution. Nature 396, 336–342 (1998).

19. Raj, A., Rifkin, S. A., Andersen, E. & van Oudenaarden, A. Variability in gene expression underlies incomplete penetrance. Nature 463, 913–918 (2010).

20. Torres-Padilla, M.-E. & Chambers, I. Transcription factor heterogeneity in pluripotent stem cells: a stochastic advantage. Development 141, 2173–2181 (2014).

21. Sendra, M. et al. Epigenetic priming of embryonic lineages in the mammalian epiblast. bioRxiv 2024.01.11.575188 (2024) doi:10.1101/2024.01.11.575188.

22. Marks, H. et al. The transcriptional and epigenomic foundations of ground state pluripotency. Cell 149, 590–604 (2012).

23. Klein, A. M. et al. Droplet barcoding for single-cell transcriptomics applied to embryonic stem cells. Cell 161, 1187–1201 (2015).

24. Kalmar, T. et al. Regulated fluctuations in nanog expression mediate cell fate decisions in embryonic stem cells. PLoS Biol 7, e1000149 (2009).

25. Lineages of embryonic stem cells show non-Markovian state transitions. iScience 24, 102879 (2021).

26. McNamara, H. M., Solley, S. C., Adamson, B., Chan, M. M. & Toettcher, J. E. Recording morphogen signals reveals mechanisms underlying gastruloid symmetry breaking. Nat Cell Biol 26, 1832–1844 (2024).

27. Braccioli, L. et al. Identifying cross-lineage dependencies of cell-type-specific regulators in mouse gastruloids. Dev Cell (2025) doi:10.1016/j.devcel.2025.02.013.

28. Canham, M. A., Sharov, A. A., Ko, M. S. H. & Brickman, J. M. Functional heterogeneity of embryonic stem cells revealed through translational amplification of an early endodermal transcript. PLoS Biol. 8, e1000379 (2010).

29. Kolodziejczyk, A. A. et al. Single Cell RNA-Sequencing of Pluripotent States Unlocks Modular Transcriptional Variation. Cell Stem Cell 17, 471–485 (2015).

30. Messmer, T. et al. Transcriptional Heterogeneity in Naive and Primed Human Pluripotent Stem Cells at Single-Cell Resolution. Cell Rep 26, 815–824.e4 (2019).

31. Nair, G., Abranches, E., Guedes, A. M. V., Henrique, D. & Raj, A. Heterogeneous lineage marker expression in naive embryonic stem cells is mostly due to spontaneous differentiation. Sci Rep 5, 13339 (2015).

32. Abranches, E. et al. Stochastic NANOG fluctuations allow mouse embryonic stem cells to explore pluripotency. Development 141, 2770–2779 (2014).

33. Merle, M., Friedman, L., Chureau, C., Shoushtarizadeh, A. & Gregor, T. Precise and scalable self-organization in mammalian pseudo-embryos. Nat. Struct. Mol. Biol. (2024) doi:10.1038/s41594-024-01251-4.

34. Cermola, F. et al. Gastruloid Development Competence Discriminates Different States of Pluripotency. Stem Cell Reports 16, 354–369 (2021).

35. Moris, N. et al. An in vitro model of early anteroposterior organization during human development. Nature 582, 410–415 (2020).

36. de Jong, M. A. et al. The shapes of elongating gastruloids are consistent with convergent extension driven by a combination of active cell crawling and differential adhesion. PLoS Comput. Biol. 20, e1011825 (2024).

37. Elowitz, M. B., Levine, A. J., Siggia, E. D. & Swain, P. S. Stochastic Gene Expression in a Single Cell. Science (2002) doi:10.1126/science.1070919.

38. Süel, G. M., Kulkarni, R. P., Dworkin, J., Garcia-Ojalvo, J. & Elowitz, M. B. Tunability and noise dependence in differentiation dynamics. Science (New York, N.Y.) 315, (2007).

39. Ricardo, D. On the Principles of Political Economy and Taxation. (1817).

40. Triandafillou, C. G., Sompalle, P., Heyman, Y. & Raj, A. Single-cell spatial mapping reveals reproducible cell type organization and spatially-dependent gene expression in gastruloids. bioRxiv: the preprint server for biology (2025) doi:10.1101/2025.07.14.664617.

41. Eng, C.-H. L. et al. Transcriptome-scale super-resolved imaging in tissues by RNA seqFISH+. Nature 568, 235–239 (2019).

42. Rosen, L. U., et al. Inter-gastruloid heterogeneity revealed by single cell transcriptomics time course: implications for organoid based perturbation studies. bioRxiv 2022.09.27.509783 (2022) doi:10.1101/2022.09.27.509783.

43. Stavish, D. et al. Generation and trapping of a mesoderm biased state of human pluripotency. Nature Communications 11, 1–16 (2020).

44. Pijuan-Sala, B. et al. A single-cell molecular map of mouse gastrulation and early organogenesis. Nature 566, 490–495 (2019).

45. Zhou, X. et al. Hes1 desynchronizes differentiation of pluripotent cells by modulating STAT3 activity. Stem Cells 31, 1511–1522 (2013).

46. Kobayashi, T. et al. The cyclic gene Hes1 contributes to diverse differentiation responses of embryonic stem cells. Genes Dev 23, 1870–1875 (2009).

47. Gery, S. et al. Retinoic acid regulates C/EBP homologous protein expression (CHOP), which negatively regulates myeloid target genes. Blood 104, 3911–3917 (2004).

48. Hu, Z.-L. et al. The role of the transcription factor Rbpj in the development of dorsal root ganglia. Neural Dev 6, 14 (2011).

49. Souilhol, C. et al. NOTCH activation interferes with cell fate specification in the gastrulating mouse embryo. Development 142, 3649–3660 (2015).

50. Kuwahara, A. et al. Tcf3 represses Wnt-β-catenin signaling and maintains neural stem cell population during neocortical development. PLoS One 9, e94408 (2014).

51. Kim, C. H. et al. Repressor activity of Headless/Tcf3 is essential for vertebrate head formation. Nature 407, 913–916 (2000).

52. Website. https://www.biorxiv.org/content/10.1101/2024.12.12.628151v2.full.

53. Latres, E., Chiaur, D. S. & Pagano, M. The human F box protein beta-Trcp associates with the Cul1/Skp1 complex and regulates the stability of beta-catenin. Oncogene 18, 849–854 (1999).

54. Amel, A., Rabeling, A., Rossouw, S. & Goolam, M. Wnt and BMP signalling direct anterior–posterior differentiation in aggregates of mouse embryonic stem cells. Biol Open 12, bio059981 (2023).

55. Hennessy, M. J., Fulton, T., Turner, D. A. & Steventon, B. Negative feedback on Retinoic Acid by Brachyury guides gastruloid symmetry-breaking. bioRxiv 2023.06.02.543388 (2023) doi:10.1101/2023.06.02.543388.

56. Dias, A. et al. Opposing Nodal and Wnt signalling activities govern the emergence of the mammalian body plan. bioRxiv 2025.01.11.632562 (2025) doi:10.1101/2025.01.11.632562.

57. Martin, B. L. & Kimelman, D. Brachyury establishes the embryonic mesodermal progenitor niche. Genes & Development 24, 2778 (2010).

58. Underhill, E. J. & Toettcher, J. E. Control of gastruloid patterning and morphogenesis by the Erk and Akt signaling pathways. Development 150, (2023).

59. Turner, D. A. et al. Anteroposterior polarity and elongation in the absence of extra-embryonic tissues and of spatially localised signalling in gastruloids: mammalian embryonic organoids. Development 144, 3894–3906 (2017).

60. Suppinger, S. et al. Multimodal characterization of murine gastruloid development. Cell Stem Cell 30, 867–884.e11 (2023).

61. Weber, T. S. et al. LoxCode in vivo barcoding reveals epiblast clonal fate bias to fetal organs. Cell (2025) doi:10.1016/j.cell.2025.04.026.

62. Frenster, J. D. et al. A mosaic gastruloid model highlights the developmental stage-specific restriction of cell competition in mammalian pre-gastrulation. bioRxiv 2025.01.25.634859 (2025) doi:10.1101/2025.01.25.634859.

63. Regalado, S. G. et al. Barcoded monoclonal embryoids are a potential solution to confounding bottlenecks in mosaic organoid screens. bioRxiv 2025.05.23.655669 (2025) doi:10.1101/2025.05.23.655669.

64. Bowling, S. et al. P53 and mTOR signalling determine fitness selection through cell competition during early mouse embryonic development. Nat Commun 9, 1763 (2018).

65. Blastocyst complementation-based rat-derived heart generation reveals cardiac anomaly barriers to interspecies chimera development. iScience 27, 111414 (2024).

66. Ruiz-Estevez, M. et al. Liver development is restored by blastocyst complementation of HHEX knockout in mice and pigs. Stem Cell Research & Therapy 12, 1–13 (2021).

67. Esmaeili, A., Eteghadi, A., Landi, F. S., Yavari, S. F. & Taghipour, N. Recent approaches in regenerative medicine in the fight against neurodegenerative disease. Brain Res 1825, 148688 (2024).

68. Mao, J. et al. Reprogramming stem cells in regenerative medicine. Smart Medicine 1, e20220005 (2022).

69. Moriwaki, T. et al. Scalable production of homogeneous cardiac organoids derived from human pluripotent stem cells. Cell Rep Methods 3, 100666 (2023).

70. Bolondi, A. et al. Reconstructing axial progenitor field dynamics in mouse stem cell-derived embryoids. Dev. Cell (2024) doi:10.1016/j.devcel.2024.03.024.

71. Wehmeyer, A. E. et al. Competing regulatory modules control the transition between mammalian gastrulation modes. bioRxiv 2025.05.07.652670 (2025) doi:10.1101/2025.05.07.652670.

72. Cai, D., Cohen, K. B., Luo, T., Lichtman, J. W. & Sanes, J. R. Improved tools for the Brainbow toolbox. Nat. Methods 10, 540–547 (2013).

73. Stringer, C. & Pachitariu, M. Cellpose3: one-click image restoration for improved cellular segmentation. Nat Methods 22, 592–599 (2025).

74. Stringer, C., Wang, T., Michaelos, M. & Pachitariu, M. Cellpose: a generalist algorithm for cellular segmentation. Nat Methods 18, 100–106 (2021).

75. Comparable Generation of Activin-Induced Definitive Endoderm via Additive Wnt or BMP Signaling in Absence of Serum. Stem Cell Reports 3, 5–14 (2014).

76. Corces, M. R. et al. An improved ATAC-seq protocol reduces background and enables interrogation of frozen tissues. Nat Methods 14, 959–962 (2017).

77. Wang, Q., et al. Exploring Epigenomic Datasets by ChIPseeker. Curr Protoc 2, e585 (2022).

78. Heinz, S. et al. Simple combinations of lineage-determining transcription factors prime cis-regulatory elements required for macrophage and B cell identities. Mol. Cell 38, 576–589 (2010).

79. van der Walt, S. et al. scikit-image: image processing in Python. PeerJ 2, e453 (2014).

