## Supplemental Data for "Gastruloid patterning reflects division of labor among biased stem cell clones"

#### Supplemental Information

Supplemental Figures 1-24, Supplemental Movies 1-2, Supplemental Table 1

#### Supplemental Figure 1

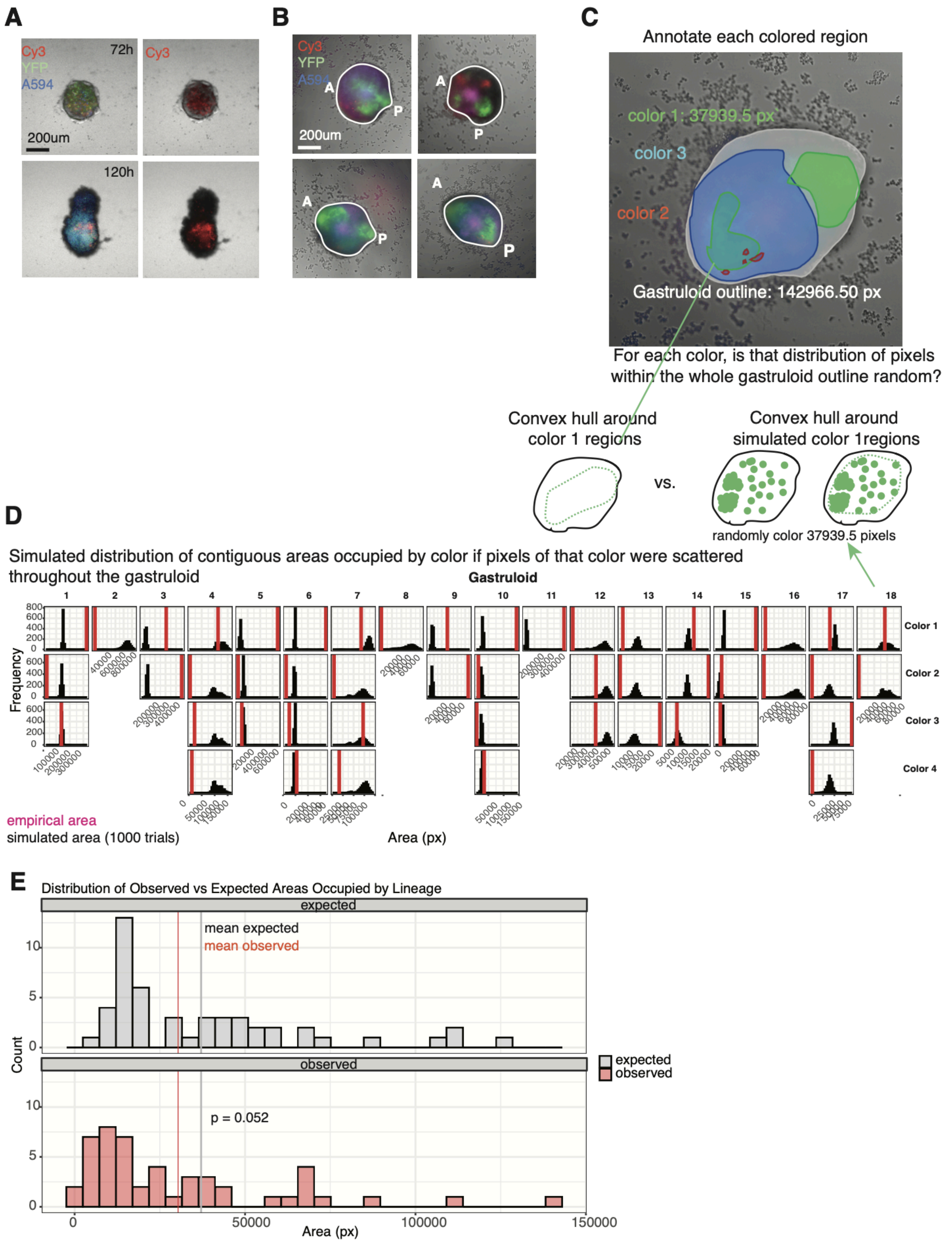

**Supplemental Figure 1. Additional evidence for clones' propensity in polyclonal gastruloids.**
**(A)** Stills from a time-lapse of gastruloid elongation in a polyclonal gastruloid comprising cells each expressing a unique combination of eGFP, mKate2, and mOrange2. Panels on the right highlight cells visible in the Cy3 channel.
**(B)** Representative polyclonal gastruloids.
**(C)** Annotations of regions within a representative gastruloid corresponding to a given color. The whole gastruloid is outlined, and the number of pixels contained within the gastruloid is labeled, as is the number of pixels annotated as belonging to color 1. The convex hull around regions of each color was compared to the convex hull around pixels of that color if randomly scattered throughout the gastruloid.
**(D)** Histograms corresponding to the area occupied by each annotation color for simulated gastruloids. A vertical red line is drawn to represent the empirical area of the convex hull corresponding to that annotation.
**(E)** Histograms comparing the mean area across 1000 simulations described in D with the mean empirical area occupied by each color across 18 gastruloids. Statistical analysis: Kolmogorov-Smirnov test.

To test whether the distribution of colored lineages within polyclonal gastruloids (**Supplemental Figure 1A**, **B**) was random, we drew annotations around contiguous regions that had the same color (**Supplemental Figure 1C**). We assume that each color corresponds to a clonal lineage. That said, in theory, multiple lineages can express combinations of fluorescent proteins that result in the same color by eye. For each color annotation within each gastruloid, the area of the convex hull around the annotations was computed. Next, we took all of the pixels belonging to a given annotation and performed 1000 simulations where these pixels were randomly scattered within the polygon annotation of the gastruloid outline. The area of the convex hull around these pixels was calculated (**Supplemental Figure 1D**). We compared the empirical distribution of (observed) convex-hull areas (across all colors across all gastruloids) with the distribution of means (expected) of each simulated distribution (**Supplemental Figure 1D**). The mean of the observed distribution was less than the mean of the expected distribution; the p-value corresponding to the difference between distributions was 0.052 by a two-sided KS test. A caveat to this analysis is that lineages/clones were manually annotated according to their color. It is possible that, for example, two different annotations annotated as belonging to the same clone actually belong to different clones that have similar fluorescent protein expression and thus could not be discerned by eye. Assigning regions to a single clone when they belong to two clones would result in a larger convex hull area around the two annotations, as opposed to two separate, smaller annotations labelling different clones.

#### Supplemental Figure 2

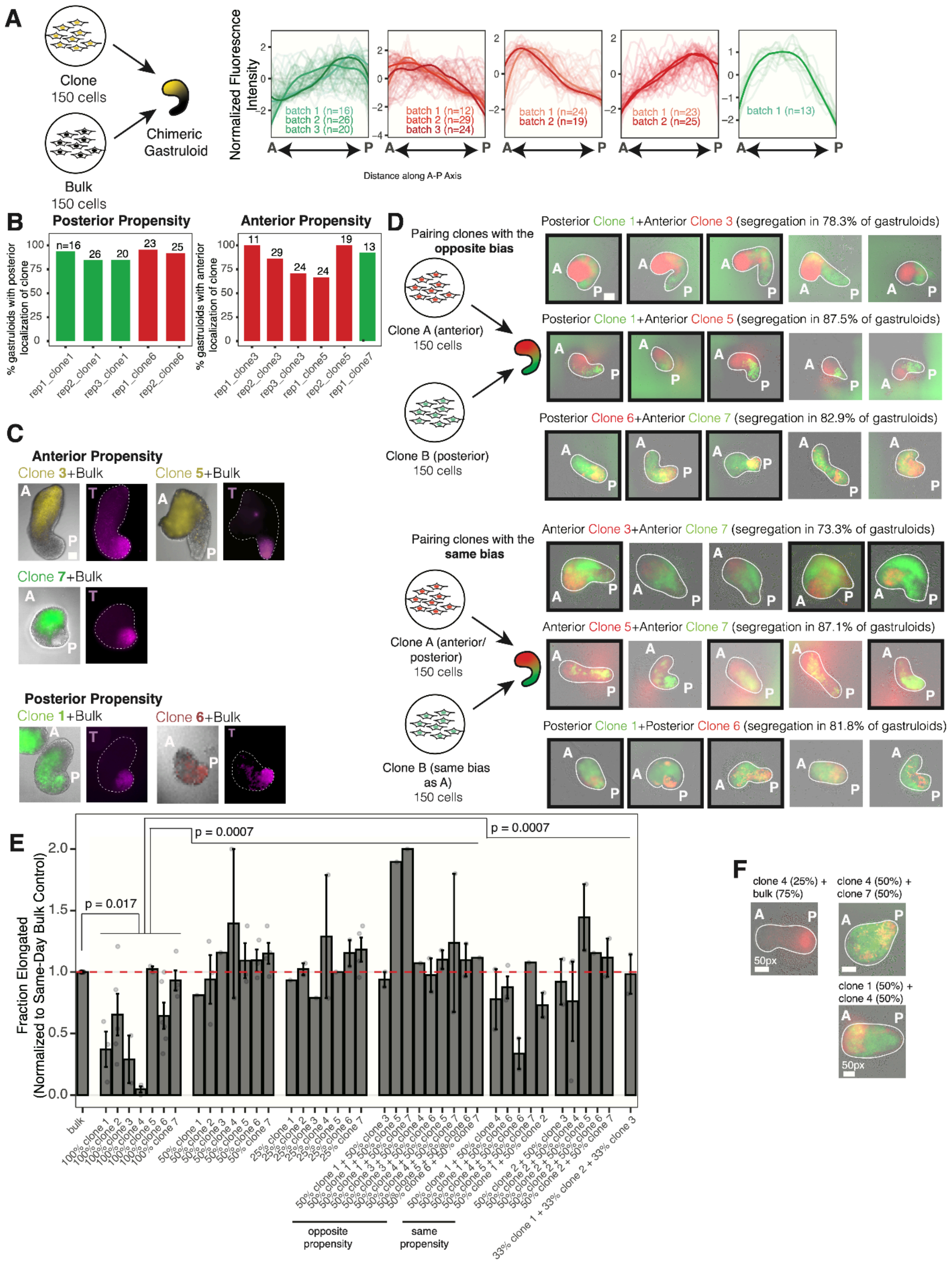

**Supplemental Figure 2. Relative propensities among clones.**
**(A)** Individual fluorescence intensity curves for each chimeric gastruloid comprising 50% clone and 50% bulk. Plotted for each gastruloid is the z-score of its fluorescence intensity along the A-P axis.
**(B)** Bar graphs showing, for each 50% clone/50% bulk combination, the fraction of gastruloids. with fluorescence intensity in the anterior ⅓ of the gastruloid exceeding that in the posterior ⅓ of the gastruloid or with fluorescence intensity in the posterior ⅓ of the gastruloid exceeding that in the anterior ⅓ of the gastruloid. The plot is faceted by propensity.
**(C)** Images showing 50% clone/50% bulk gastruloids along with corresponding immunofluorescence staining for T. The scale bar represents 100μm.
**(D)** Five representative images for each combination of clones. Images shown in **Figure 1E** are boxed with a black outline. The scale bar represents 100px. The percentage of gastruloids in which clones occupy distinct spatial domains along the A-P axis is listed.
**(E)** Bargraph showing the normalized fraction of elongated gastruloids for pure clone gastruloids, gastruloids made from a combination of a pure clone and bulk, and gastruloids made from a combination of clones. Normalization was performed by dividing the fraction of elongated gastruloids for a given condition within a batch by the fraction of elongated gastruloids among bulk gastruloids within that batch. Bars for clone-clone pairings are arranged according to whether clones in the pair have the same or opposite propensity. Since we could not confidently assign a propensity to clone 2, bars corresponding to clone combinations including clone 2 are grouped separately
**(F)** Representative images of chimeric gastruloids including clone 4. The scale bar represents 50px.

**Supplemental Figure 3**

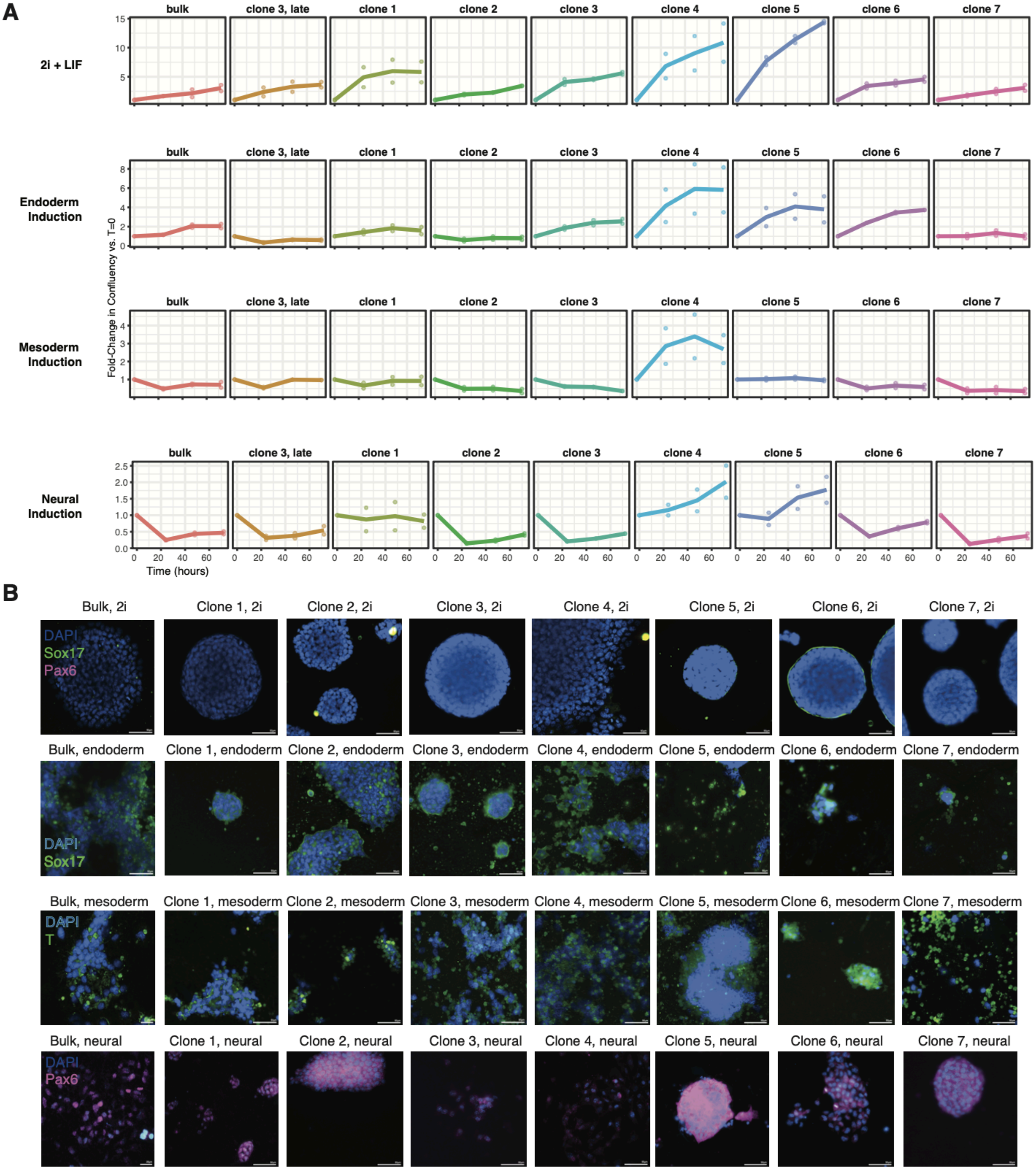


**Supplemental Figure 3. Response of clones to culture conditions intended to induce differentiation to endoderm, mesoderm, or neural cell types.**
**(A)** 72-hour growth curves for all clones in 2i+LIF, or endoderm, mesoderm, or neural induction medias.
**(B)** Immunofluorescence for T, Sox17, or Pax6.

As shown in **Figure 4B**, mES cell clones did not show significant differences in their expression of pluripotency genes from bulk mES cells. We were curious whether these cells could still differentiate into different germ layer identities. We trypsinized cells and plated them in 24-well plates under pluripotency-promoting conditions, endoderm-induction conditions, mesoderm-induction conditions, or neural-induction conditions, as detailed in **Methods**. We measured confluency of cells over 72 hours using an Incucyte. At the end of 72 hours, samples were fixed, and immunofluorescence for T, Sox17, and Pax6 was performed. We tested all 7 clones, bulk mES cells, and a late-passage sample from clone 3.

Clones varied in their growth under different culture conditions. For example, clone 5 had the fastest growth rate in 2i+LIF media, followed by clone 4, which generally had the fastest growth rates overall. However, growth rates in differentiation media did not correspond to gastruloid formation ability, since clone 4 rarely formed elongated gastruloids (**Figure 1B**).

Immunofluorescence showed that, under 2i+LIF media, clones had a dome-shaped colony morphology, which we expect. There was no expression of germ layer markers under these conditions. These markers were subsequently induced in their respective differentiation conditions. Thus, clones vary in their tolerance of different differentiation induction conditions but are capable of being induced to express markers of their differentiated fates in these conditions.

#### Supplemental Figure 4

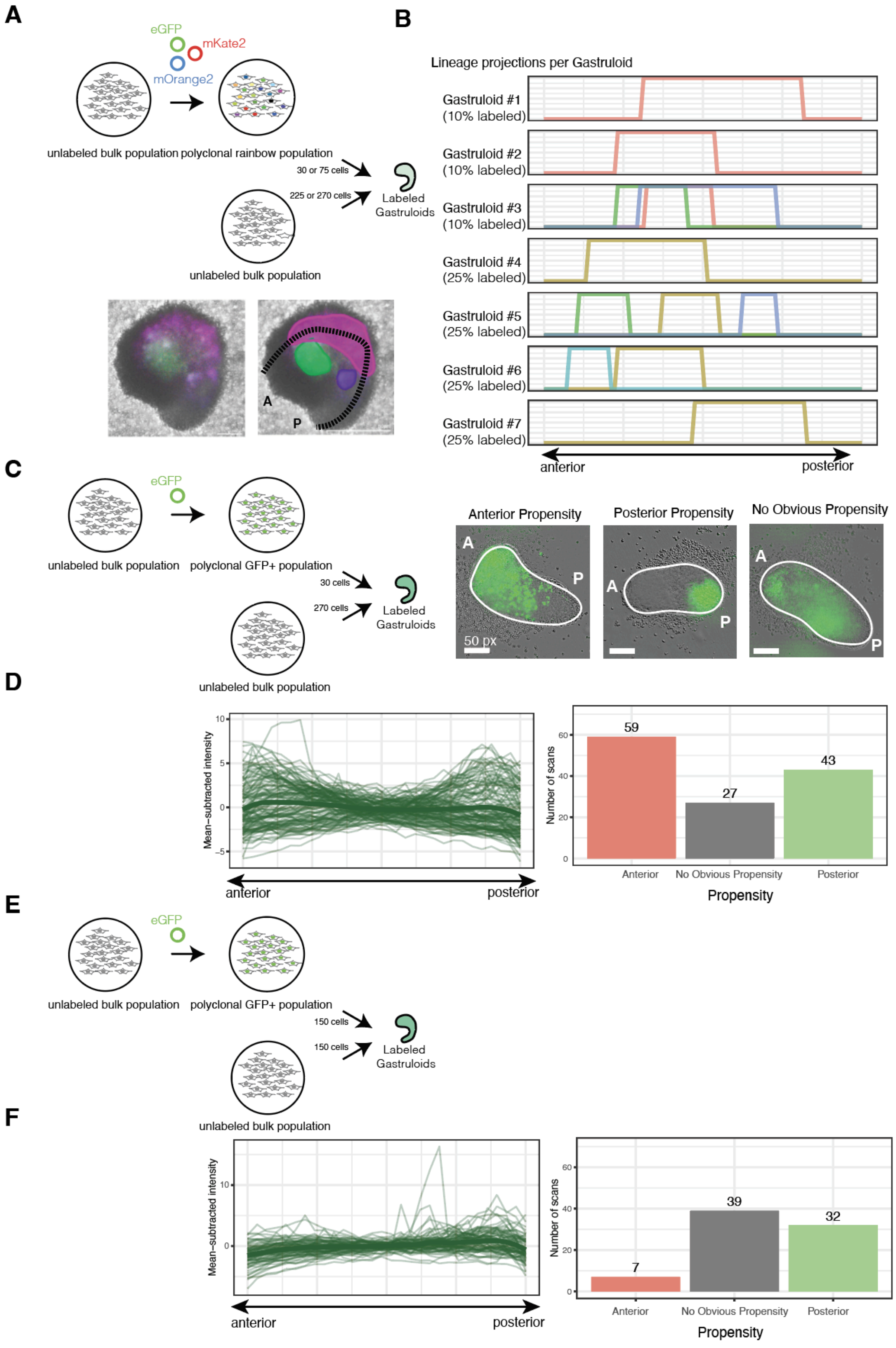


**Supplemental Figure 4. Propensities within a polyclonal population.**

**(A)** An unlabeled bulk mES cell population was transduced at high MOI with plasmids encoding eGFP, mOrange2, and mKate2. Cells from the resulting multicolor population were aggregated alongside unlabeled mES cells in defined proportions. Color-labeled regions from the resulting 120-hour gastruloids were annotated and projected onto the gastruloid A-P axis.

**(B)** The regions along the A-P axis occupied by each colored population are shown.

**(C)** An unlabeled bulk mES cell population was transduced with a plasmid encoding eGFP. Cells from the resulting polyclonal GFP+ population were aggregated alongside unlabeled mES cells such that 10% of cells in each aggregate were from the GFP+ population.
**(D)** Color-labeled regions from the resulting 120-hour gastruloids were annotated and projected onto the gastruloid A-P axis. Mean-subtracted green fluorescence intensities along the A-P axis are plotted for each gastruloid, as are the numbers of gastruloids where green cells demonstrated an “anterior” propensity (anterior ⅓ of the gastruloid had higher fluorescence intensity than the posterior ⅓ above a threshold set to 1.5 units), “posterior” propensity (posterior ⅓ of the gastruloid had higher fluorescence intensity than the anterior ⅓), or neither.
**(E)** Similarly, gastruloids were generated such that 50% of cells in each aggregate were from the polyclonal GFP+ population described above, while the remaining 50% were from an unlabeled polyclonal population.
**(F)** Mean-subtracted green fluorescence intensities along the A-P axis are plotted for each gastruloid, as are the numbers of gastruloids where green cells demonstrated an “anterior” propensity (anterior ⅓ of the gastruloid had higher fluorescence intensity than the posterior ⅓ above a threshold set to 1.5 units), “posterior” propensity (posterior ⅓ of the gastruloid had higher fluorescence intensity than the anterior ⅓), or neither.

We needed to establish how many clones are typically represented when labeled cells are mixed into an unlabeled bulk population. We first generated chimeric aggregates, including either 225 (75%) or 270 (90%) cells from an unlabeled bulk population and, respectively, 75 (25%) or 30 (10%) cells from a polyclonal, multicolor population. We could then estimate how many clones appear in chimeric aggregates comprising both unlabeled and labeled populations, which we could track (Supplemental Figure 3A). We identified between 1 and 3 distinctly-colored populations within 120-hr chimeric gastruloids (**Supplemental Figure 4A-B**).

We then tested whether this small labeled population showed consistent spatial bias. We transduced a polyclonal population to express GFP and generated chimeric aggregates where 10% of the cells were from this GFP+ population and compared GFP fluorescence intensity between the anterior or posterior of gastruloids that elongated after 120 hours (**Supplemental Figure 4C**).

59/129 elongated gastruloids included GFP-labeled cells in the gastruloid anterior, and 43/129 included GFP-labeled cells in the posterior, with the remaining 27 gastruloids including GFP+ cells throughout the anterior-posterior axis (**Supplemental Figure 4D**). Because the labeled fraction represents only 1-3 clones, the fact that most gastruloids showed spatial enrichment in one direction suggests that most clones do harbor a spatial propensity for occupying the gastruloid anterior or posterior. Meanwhile, when we generated chimeric aggregates where the labeled fraction was increased to 50% (**Supplemental Figure 4E**), 50% of cells had no difference in GFP fluorescence intensity between anterior or posterior, and there were overall far more modest differences in fluorescence intensity between the gastruloid anterior or posterior (**Supplemental Figure 4F**), as expected if individual clone propensities are real but are averaged out when many clones are present together.

**Supplemental Figure 5**

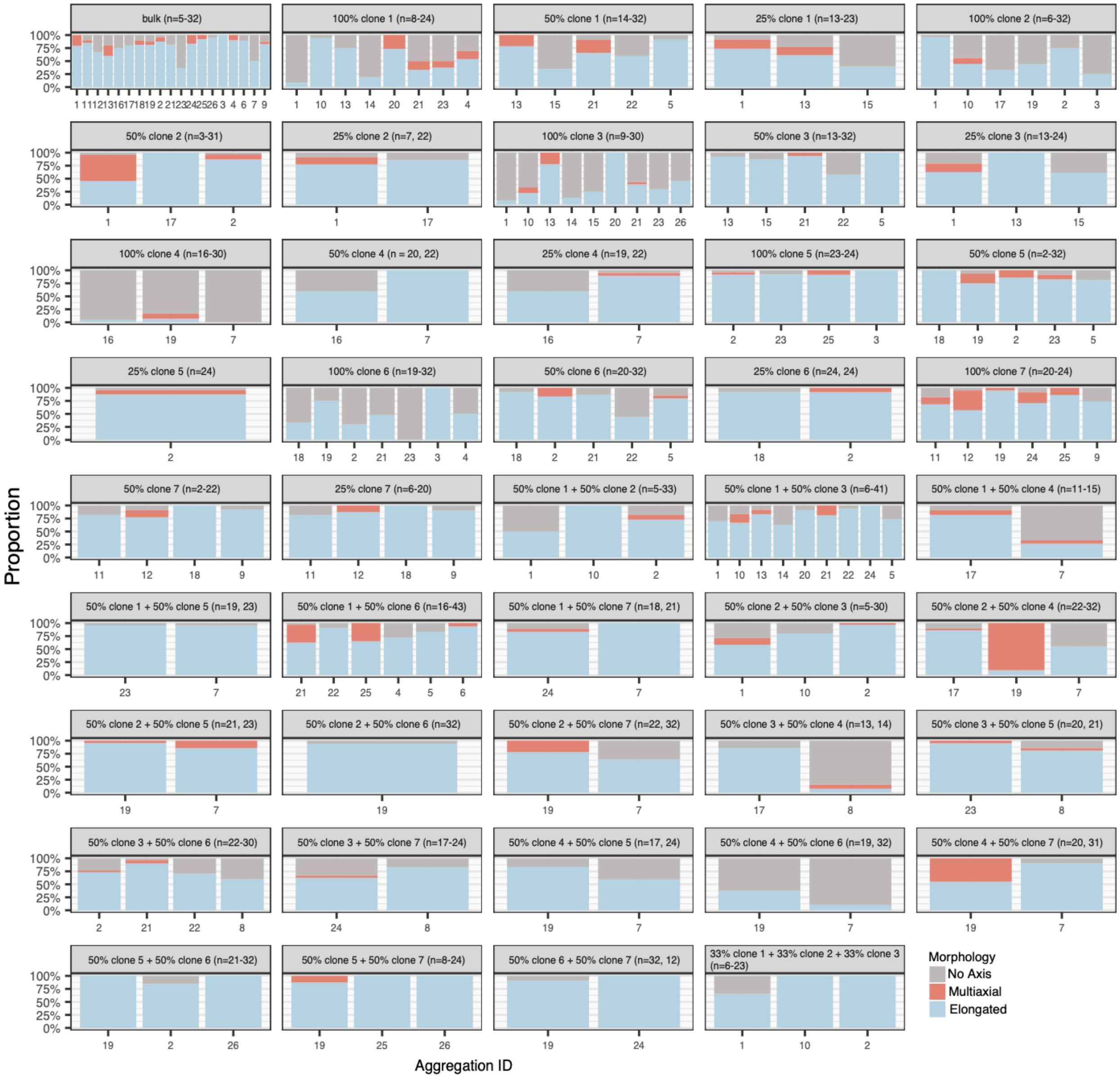


**Supplemental Figure 5.** Proportion of elongated, multiaxial, and no-axis gastruloids for each aggregation batch.

Each facet corresponds to a given condition (bulk, pure clone, clone-bulk combination, clone-clone combination). Stacked bars correspond to the proportions of gastruloids within each batch that are annotated as “elongated,” “multiaxial,” or “no axis.” This data is used to generate bar graphs in Figure 1 and Supplemental Figure 2. Each aggregation ID corresponds to a single round of aggregation; gastruloids aggregated on the same day share an aggregation ID number.

**Supplemental Figure 6**

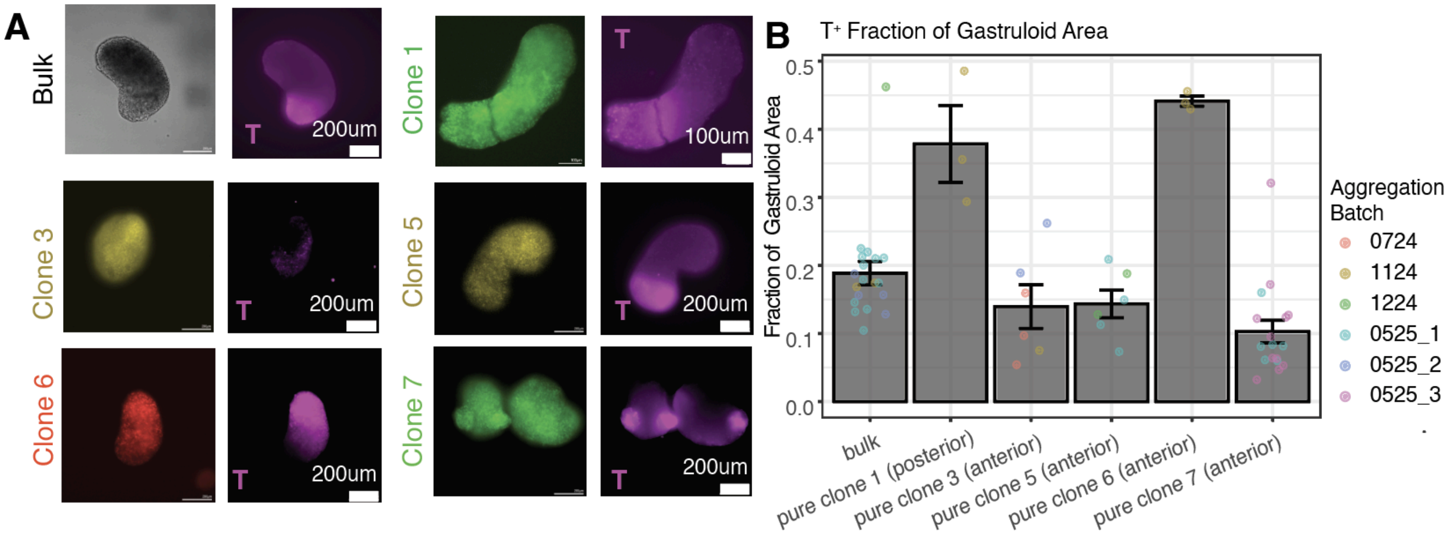


**Supplemental Figure 6. Immunofluorescence for T in gastruloids comprising a pure clone.**
**(A)** Images corresponding to T expression in gastruloids formed from bulk mES cells or a pure clone.
**(B)** Bargraphs representing the fraction of a gastruloid's area that stains positively for T. Each point corresponds to a gastruloid, colored by the batch in which it was aggregated.

**Supplemental Figure 7**

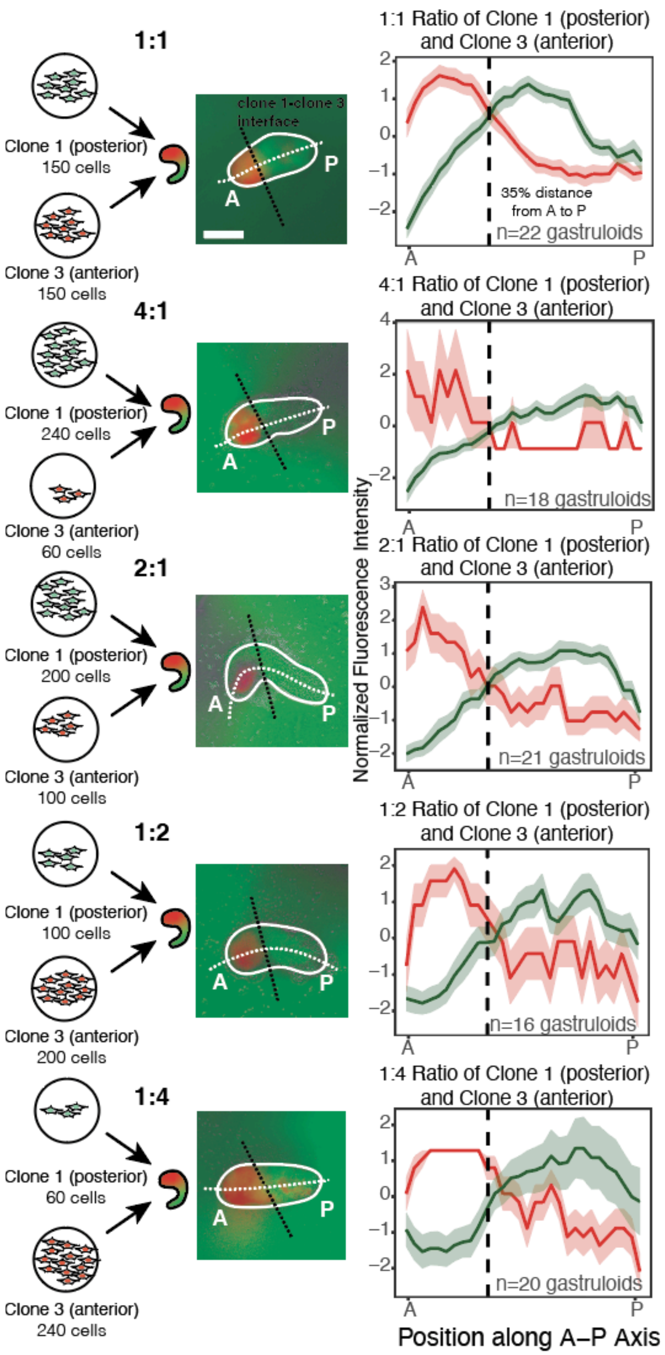


**Supplemental Figure 7. Spatial specialization among mES cell clones during gastruloid patterning.** Aggregation of clone 1 and clone 3 at various ratios. Representative images of gastruloids are shown, along with a white annotation depicting the A-P axis and a black dotted line depicting the interface between each clone. The scale bar corresponds to 100px.

**Supplemental Figure 8**

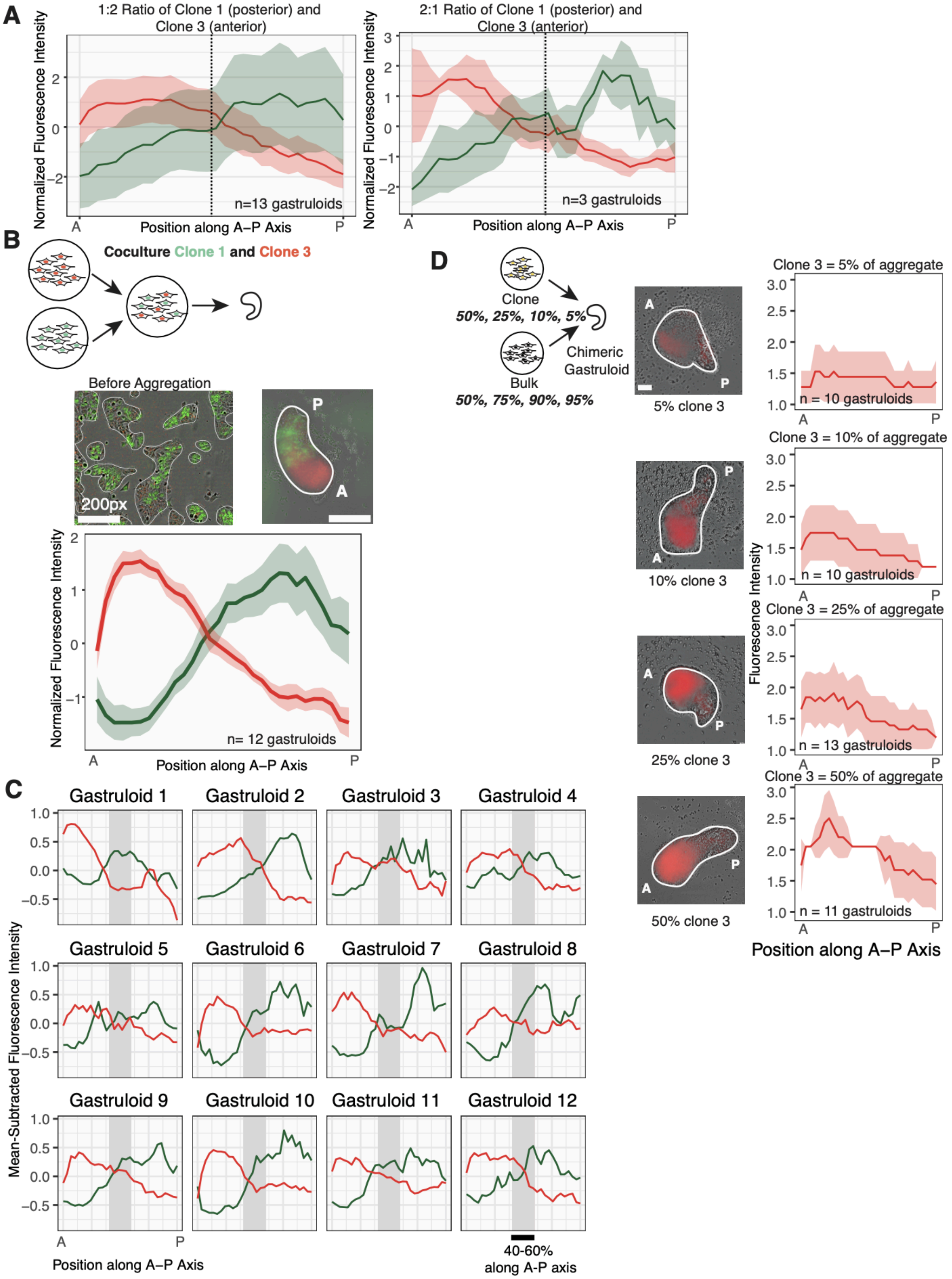


**Supplemental Figure 8. Additional data showing robustness of spatial specialization among mES cell clones during gastruloid patterning.**

**(A)** Additional replicate data for gastruloids formed from clone 1 and clone 3 in a 1:2 ratio and a 2:1 ratio. (*left*) 1:2 ratio of clone 1 to clone 3. (*right*) 2:1 ratio of clone 1 to clone 3. A vertical black line marks 50% of the distance along the A-P axis.
**(B)** Fluorescence intensity curves in red and green channels for gastruloids generated from cocultured clone 1 and clone 3 mES cells. Also shown are representative images of cocultured cells prior to aggregation and of a representative gastruloid made from the cocultured cells. Scale bars correspond to 50px.
**(C)** Individual fluorescence intensity curves for each gastruloid in **(B)**. The average plot is generated by averaging all line scans and then subtracting the mean of the average curve and scaling by standard deviation of values within the curve. Individual line scans simply have the mean subtracted. The green channel is divided by a factor of 10 for appropriate scaling with the red channel. For all gastruloids, the intersection point between curves was 34-56% along the A-P axis.
**(D)** Average fluorescence intensity profiles for gastruloids generated from clone 3 and bulk. Clone 3 comprised 5%, 10%, 25%, or 50% of the initial aggregate as shown. Scale bars correspond to 50px.

Spatial specialization among clones and improved elongation in chimeric gastruloids could, in principle, result from competition, where one clone outcompetes or displaces another during development. We asked whether biased clones could be excluded by a larger, unlabeled population. Our strategy was to form chimeric aggregates containing a fluorescently labeled clone at 5%, 10%, 25%, or 50% of the total cells, with the remainder made up of unlabeled bulk mES cells. If sorting were driven by competitive advantage, the clone might be eliminated from the gastruloid, particularly in the 5% condition (or alternatively might take over the gastruloid, most likely in the 50% condition). Even at the lowest starting fraction (5%), labeled cells still localized robustly to their characteristic domain along the A–P axis (**Supplemental Figure 8D**). Our results do not exclude the possibility that competition occurs during gastruloid development, but competition does not seem to explain propensities that we observed.

### Supplemental Figure 9

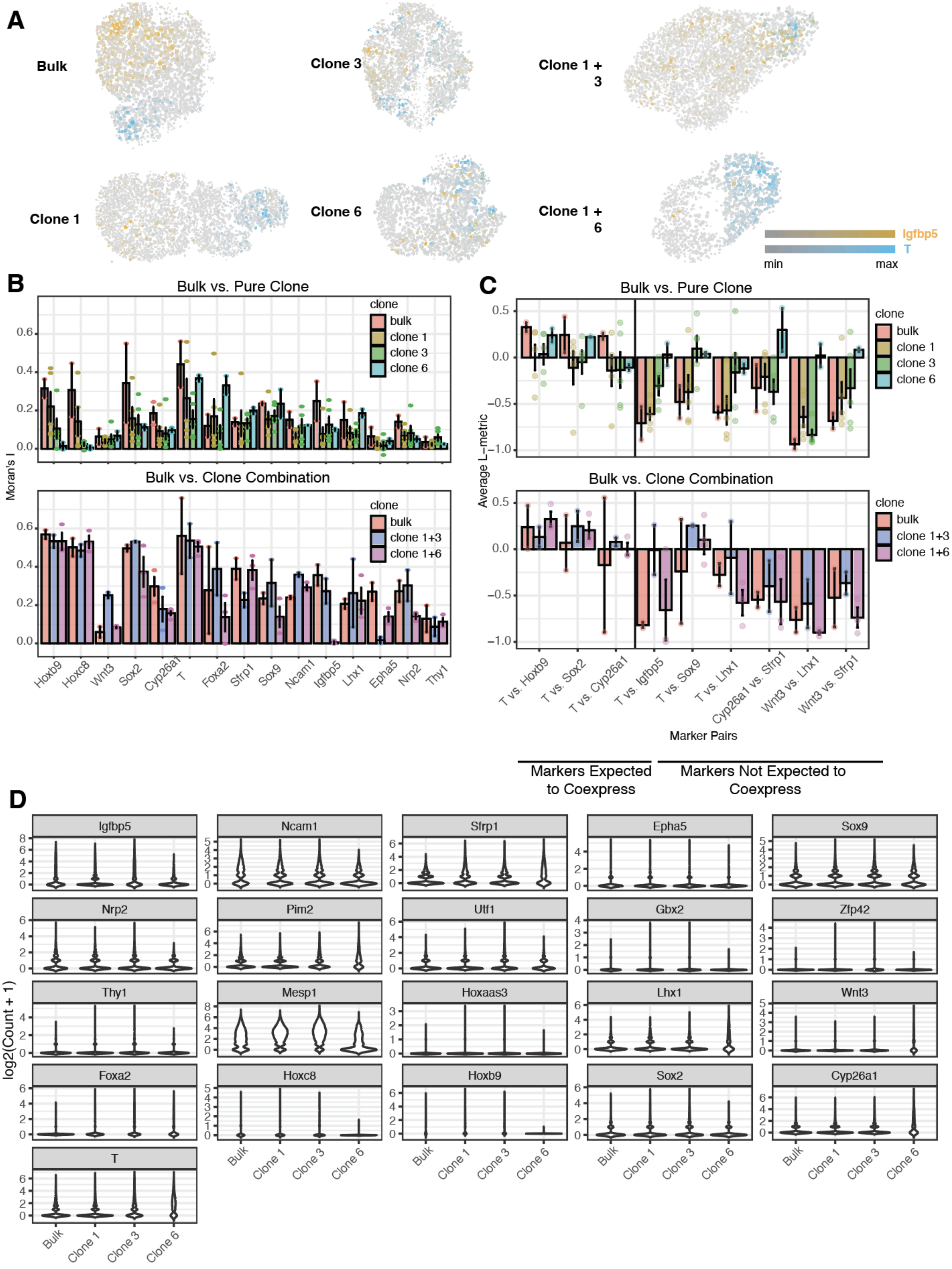

**Supplemental Figure 9. L-metric and Moran's I for additional genes.**
**(A)** Plots showing the spatial distribution of *Igfbp5* and *T* mRNA expression for gastruloids formed from bulk, pure clones, or combinations of clones. Segmentation masks for cells within each gastruloid are shown, colored by their expression of *T* and *Igfbp5*.
**(B)** Bargraphs showing Moran's I for additional genes in our seqFISH panel.
**(C)** Bargraphs showing L-metric values for additional pairs of genes in our seqFISH panel.
**(D)** Violin plots showing expression of genes in our 21-gene panel across whole gastruloids generated from bulk mES cells or pure clones.

To compare the joint distributions of gene expression pairs between bulk and clone gastruloids, we also used a permutation test as described in **Methods**. Gene pairs with different coexpression patterns were, between clone 3 gastruloids and bulk gastruloids: *T* and *Sox2* (p = 0.011198), *T* and *Sox9* (p = 0.011398), *T* and *Lhx1* (p = 0.011798), *T* and *Cyp26a1* (p = 0.012198), *T* and *Hoxb9* (p = 0.012597), *T* and *Igfbp5* (p = 0.04910); and, between clone 6 gastruloids and bulk gastruloids: *T* and *Igfbp5* (p = 0.030994), *T* and *Sox2* (p = 0.031594), *T* and *Lhx1* (p = 0.033993), *T* and *Hoxb9* (p = 0.034793), *T* and *Cyp26a1* (p = 0.035193), *T* and *Sox9* (p = 0.036793), *Wnt3* and *Lhx1* (p = 0.037592), *Cyp26a1* and *Sfrp1* (p = 0.039192), *Wnt3* and *Sfrp1* (0.040792).

**Supplemental Figure 10**

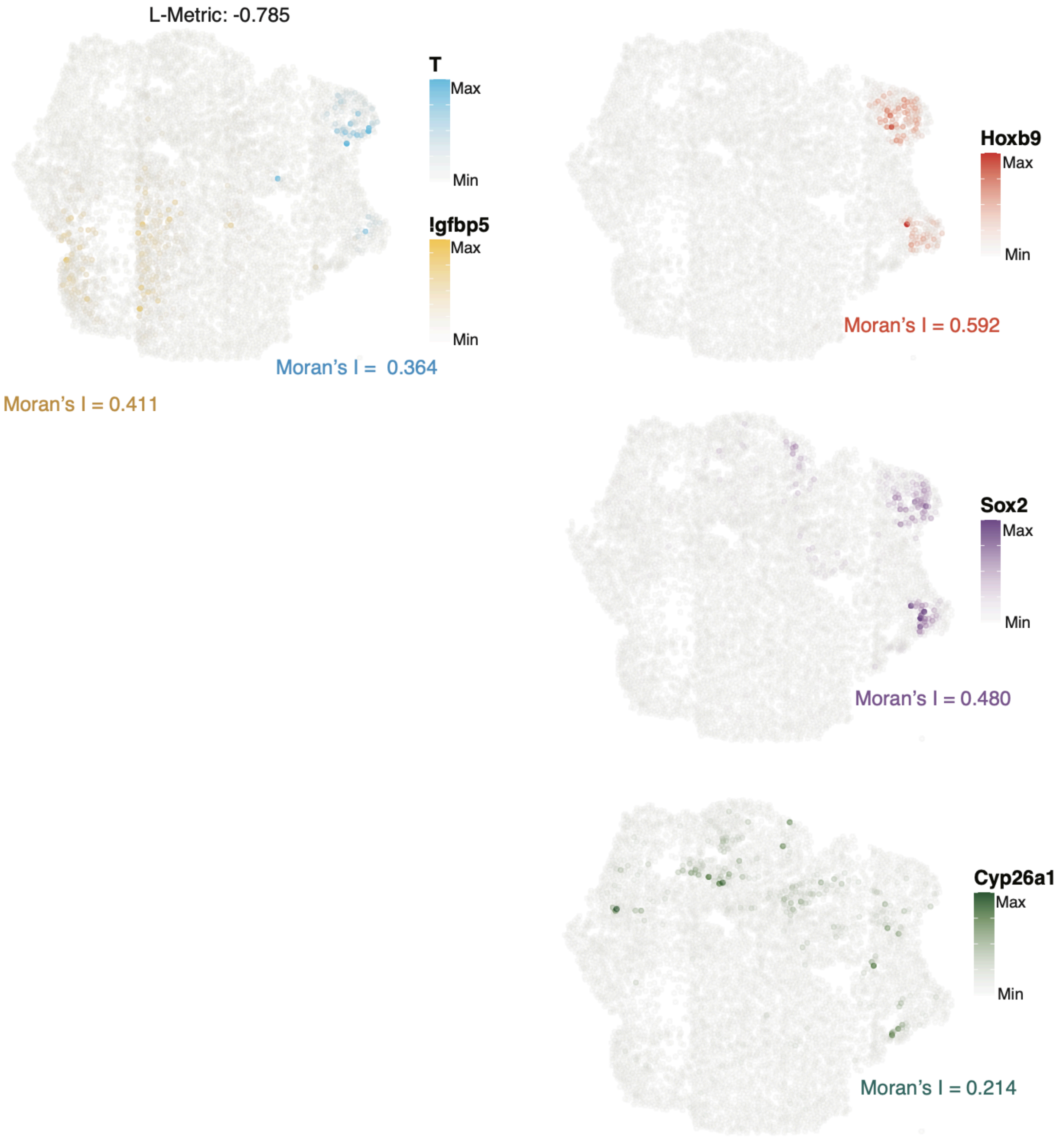


**Supplemental Figure 10. Example of spatial gene expression in a bulk gastruloid that does not elongate along a single axis.**
Spatial gene expression of *T*, *Igfbp5*, *Hoxb9*, *Sox2*, *Cyp26a1* in a bulk gastruloid that was not elongated. Moran's I scores for each gene are shown.

**Supplemental Figure 11**

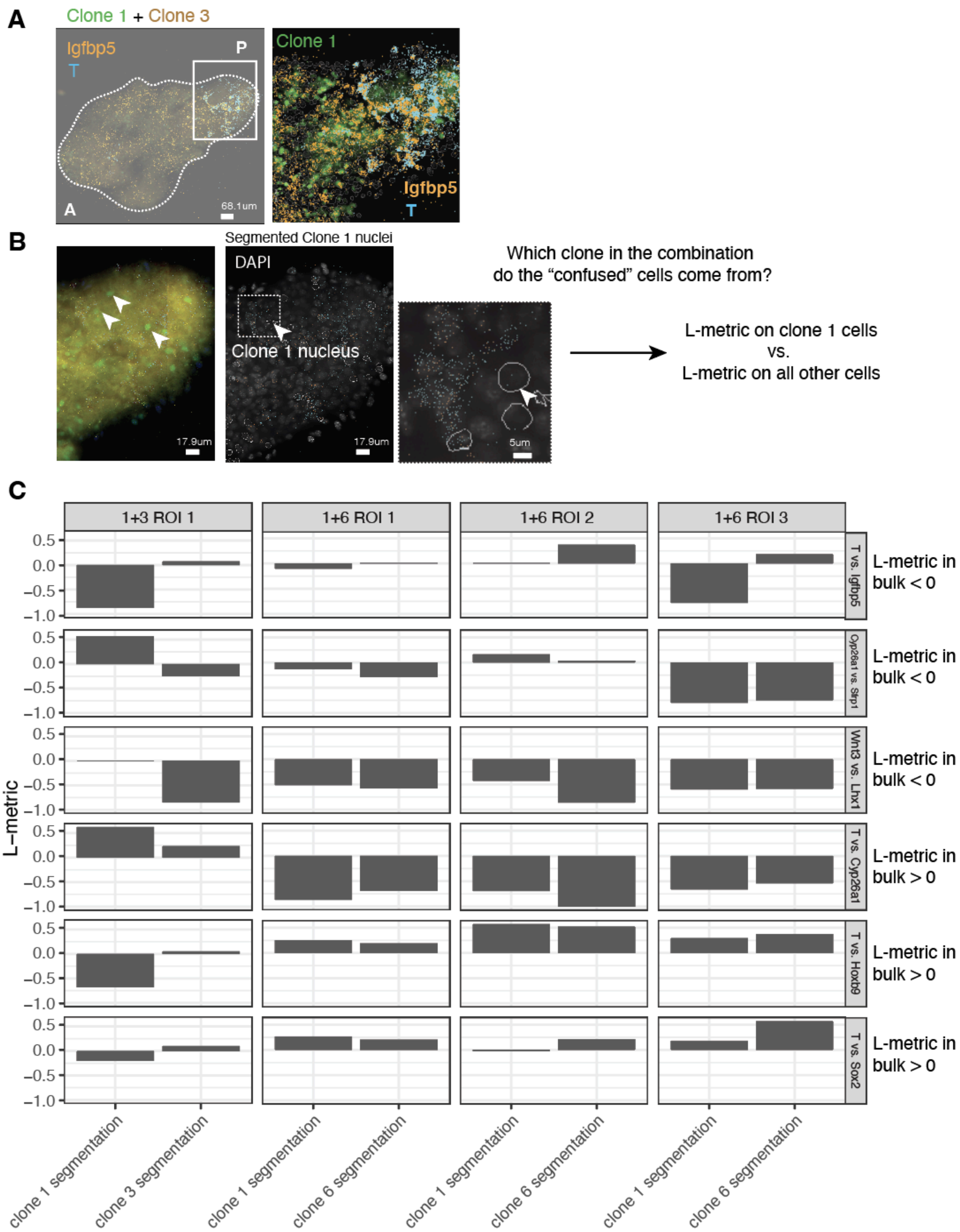


**Supplemental Figure 11. Comparison of L-metric values between clones within individual chimeric gastruloids.**

**(A)** Images from a representative gastruloid. Insets show expression of *T* and *Igfbp5* in the gastruloid posterior. Cells from clone 1 are shown in green.
**(B)** Images from a representative gastruloid showing the visibility of GFP-labeled nuclei from clone 1. Additional insets show segmentation masks surrounding these clone 1 nuclei.
**(C)** L-metric values for pairs of genes. Each column corresponds to an individual gastruloid and each row corresponds to a given gene pair. Bars within each facet correspond to L-metric values for cells within each gastruloid annotated as belonging to clone 1 and all other cells within the gastruloid.

**Supplemental Figure 12**

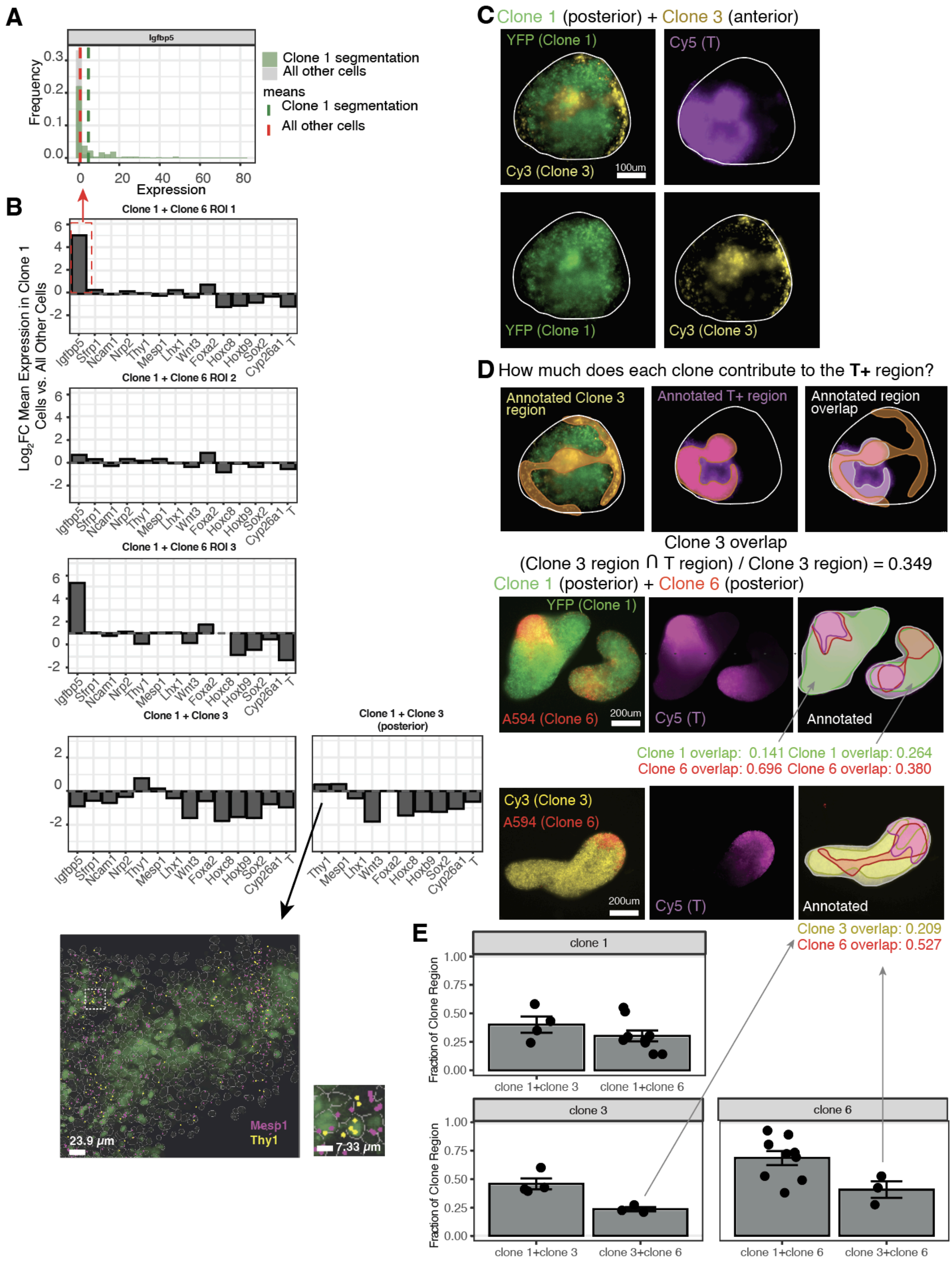

**Supplemental Figure 12. Marker expression of clones within chimeric gastruloids.**

**(A)** Examples of histograms for the expression of *Igfbp5* in clone 1 cells and all other cells are also shown. Vertical lines are drawn to show the mean expression of *Igfbp5* in clone 1 cells and in all other cells.

**(B)** Bar graphs showing the fold-change in mean gene expression between cells segmented as belonging to clone 1 versus all other cells in chimeric gastruloids comprising clone 1 (posterior) and clone 3 (anterior) or clone 1 (posterior) and clone 6 (posterior). Three gastruloids from the combination of clone 1 and clone 6 are shown, and one gastruloid from the combination of clone 1 and clone 3 is shown. Differences in gene expression between clones within the posterior tip (starting from where *T* expression is observed, shown in **Supplemental Figure 11**) of the clone 1+3 gastruloid are also shown. Images of the posterior of the same gastruloid displayed in **Supplemental Figure 11** are shown, with FISH spots corresponding to *Mesp1* and *Thy1* transcripts.

**(C)** A representative gastruloid made from clones 1 and 3. YFP (clone 1) and Cy3 (clone 3) channels are shown, as well as the distribution of T.

**(D)** Annotations showing the spatial distributions of clone 3 and T expression in this representative gastruloid.

**(E)** Representative gastruloid made from clone 1 and clone 6. The distribution of T, and annotations marking each clone and the distribution of T are shown.

**(F)** Bar graphs depicting the overlap of each clone's spatial distribution with the distribution of T expression for chimeric gastruloids within a single batch of aggregation. Facets correspond to each individual clone within a combo, and each bar corresponds to that clone's behavior within a given clone combination.

**Supplemental Figure 13**

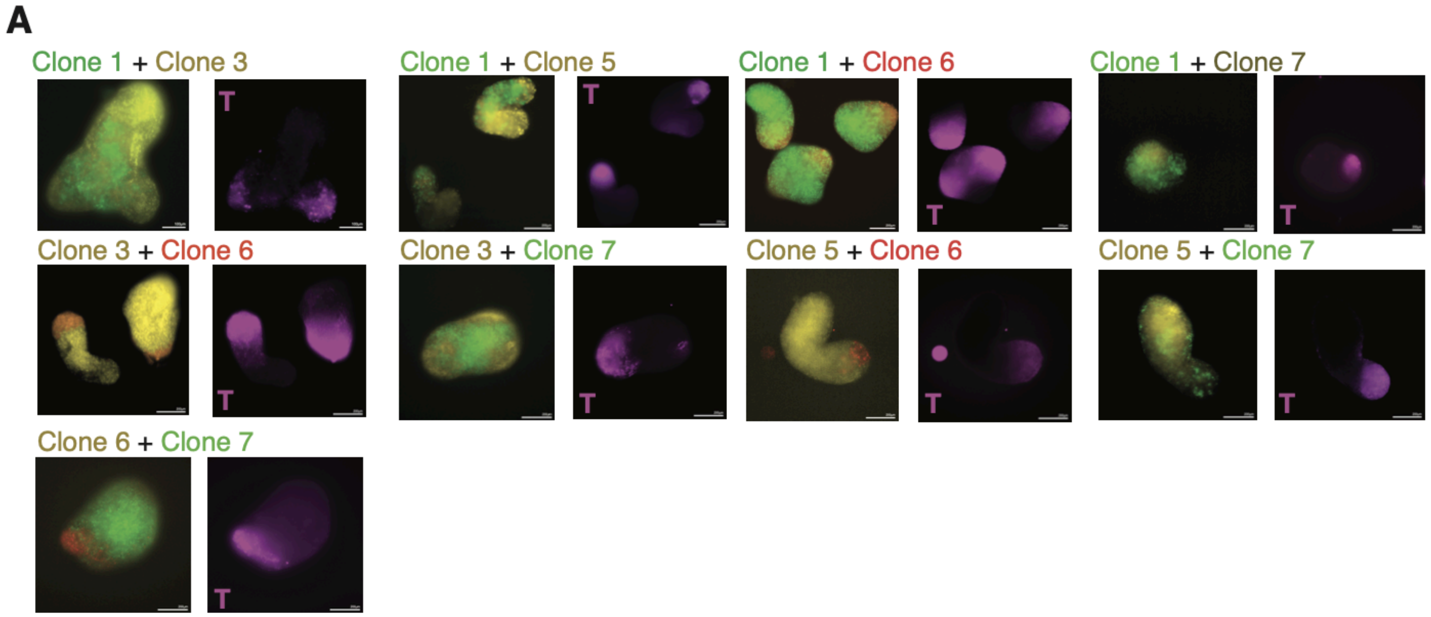


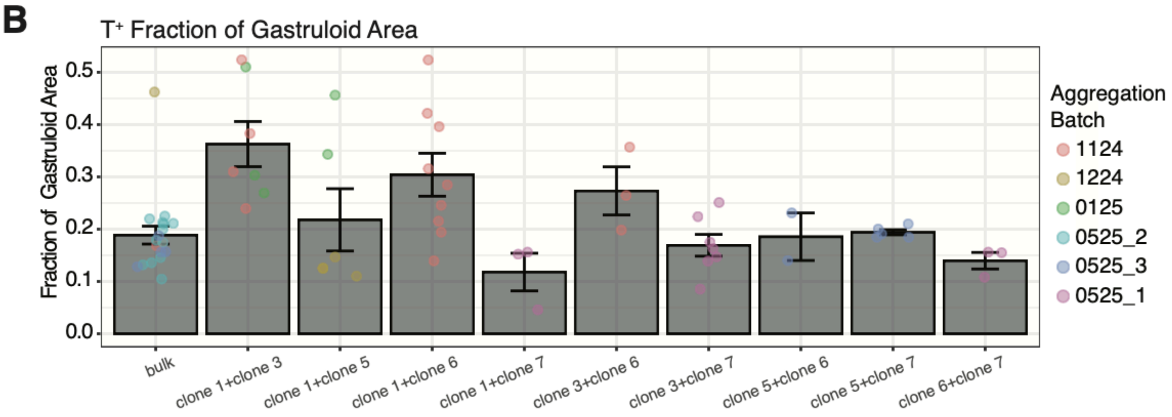


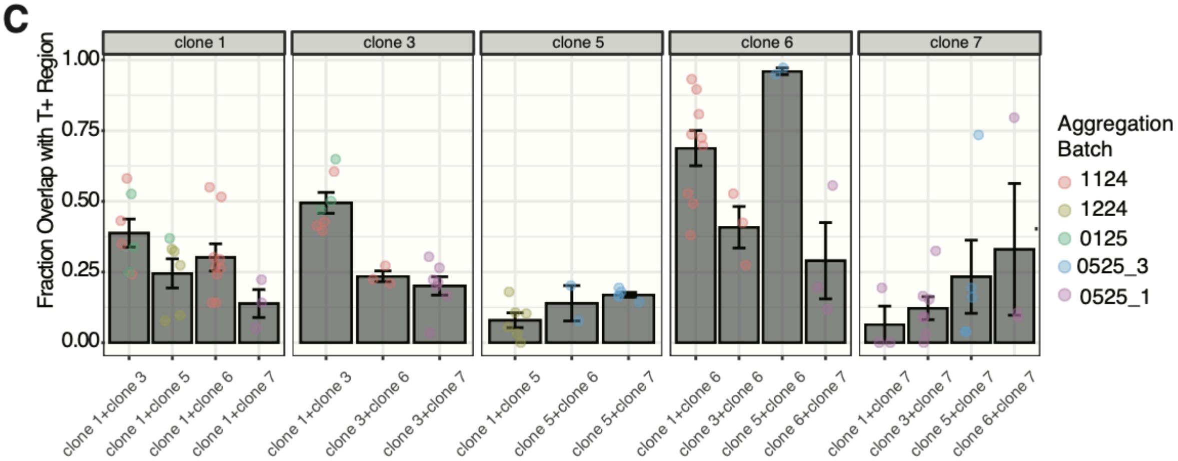


**Supplemental Figure 13. T expression in additional chimeric gastruloids.**
**(A)** Images corresponding to T expression in gastruloids formed from combinations of two clones.
**(B)** Bargraphs representing the fraction of a chimeric gastruloid's area that stains positively for T. Each point corresponds to a gastruloid, colored by the batch in which it was aggregated.
**(C)** Bar graphs depicting the overlap of each clone's spatial distribution with the distribution of T expression for chimeric gastruloids within a single batch of aggregation. Facets correspond to each individual clone within a combo, and each bar corresponds to that clone's behavior within a given clone combination. Each point

corresponds to a single gastruloid, and colors correspond to the batch in which each gastruloid was aggregated.

To determine whether clones occupy regions outside their usual spatial territories,  we immunostained gastruloids formed from different clone combinations (**Supplemental Figure 12C-E**, **Supplemental Figure 13**) and looked at how much of the area occupied by each clone in a pair expressed T. In most chimeric gastruloids, a small proportion (typically < 25%) of the anterior clone's area was positive for T. In some specific clone pairings, like clone 1+3 (**Supplemental Figure 12E**, **Supplemental Figure 13**) and clone 6+7 (**Supplemental Figure 13**), the anterior clone occupied the anterior region of the gastruloid, and also had cells residing in the T+ posterior tip of the gastruloid.

**Supplemental Figure 14**

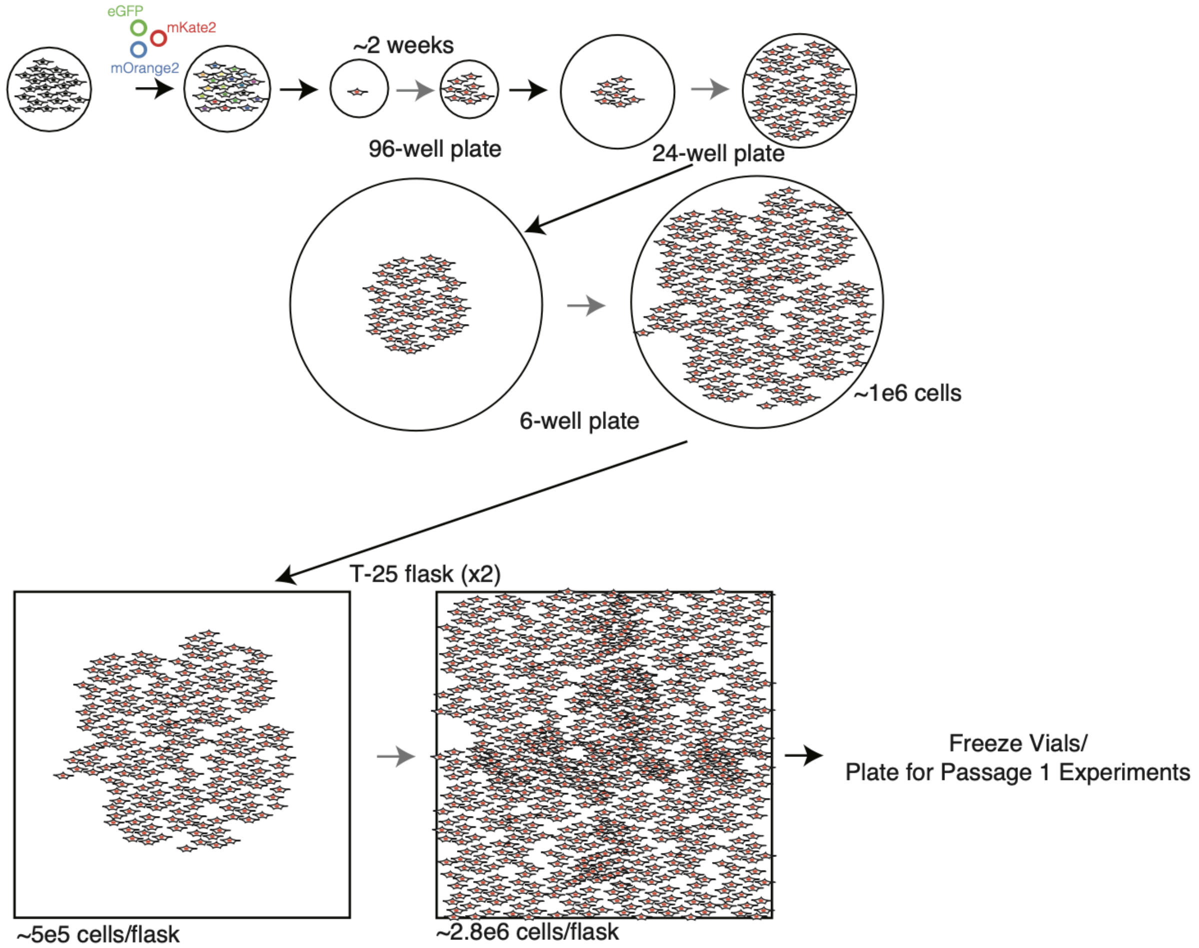


**Supplemental Figure 14. Expansion of mES cell clones.**
Schematic describing the workflow for isolating mES cell clones. Cell proliferation is labeled with a gray arrow, transfer of cells to larger plates/flasks is labeled with a black arrow.

A bulk population of E14-Tg2a mESCs was transduced at high multiplicity of infection (MOI) with lentiviral vectors encoding eGFP, mKate2, or mOrange2 to generate a multicolor, polyclonal population. This population was bottlenecked by single-cell dilution into 96-well plates (1 cell per well), and individual clones were expanded in 2i+LIF for ~2 weeks. Each clone then underwent multiple rounds of expansion: from 24-well plates to 6-well plates, and finally to T-25 flasks, where cell numbers reached ~2.8 million/flask. Cells were then frozen, and experiments were conducted on cells passaged 1–2 times after thawing (passage numbering starts at thawing, experiments are thus labeled in results as Passage 1, 2, etc.). Each clone therefore underwent many rounds of proliferation and expansion prior to experimentation, yet retained stable spatial propensity. Only with additional rounds of passaging beyond this expansion phase did clones begin to lose their positional bias. Thus, loss of propensity is not due to cumulative cell divisions per se, but rather serial passaging.

**Supplemental Figure 15**

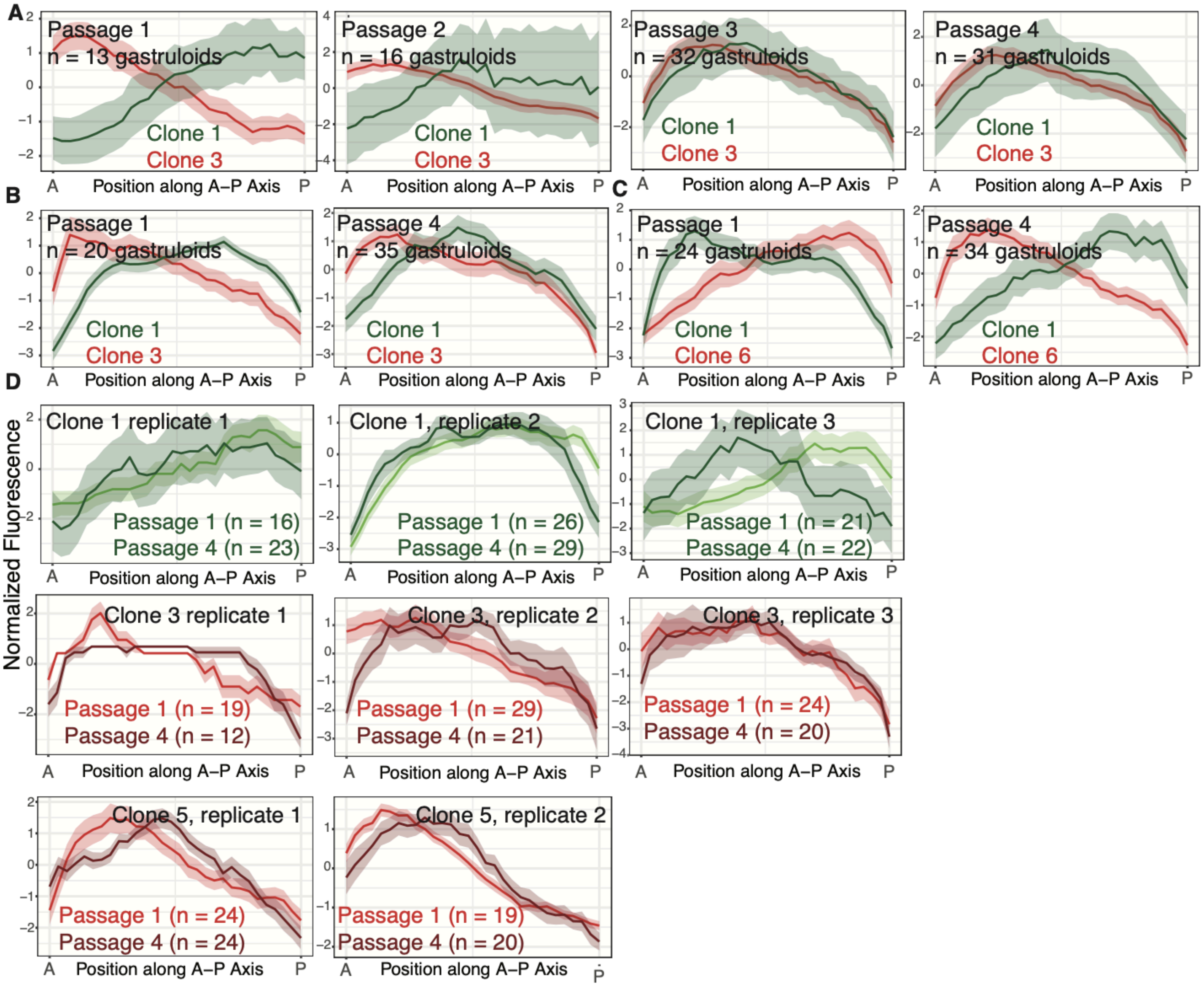


**Supplemental Figure 15. Additional replicate data comparing clones' propensity across passages.**
**(A)** Gastruloids generated from clone 1 and clone 3. Line-scans corresponding to passages 1 and 3 are shown in **Figure 3A**.
**(B)** Additional replicate data for this clone combination.
**(C)** Line-scans corresponding to fluorescence intensities along the A-P axis in red and green channels for gastruloids generated from a combination of clone 1 and clone 6.
**(D,** *first row*) Line-scans corresponding to the propensity of clone 1 at passages 1 and 4 in three independent experiments. Line scans correspond to the fluorescence intensity along the A-P axis in the green channel, for gastruloids generated from clone 1 and bulk mES cells. (*second row*) Line-scans corresponding to the propensity of clone 3 at passages 1 and 4 in three independent experiments. Line scans correspond to the fluorescence intensity along the A-P axis in the red channel, for gastruloids generated from clone 3 and bulk mES cells. (*third row*) Line-scans corresponding to the propensity of clone 5 at passages 1 and 4 in two independent experiments. Line scans correspond to the fluorescence intensity along the A-P axis in the red channel, for gastruloids generated from clone 5 and bulk mES cells.

In **Figure 3**, we showed that clones' propensity is not stable over repeated rounds of passaging. In most cases, by passage 3 or passage 4, chimeric gastruloids had a well-mixed arrangement of clones. However, there were some exceptions. For example, in one replicate, clone 3 appeared to have maintained its propensity at passage 4 (**Supplemental Figure 15D**). Moreover, in a chimeric gastruloid formed from clone 1 and clone 6 at passage 4, clone 6 assumed anterior positions (**Supplemental Figure 15C**). It could be that each clone loses propensity at a different rate, and so by passage 4, clone 6 has no obvious propensity but clone 1 still has residual posterior propensity, resulting in clone 1 occupying posterior positions in this combination.

**Supplemental Figure 16**

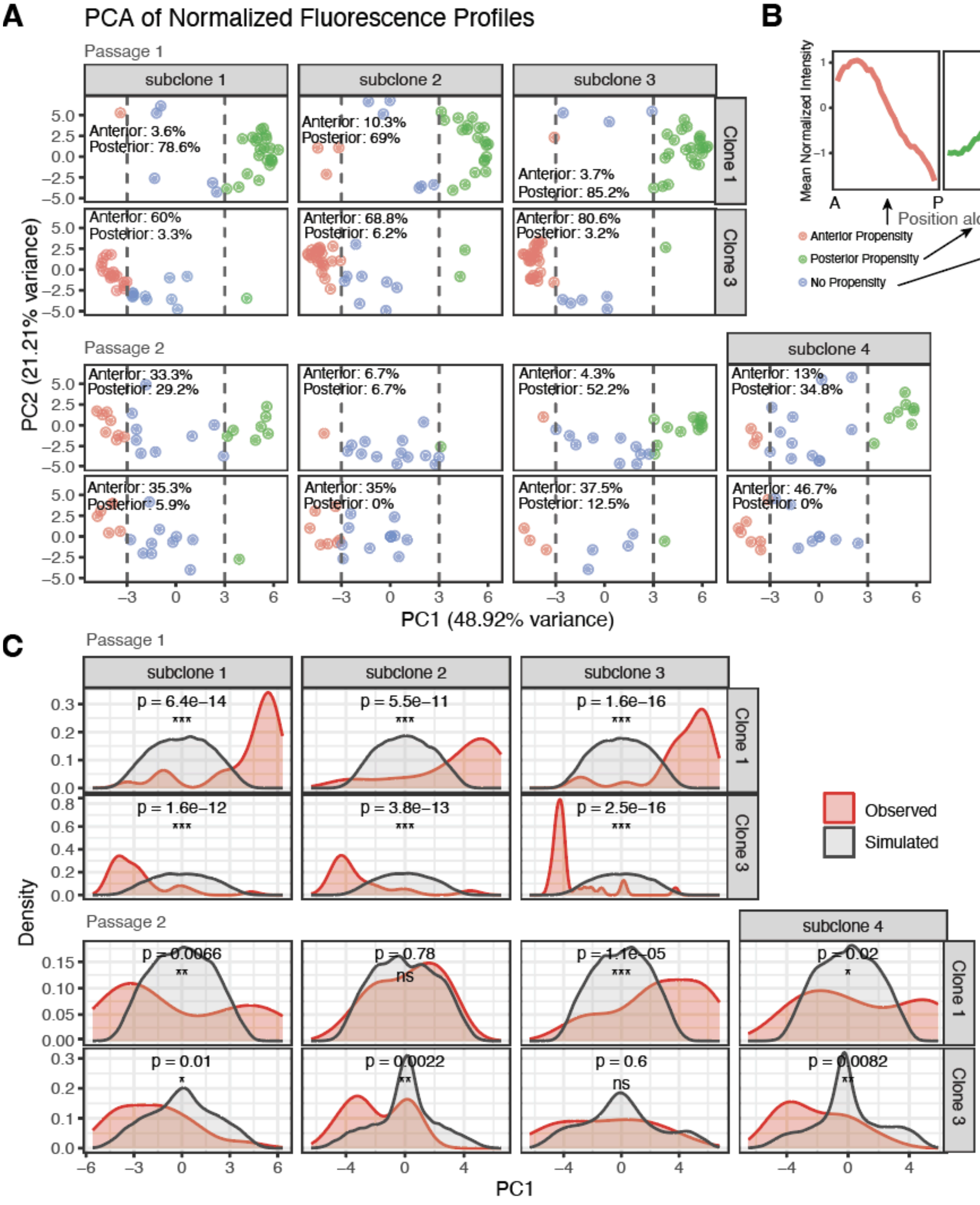


**Supplemental Figure 16. Additional evidence describing the propensities of subclones isolated before and after passaging.**

**(A)** Principal component analysis (PCA) of gastruloid fluorescence intensity profiles. Each point represents a gastruloid, with PC1 values < -3 considered to represent gastruloids with greater fluorescence intensity in the anterior ("Anterior Propensity") and PC1 values > 3 considered to represent gastruloids with greater fluorescence intensity in the posterior ("Posterior Propensity"). The proportions of gastruloids from each subclone that fall into each category are written.

**(B)** Linescans generated by averaging linescans from all gastruloids annotated as "Anterior Propensity", "Posterior Propensity", or "No Propensity", respectively.

**(C)** Kernel density estimates comparing observed and simulated distributions of PC1 values. The simulated/null distribution represents PC1 values for gastruloids containing labeled cells that have no propensity and could therefore fall anywhere on the A-P axis. P-values were computed using a K-S test for significance.

In **Figure 3**, we showed that clones' propensity is not stable over repeated rounds of passaging and that subclones isolated after passaging do not appear to retain the memory of their respective parental clones. We wanted to better characterize whether there might be mixed propensities represented within the expanded post-passage subclones. We first took the fluorescence intensity profiles along each gastruloid's A-P axis and performed principal component analysis (PCA). Gastruloids with labeled cells in their anterior were clearly separated in PCA space, along the first principal component (PC1) from gastruloids with labeled cells in their posterior (**Supplemental Figure 16A**). We set thresholds such that gastruloids with PC1 values less than -3 were considered to have labeled cells with "anterior propensity", and those with PC1 values greater than +3 were considered to have labeled cells with "posterior propensity." Those with PC1 values between -3 and 3 did not have propensity, which we could confirm by visualizing the average fluorescence intensity profiles of all gastruloids within each category (**Supplemental Figure 16B**). While the majority of gastruloids generated from each Passage 1 subclone had fluorescence intensity profiles consistent with the propensity of their parental clone, gastruloids generated from Passage 2 subclones had mixed propensities; some had fluorescence intensity profiles that were the direct opposite of what the propensity of their parental clone would predict, and others had no propensity with fluorescence evenly distributed along the A-P axis (**Supplemental Figure 16C**). To distinguish subclones with anterior or posterior propensity from subclones with no propensity, and to also distinguish subclones with mixed propensities (i.e., the expanded subclones had some subsets with anterior/posterior propensity), we took fluorescence intensity profiles of each gastruloid from each subclone, and scrambled the fluorescence intensities to generate a "random linescan," which could represent a gastruloid with no propensity (i.e., labeled cells could be anywhere on the A-P axis). We could then see where this random linescan fell in PCA space. After performing 1000 simulations, we could obtain a null distribution of PC1 values and compare the simulated distribution to the observed distribution of PC1 values for each subclone, performing a Kolmogorov-Smirnov (KS) test for significance. Some subclones isolated after passaging generated distributions that did not differ from the null distribution; we consider these subclones to truly lack propensity. However, others generated bimodal distributions, suggesting that within these expanded subclones, there are some cells with anterior propensity and others with posterior propensity.

**Supplemental Figure 17**

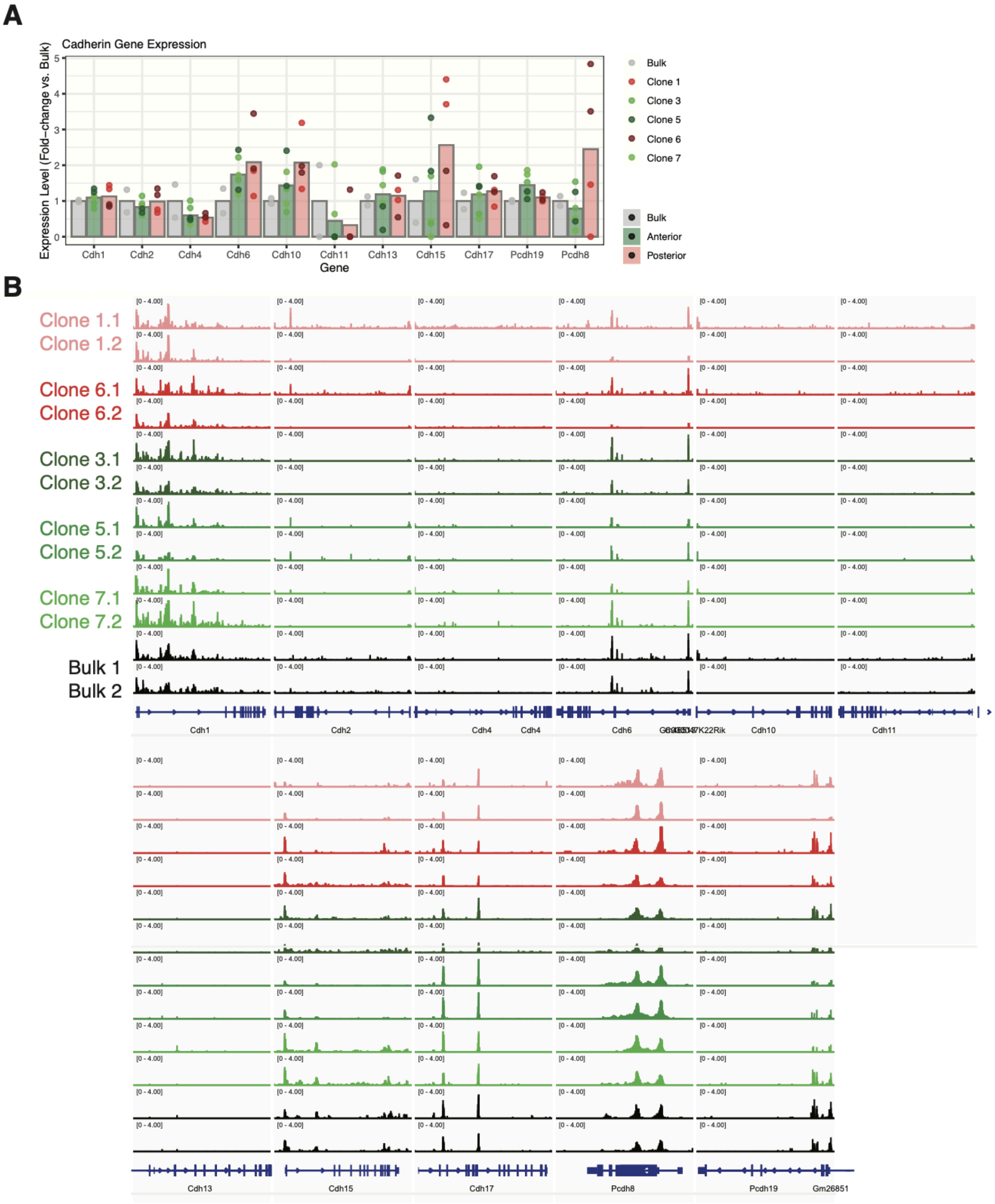


**Supplemental Figure 17. Comparison of cadherin gene expression and accessibility of genomic regions near cadherin genes.**

**(A)** Bar graphs comparing expression of *Cdh1, Cdh2, Cdh4, Cdh6, Cdh10, Cdh11, Cdh13, Cdh15, Cdh17, Pcdh19, and Pcdh8* among bulk mES cells, anterior clones, and posterior clones.

**(B)** ATAC-seq tracks comparing accessibility of genomic regions around *Cdh1*, *Cdh2, Cdh4, Cdh6, Cdh10, Cdh11, Cdh13, Cdh15, Cdh17, Pcdh8, and Pcdh19* among all clones and bulk.

**Supplemental Figure 18**

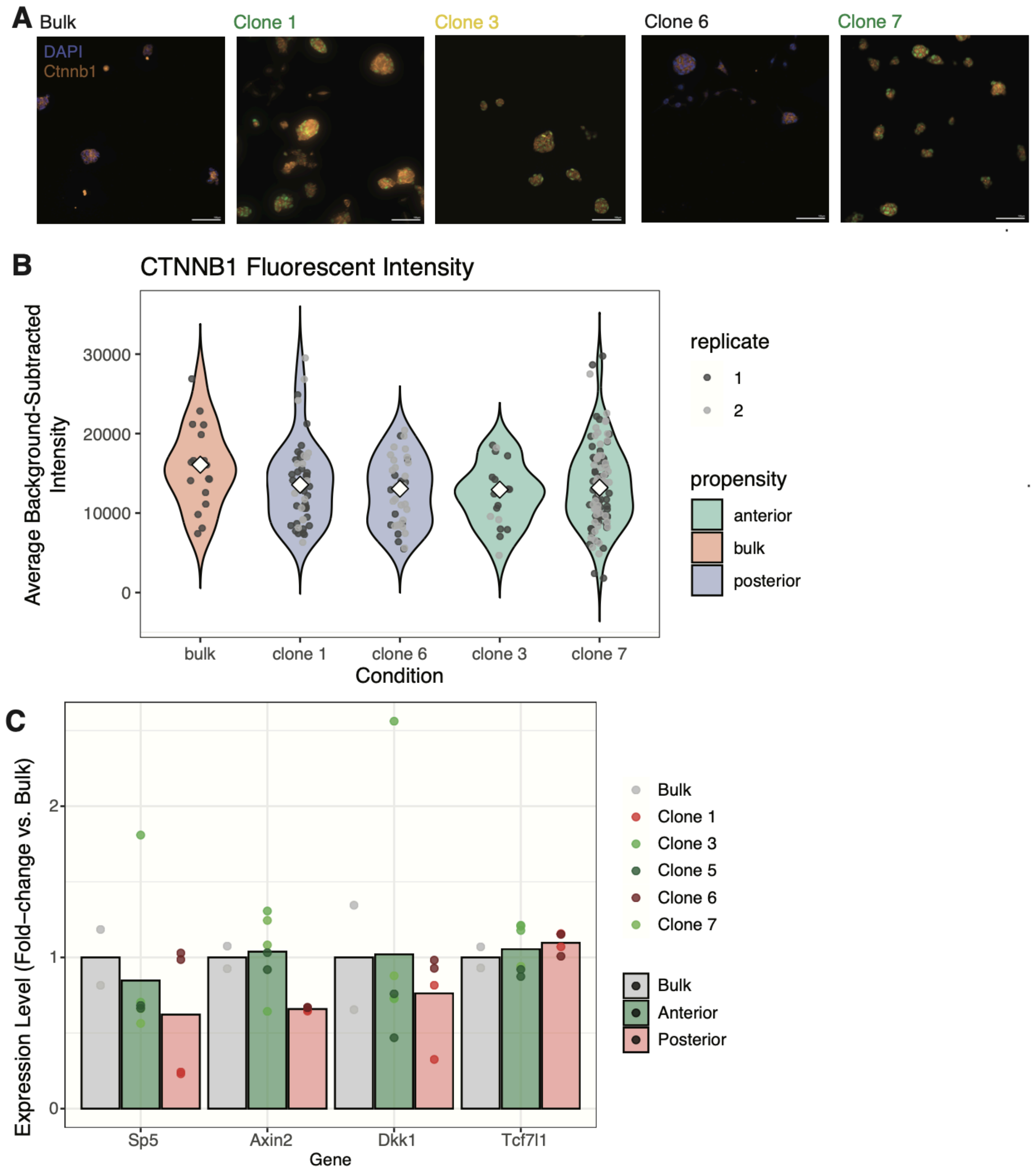


**Supplemental Figure 18. Beta-catenin expression in mES cell clones.**
**(A)** Representative images showing beta-catenin expression in bulk mES cells and posterior clones 1 and 6 and anterior clones 3 and 7.
**(B)** Comparison of immunofluorescence staining intensities across mES cell colonies from bulk and each clone. Each point corresponds to a colony, colored by technical replicate. Violins are colored by propensity.

One question is whether mES cell clones differ in their baseline Wnt pathway activity, even under 2i conditions. We therefore plated cells sparsely, and after 48 hours of growth (to make it easier to segment individual colonies), we immunostained for beta-catenin and imaged. Each colony was manually annotated, and the average fluorescence staining intensity was computed for all z-planes. For each annotation, we then selected the z-slice with the highest average staining intensity for each colony and subtracted background by finding the average intensity of a 10px annulus around the annotation. We did not find differences in beta-catenin among clones and bulk.

**Supplemental Figure 19**

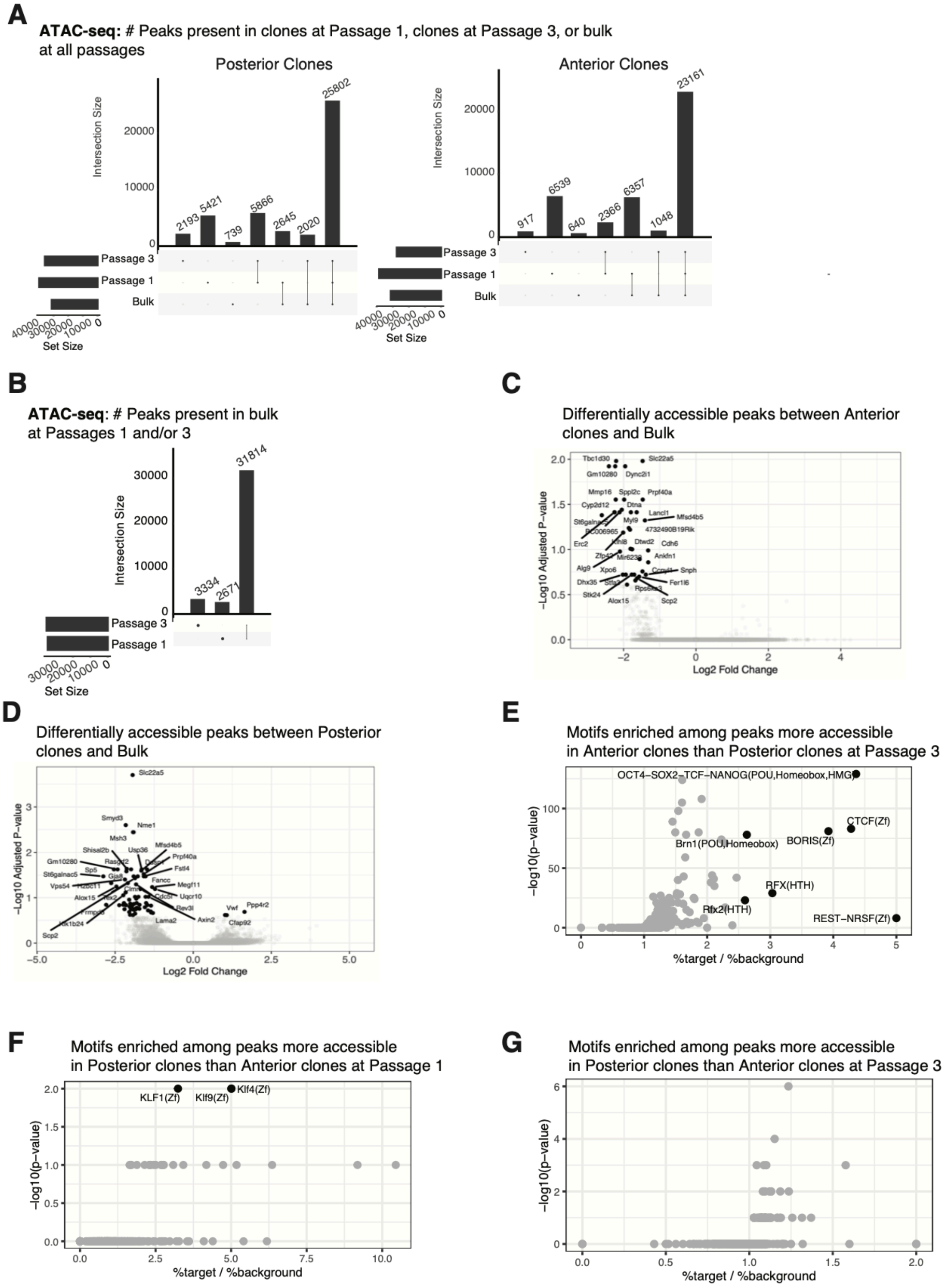


**Supplemental Figure 19. Additional data for differential accessibility and motif enrichment among clones.**

**(A)** Upset plots comparing unique and shared ATAC-seq peaks among bulk, clone samples harvested at passage 1, and clone samples harvested at passage 3. Separate plots were drawn for posterior clones and for anterior clones.

**(B)** Peaks unique to bulk samples harvested at passage 1 or passage 3, and shared between passage 1 and passage 3.
**(C)** Peaks differentially expressed between anterior clones and bulk.
**(D)** Peaks differentially expressed between posterior clones and bulk.
**(E)** Motifs enriched among peaks more accessible in anterior clones than posterior clones at passage 3.
(**F**) Motifs enriched among peaks more accessible in posterior clones than anterior clones at passage 1.
(**G**) Motifs enriched among peaks more accessible in posterior clones than anterior clones at passage 3.

#### Supplemental Figure 20

**A**

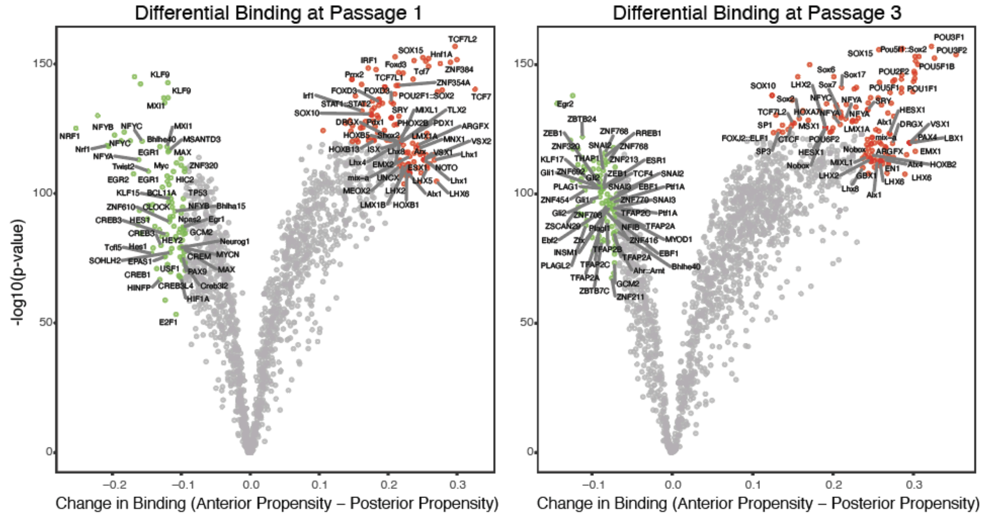


**B**

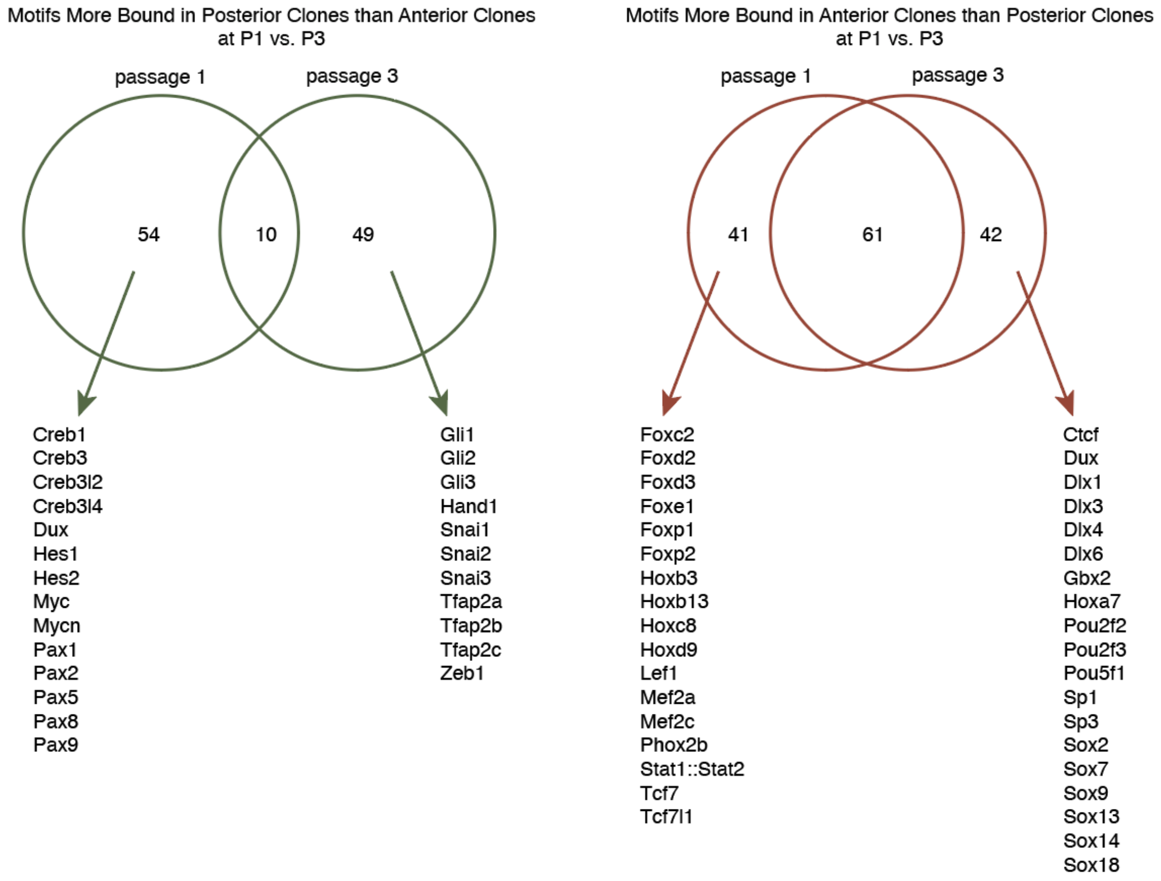


**Supplemental Figure 20. Transcription factor footprinting analysis using TOBIAS.**
**(A)** Volcano plots showing differentially-bound motifs between clones with an anterior propensity and those with a posterior propensity at passage 1 and at passage 3.
**(B)** Venn diagrams showing unique motifs more bound in anterior clones or posterior clones at passage 1 and at passage 3.

#### Supplemental Figure 21

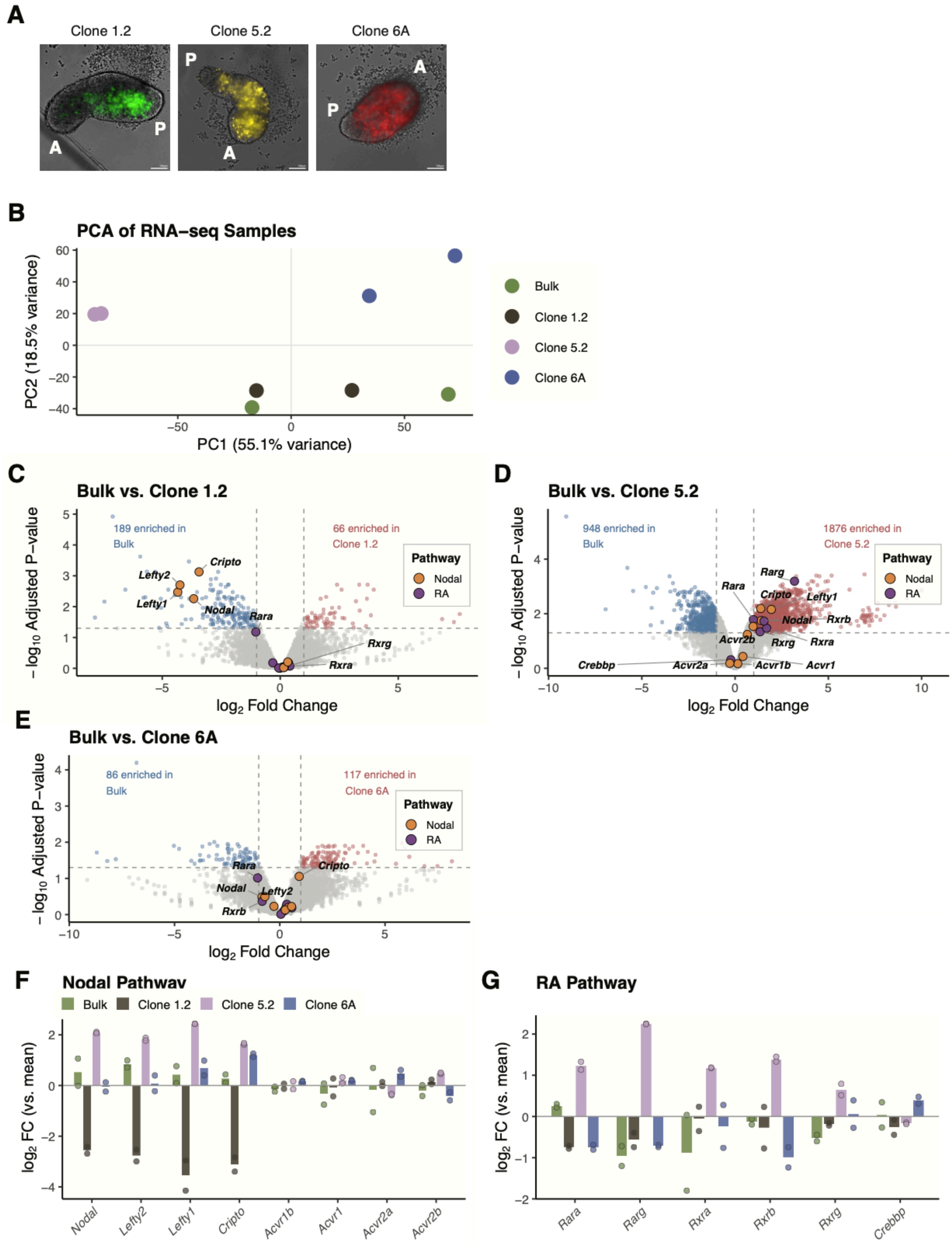

**Supplemental Figure 21. Bulk RNA-seq of 48h gastruloids. (A)** Representative images of chimeric gastruloids formed from a combination of unlabeled bulk mES cells (50%) and fluorescently-labeled clones (50%). **(B)** PCA of RNA-seq samples. **(C)** Volcano plots showing differentially-expressed genes between bulk mES cells and each clone that was sequenced. **(D)** Bargraphs showing differences in gene expression of genes associated with Nodal and RA signaling among clones.

**Supplemental Figure 22**

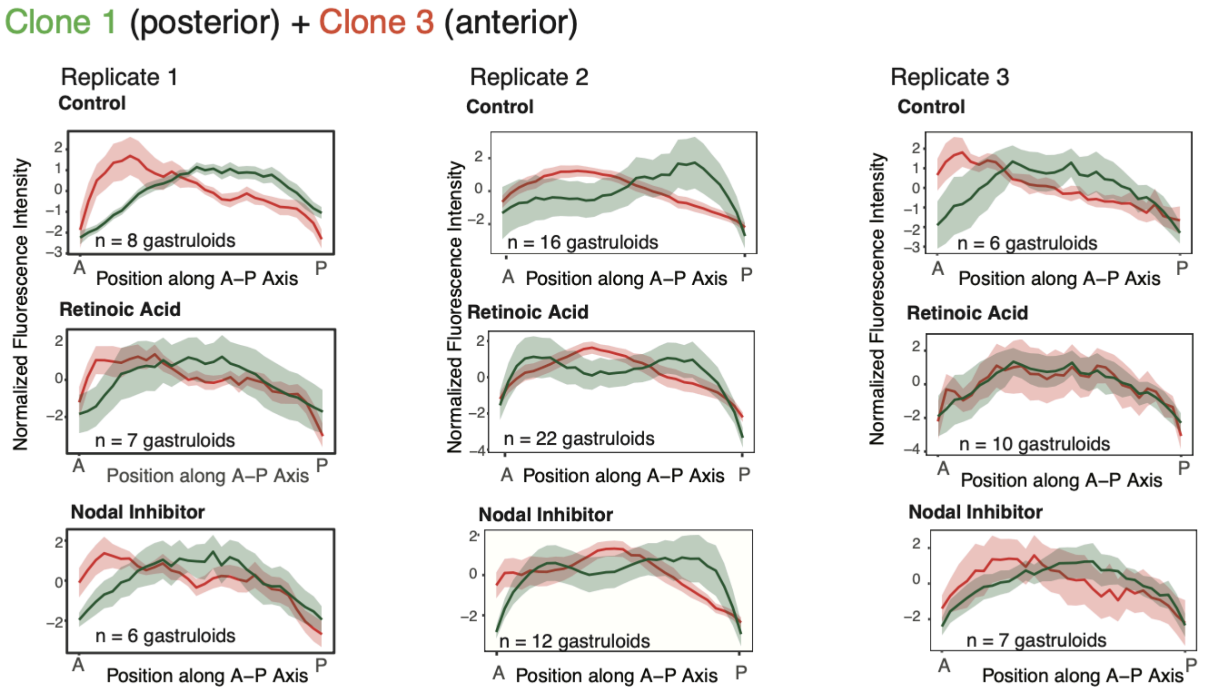


**Supplemental Figure 22. Additional data showing the effect of retinoic acid treatment and Nodal inhibition on propensity.**

Fluorescence intensity in red and green channels along the A-P axis of gastruloids under control conditions, under retinoic acid treatment, and under Nodal inhibition with SB-431542, similar to **Figure 5C**. Data here correspond to the second and third independent experiments for gastruloids generated from the combination of clone 1 and clone 3.

**Supplemental Figure 23**

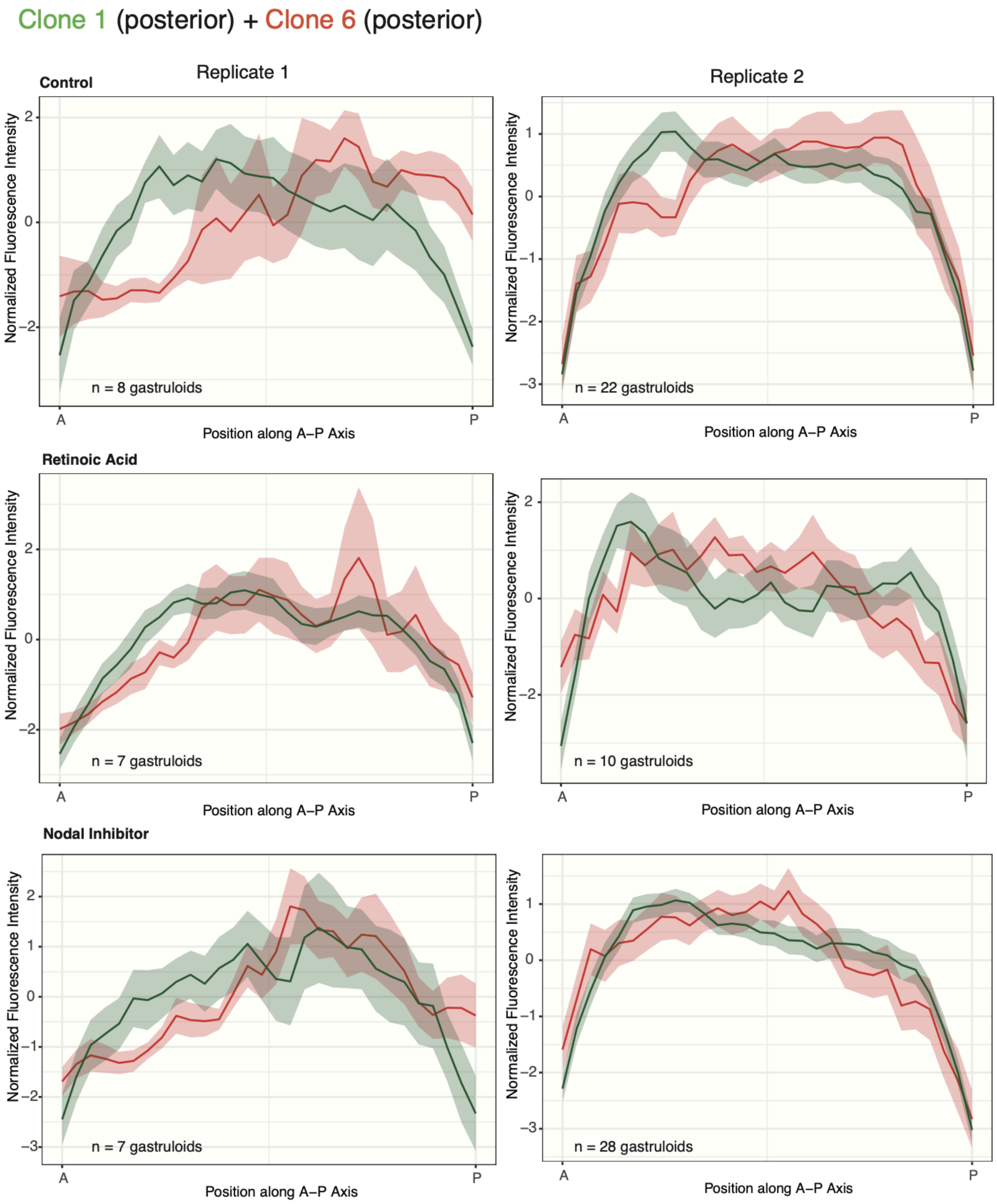


**Supplemental Figure 23. Additional data showing the effect of retinoic acid treatment and Nodal inhibition on propensity in an additional clone combination.**
Fluorescence intensity in red and green channels along the A-P axis of gastruloids under control conditions, under retinoic acid treatment, and under Nodal inhibition with SB-431542, similar to **Figure 5C**. Data here correspond to two independent experiments for gastruloids generated from the combination of clone 1 and clone 6.

**Supplemental Figure 24**

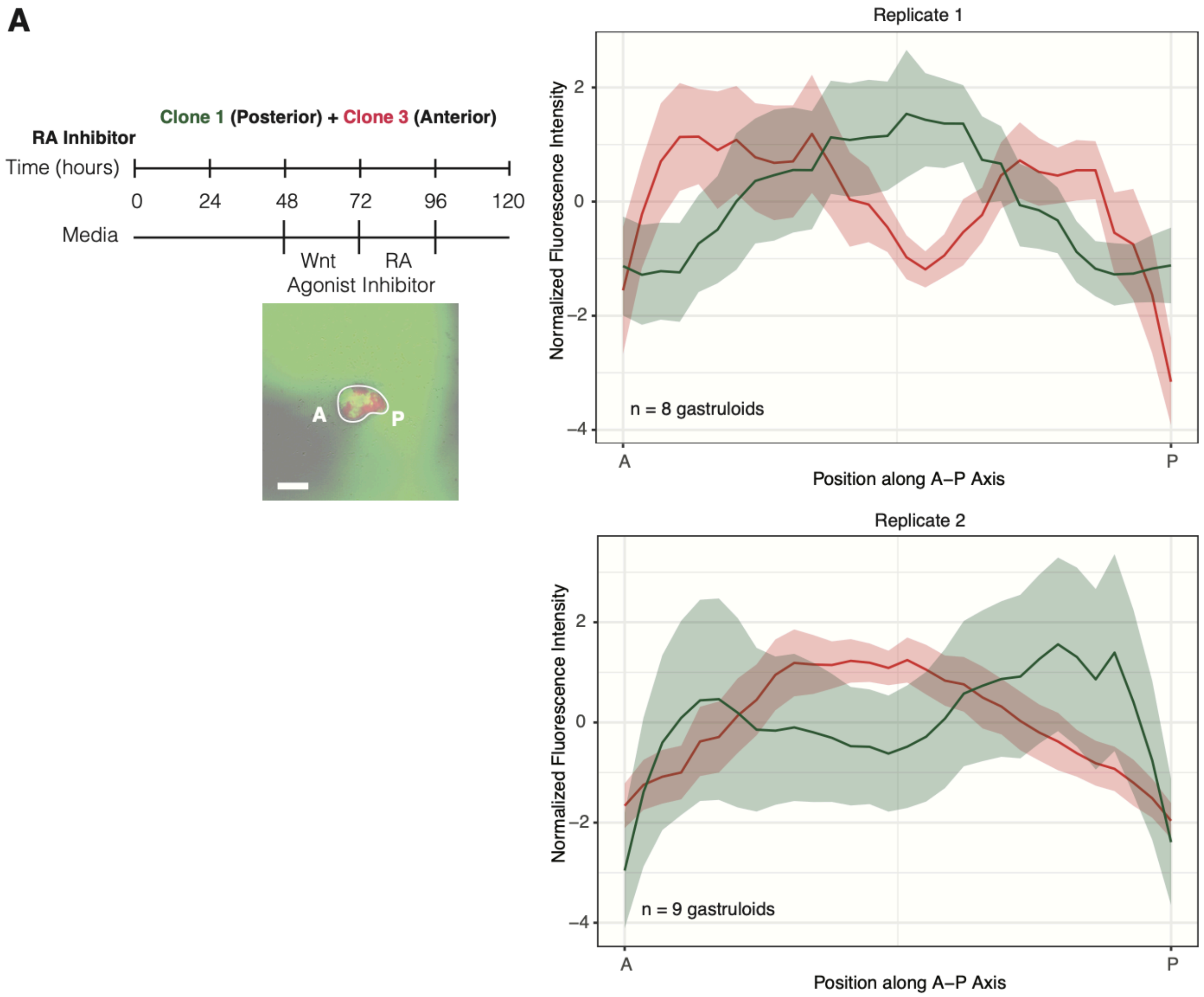


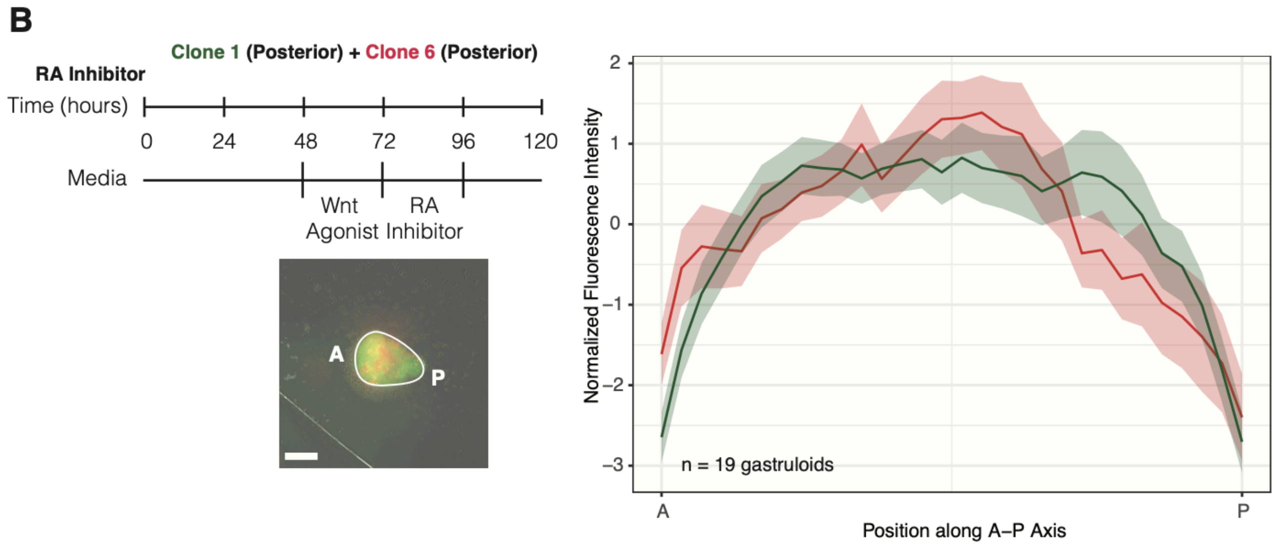


**Supplemental Figure 24. Effect of RA inhibition on clones' propensity.**
Gastruloids made from the combination of **(A)** clone 1 and clone 3 or **(B)** clone 1 and clone 6 were treated with 100nM AGN-193109 from 96-120 hours after aggregation. Experiments were performed alongside those

displayed in **Figure 5C** (for panel **A**, replicate 1); **Supplemental Figure 22** (for panel **A**, replicate 2); and **Supplemental Figure 23** (for panel **B**)

**Supplemental Figure 25**

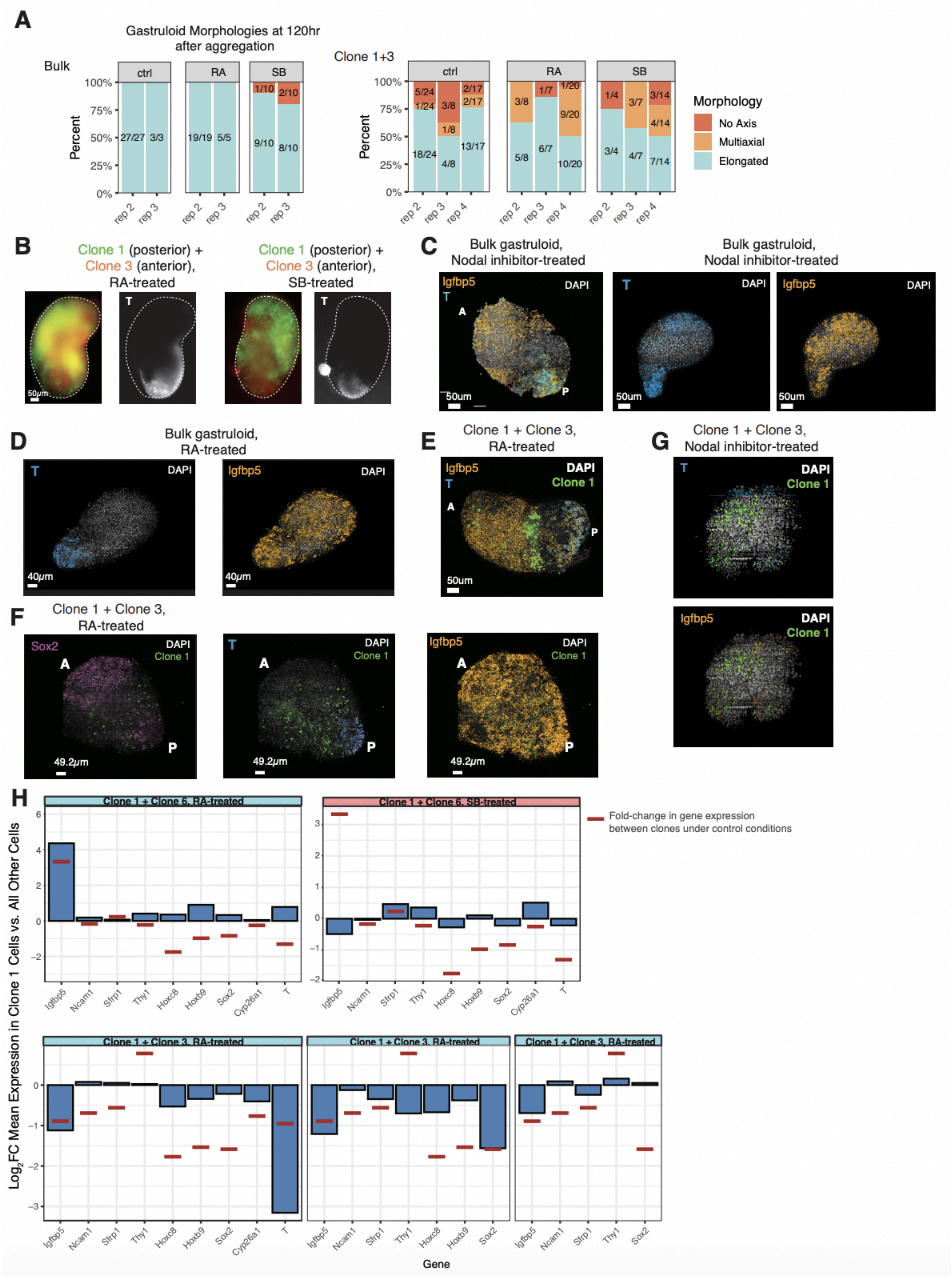


**Supplemental Figure 25. Spatial transcriptomics of RA treatment and Nodal inhibition.**

**(A)** Additional replicate data showing the fraction of gastruloids that elongate following treatment with retinoic acid (RA) or Nodal inhibitor (SB).
**(B)** Representative immunofluorescence images showing expression of T in RA-treated and Nodal-inhibited (SB) clone 1+3 gastruloids
**(C)** Representative images of SB-treated bulk gastruloids. The image shows spatial expression of *Igfbp5* and *T*.
**(D)** Representative images of an RA-treated bulk gastruloids. The image shows spatial expression of *Igfbp5* and *T*.
**(E)** Representative image of an RA-treated gastruloid comprising clone 1 and clone 3. The image shows spatial expression of *Igfbp5* and *T*.
**(F)** Representative image of an RA-treated gastruloid comprising clone 1 and clone 3. The image shows spatial expression of *Sox2* as well as of *T*, and of *Igfbp5.*
**(G)** Representative image of an SB-treated gastruloid comprising clone 1 and clone 3. The image shows spatial expression of *Igfbp5* and *T*.
**(H)** Bar graphs showing the fold-change in gene expression between cells segmented as belonging to clone 1 versus all other cells in chemically-treated (RA or Nodal-inhibitor) chimeric gastruloids comprising clone 1 and clone 3 or clone 1 and clone 6. As described in **Methods**, for rounds of hybridization in which focus issues prevented spot counting, affected genes were not included in this analysis. For one RA-treated gastruloid, *Cyp26a1* and *T* were removed from analysis; for another, *Cyp26a1*, *Hoxc8*, and *Hoxb9* were removed. Red lines near each bar correspond to the fold change in gene expression between clone 1 and other cells in control gastruloids.

Earlier, in **Supplemental Figure 11**, we observed differences in gene expression between clone 1 and clone 6 or clone 1 and clone 3 within chimeric gastruloids. We were curious how RA treatment or Nodal inhibition would affect gene expression differences between clones in chimeric gastruloids. We specifically focused on markers that are markers for spatial regions of gastruloids, like anterior markers *Igfbp5*, *Ncam1*, and *Sfrp1,* and posterior markers (*Hoxc8*, *Hoxb9*, *Sox2*, *Cyp26a1*, and *T*; notably, these are also markers for neuromesodermal progenitors). We also included *Thy1* in this analysis, since its expression was higher in clone 1 cells within clone 1+3 gastruloids under control conditions. For most of the genes we analyzed, RA treatment and Nodal inhibition decreased relative gene expression differences between clones (**Supplemental Figure 25B**). However, RA treatment actually made differences in *Igfbp5* expression more extreme in clone 1+3 and clone 1+6 gastruloids and made differences in *T* more extreme in clone 1+3 gastruloids. Increases in differential *Igfbp5* and *T* expression may reflect a global reduction in expression of posterior markers like *T* and more widespread expression of *Igfbp5* under RA treatment (**Figure 5H**).

#### Supplemental Figure 26

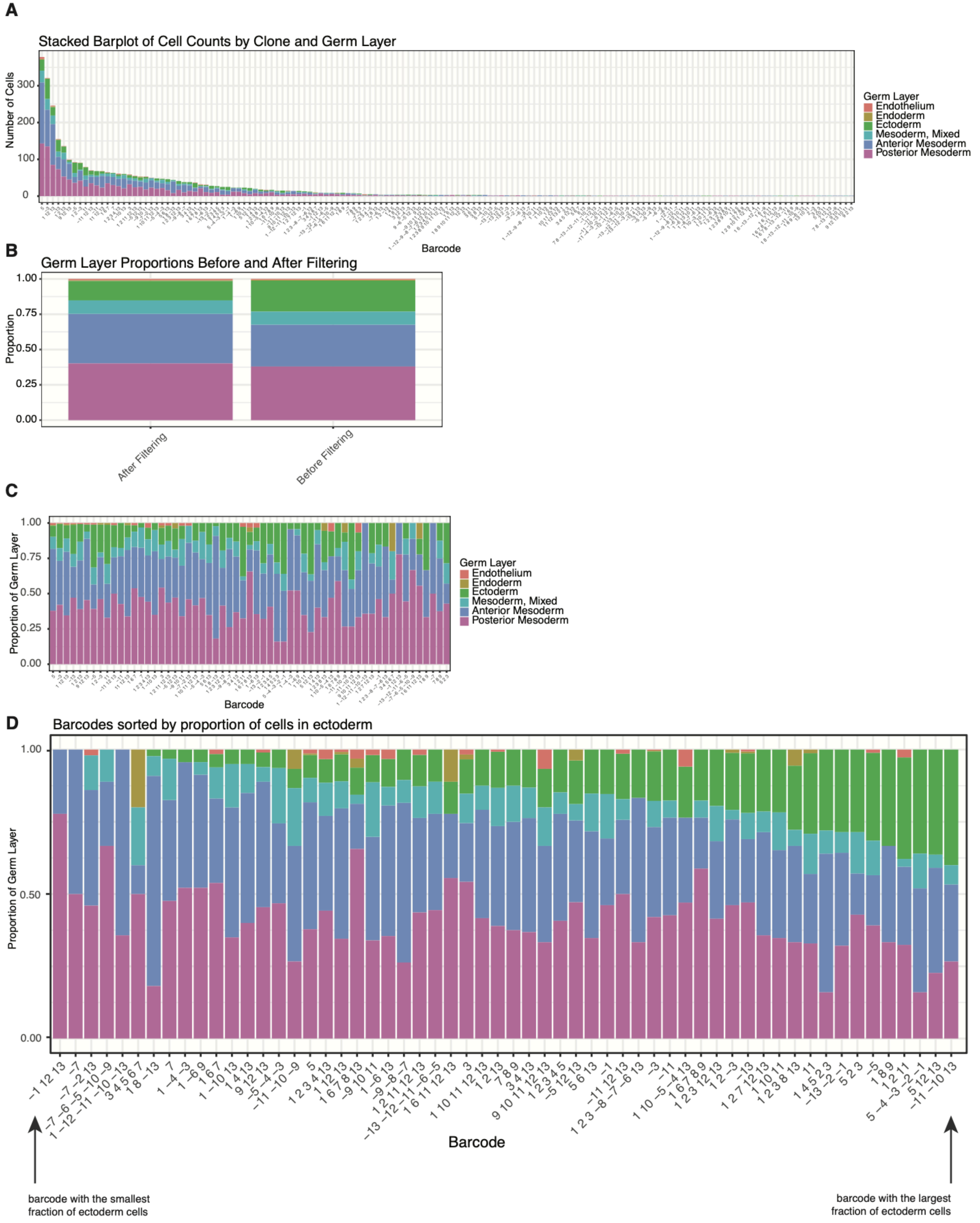

**Supplemental Figure 26. Analysis of Weber et al.**
**(A)** Stacked barplot of cell counts by clone barcode, colored by germ layer.
**(B)** Proportions of cells within the dataset annotated as belonging to each germ layer before and after filtering barcodes from the exclusion list provided by Weber et al. and those with low cell number.
**(C)** Normalized stacked bar graphs, showing proportions of cells belonging to each germ layer for clones used in the final analysis.
**(D)** Clones used in the final analysis, ordered left-to-right by increasing proportion of ectoderm cells.

**Supplemental Movie 1.**
Elongating gastruloid generated from a polyclonal mES cell population. Colors correspond to clonal lineages of mES cells.

**Supplemental Movie 2.**
Gastruloid generated from bulk mES cells and a clone with anterior propensity (50%/50% chimera).

**Supplemental Table 1. Primers used to index ATAC sequencing libraries.**

| Primer Name | Index | Sequence |
|---|---|---|
| N5001 | TAGATCGC | AATGATACGGCGACCACCGAGATCTACACTAGATCGCTCGTCGGCAGCGTCAGATGTGTAT |
| N5002 | CTCTCTAT | AATGATACGGCGACCACCGAGATCTACACCTCTCTATTCGTCGGCAGCGTCAGATGTGTAT |
| N5003 | TATCCTCT | AATGATACGGCGACCACCGAGATCTACACTATCCTCTTCGTCGGCAGCGTCAGATGTGTAT |
| N5004 | AGAGTAGA | AATGATACGGCGACCACCGAGATCTACACAGAGTAGATCGTCGGCAGCGTCAGATGTGTAT |
| N5005 | GTAAGGAG | AATGATACGGCGACCACCGAGATCTACACGTAAGGAGTCGTCGGCAGCGTCAGATGTGTAT |
| N5006 | ACTGCATA | AATGATACGGCGACCACCGAGATCTACACACTGCATATCGTCGGCAGCGTCAGATGTGTAT |
| N5007 | AAGGAGTA | AATGATACGGCGACCACCGAGATCTACACAAGGAGTATCGTCGGCAGCGTCAGATGTGTAT |
| N5008 | CTAAGCCT | AATGATACGGCGACCACCGAGATCTACACCTAAGCCTTCGTCGGCAGCGTCAGATGTGTAT |
| N5009 | TGGAAATC | AATGATACGGCGACCACCGAGATCTACACTGGAAATCTCGTCGGCAGCGTCAGATGTGTAT |
| N5010 | AACATGAT | AATGATACGGCGACCACCGAGATCTACACAACATGATTCGTCGGCAGCGTCAGATGTGTAT |
| N5011 | TGATGAAA | AATGATACGGCGACCACCGAGATCTACACTGATGAAATCGTCGGCAGCGTCAGATGTGTAT |
| N5012 | GTCGGACT | AATGATACGGCGACCACCGAGATCTACACGTCGGACTTCGTCGGCAGCGTCAGATGTGTAT |
| N7001 | TAAGGCGA | CAAGCAGAAGACGGCATACGAGATTCGCCTTAGTCTCGTGGGCTCGGAGATGTG |
| N7002 | CGTACTAG | CAAGCAGAAGACGGCATACGAGATCTAGTACGGTCTCGTGGGCTCGGAGATGTG |
| N7003 | AGGCAGAA | CAAGCAGAAGACGGCATACGAGATTTCTGCCTGTCTCGTGGGCTCGGAGATGTG |
| N7004 | TCCTGAGC | CAAGCAGAAGACGGCATACGAGATGCTCAGGAGTCTCGTGGGCTCGGAGATGTG |
| N7005 | GGACTCCT | CAAGCAGAAGACGGCATACGAGATAGGAGTCCGTCTCGTGGGCTCGGAGATGTG |
| N7006 | TAGGCATG | CAAGCAGAAGACGGCATACGAGATCATGCCTAGTCTCGTGGGCTCGGAGATGTG |
| N7007 | CTCTCTAC | CAAGCAGAAGACGGCATACGAGATGTAGAGAGGTCTCGTGGGCTCGGAGATGTG |
| N7008 | CAGAGAGG | CAAGCAGAAGACGGCATACGAGATCCTCTCTGGTCTCGTGGGCTCGGAGATGTG |
| N7009 | GCTACGCT | CAAGCAGAAGACGGCATACGAGATAGCGTAGCGTCTCGTGGGCTCGGAGATGTG |
| N7010 | CGAGGCTG | CAAGCAGAAGACGGCATACGAGATCAGCCTCGGTCTCGTGGGCTCGGAGATGTG |
| N7011 | AAGAGGCA | CAAGCAGAAGACGGCATACGAGATTGCCTCTTGTCTCGTGGGCTCGGAGATGTG |
| N7012 | GTAGAGGA | CAAGCAGAAGACGGCATACGAGATTCCTCTACGTCTCGTGGGCTCGGAGATGTG |
